# Opportunities and obstacles for deep learning in biology and medicine

**DOI:** 10.1101/142760

**Authors:** Travers Ching, Daniel S. Himmelstein, Brett K. Beaulieu-Jones, Alexandr A. Kalinin, Brian T. Do, Gregory P. Way, Enrico Ferrero, Paul-Michael Agapow, Michael Zietz, Michael M. Hoffman, Wei Xie, Gail L. Rosen, Benjamin J. Lengerich, Johnny Israeli, Jack Lanchantin, Stephen Woloszynek, Anne E. Carpenter, Avanti Shrikumar, Jinbo Xu, Evan M. Cofer, Christopher A. Lavender, Srinivas C. Turaga, Amr M. Alexandari, Zhiyong Lu, David J. Harris, Dave DeCaprio, Yanjun Qi, Anshul Kundaje, Yifan Peng, Laura K. Wiley, Marwin H.S. Segler, Simina M. Boca, S. Joshua Swamidass, Austin Huang, Anthony Gitter, Casey S. Greene

**Affiliations:** Molecular Biosciences and Bioengineering Graduate Program, University of Hawaii at Manoa, Honolulu, HI; Department of Systems Pharmacology and Translational Therapeutics, Perelman School of Medicine, University of Pennsylvania, Philadelphia, PA; Genomics and Computational Biology Graduate Group, Perelman School of Medicine, University of Pennsylvania, Philadelphia, PA; Department of Computational Medicine and Bioinformatics, University of Michigan Medical School, Ann Arbor, MI; Harvard Medical School, Boston, MA; Computational Biology and Stats, Target Sciences, GlaxoSmithKline, Stevenage, United Kingdom; Data Science Institute, Imperial College London, London, United Kingdom; Princess Margaret Cancer Centre, Toronto, ON, Canada; Department of Medical Biophysics, Toronto, ON, Canada; Department of Computer Science, Toronto, ON, Canada; Electrical Engineering and Computer Science, Vanderbilt University, Nashville, TN; Ecological and Evolutionary Signal-processing and Informatics Laboratory, Department of Electrical and Computer Engineering, Drexel University, Philadelphia, PA; Computational Biology Department, School of Computer Science, Carnegie Mellon University, Pittsburgh, PA; Biophysics Program, Stanford University, Stanford, CA; Department of Computer Science, University of Virginia, Charlottesville, VA; Imaging Platform, Broad Institute of Harvard and MIT, Cambridge, MA; Department of Computer Science, Stanford University, Stanford, CA; Toyota Technological Institute at Chicago, Chicago, IL; Department of Computer Science, Trinity University, San Antonio, TX; Lewis-Sigler Institute for Integrative Genomics, Princeton University, Princeton, NJ; Integrative Bioinformatics, National Institute of Environmental Health Sciences, National Institutes of Health, Research, Triangle Park, NC; Howard Hughes Medical Institute, Janelia Research Campus, Ashburn, VA; National Center for Biotechnology Information and National Library of Medicine, National Institutes of Health, Bethesda, MD; Department of Wildlife Ecology and Conservation, University of Florida, Gainesville, FL; ClosedLoop.ai, Austin, TX; Department of Genetics, Stanford University, Stanford, CA; Division of Biomedical Informatics and Personalized Medicine, University of Colorado School of Medicine, Aurora, CO; Institute of Organic Chemistry, Westfälische Wilhelms-Universität Münster, Münster, Germany; Innovation Center for Biomedical Informatics, Georgetown University Medical Center, Washington, DC; Department of Pathology and Immunology, Washington University in Saint Louis, Saint Louis, MO; Department of Medicine, Brown University, Providence, RI; Department of Biostatistics and Medical Informatics, University of Wisconsin-Madison, Madison, WI; Morgridge Institute for Research, Madison, WI

## Abstract

Deep learning, which describes a class of machine learning algorithms, has recently showed impressive results across a variety of domains. Biology and medicine are data rich, but the data are complex and often ill-understood. Problems of this nature may be particularly well-suited to deep learning techniques. We examine applications of deep learning to a variety of biomedical problems—patient classification, fundamental biological processes, and treatment of patients—and discuss whether deep learning will transform these tasks or if the biomedical sphere poses unique challenges. We find that deep learning has yet to revolutionize or definitively resolve any of these problems, but promising advances have been made on the prior state of the art. Even when improvement over a previous baseline has been modest, we have seen signs that deep learning methods may speed or aid human investigation. More work is needed to address concerns related to interpretability and how to best model each problem. Furthermore, the limited amount of labeled data for training presents problems in some domains, as do legal and privacy constraints on work with sensitive health records. Nonetheless, we foresee deep learning powering changes at both bench and bedside with the potential to transform several areas of biology and medicine.

## Introduction to deep learning

Biology and medicine are rapidly becoming data-intensive. A recent comparison of genomics with social media, online videos, and other data-intensive disciplines suggests that genomics alone will equal or surpass other fields in data generation and analysis within the next decade [1]. The volume and complexity of these data present new opportunities, but also pose new challenges. Automated algorithms that extract meaningful patterns could lead to actionable knowledge and change how we develop treatments, categorize patients, or study diseases, all within privacy-critical environments.

The term *deep learning* has come to refer to a collection of new techniques that, together, have demonstrated breakthrough gains over existing best-in-class machine learning algorithms across several fields. For example, over the past five years these methods have revolutionized image classification and speech recognition due to their flexibility and high accuracy [2]. More recently, deep learning algorithms have shown promise in fields as diverse as high-energy physics [3], dermatology [4], and translation among written languages [5]. Across fields, “off-the-shelf” implementations of these algorithms have produced comparable or higher accuracy than previous best-in-class methods that required years of extensive customization, and specialized implementations are now being used at industrial scales.

Deep learning approaches grew from research in neural networks, which were first proposed in 1943 [6] as a model for how our brains process information. The history of neural networks is interesting in its own right [7]. In neural networks, inputs are fed into the input layer, which feeds into one or more hidden layers, which eventually link to an output layer. A layer consists of a set of nodes, sometimes called “features” or “units,” which are connected via edges to the immediately earlier and the immediately deeper layers. In some special neural network architectures, nodes can connect to themselves with a delay. The nodes of the input layer generally consist of the variables being measured in the dataset of interest—for example, each node could represent the intensity value of a specific pixel in an image or the expression level of a gene in a specific transcriptomic experiment. The neural networks used for deep learning have multiple hidden layers. Each layer essentially performs feature construction for the layers before it. The training process used often allows layers deeper in the network to contribute to the refinement of earlier layers. For this reason, these algorithms can automatically engineer features that are suitable for many tasks and customize those features for one or more specific tasks.

Deep learning does many of the same things as more familiar machine learning approaches. In particular, deep learning approaches can be used both in *supervised* applications—where the goal is to accurately predict one or more labels or outcomes associated with each data point—in the place of regression approaches, as well as in *unsupervised*, or “exploratory” applications—where the goal is to summarize, explain, or identify interesting patterns in a data set—as a form of clustering. Deep learning methods may in fact combine both of these steps. When sufficient data are available and labeled, these methods construct features tuned to a specific problem and combine those features into a predictor. In fact, if the dataset is “labeled” with binary classes, a simple neural network with no hidden layers and no cycles between units is equivalent to logistic regression if the output layer is a sigmoid (logistic) function of the input layer. Similarly, for continuous outcomes, linear regression can be seen as a simple neural network. Thus, in some ways, supervised deep learning approaches can be seen as a generalization of regression models that allow for greater flexibility. Recently, hardware improvements and very large training datasets have allowed these deep learning techniques to surpass other machine learning algorithms for many problems. In a famous and early example, scientists from Google demonstrated that a neural network “discovered” that cats, faces, and pedestrians were important components of online videos [8] without being told to look for them. What if, more generally, deep learning could solve the challenges presented by the growth of data in biomedicine? Could these algorithms identify the “cats” hidden in our data—the patterns unknown to the researcher—and suggest ways to act on them? In this review, we examine deep learning’s application to biomedical science and discuss the unique challenges that biomedical data pose for deep learning methods.

Several important advances make the current surge of work done in this area possible. Easy-to-use software packages have brought the techniques of the field out of the specialist’s toolkit to a broad community of computational scientists. Additionally, new techniques for fast training have enabled their application to larger datasets [9]. Dropout of nodes, edges, and layers makes networks more robust, even when the number of parameters is very large. Finally, the larger datasets now available are also sufficient for fitting the many parameters that exist for deep neural networks. The convergence of these factors currently makes deep learning extremely adaptable and capable of addressing the nuanced differences of each domain to which it is applied.

This review discusses recent work in the biomedical domain, and most successful applications select neural network architectures that are well suited to the problem at hand. We sketch out a few simple example architectures in Figure 1. If data have a natural adjacency structure, a convolutional neural network (CNN) can take advantage of that structure by emphasizing local relationships, especially when convolutional layers are used in early layers of the neural network. Other neural network architectures such as autoencoders require no labels and are now regularly used for unsupervised tasks. In this review, we do not exhaustively discuss the different types of deep neural network architectures; an overview of the principal terms used herein is given in Table 1. Table 1 also provides select example applications, though in practice each neural network architecture has been broadly applied across multiple types of biomedical data. A recent book from Goodfellow et al. covers neural network architectures in detail [10], and LeCun et al. provide a more general introduction [2].

**Figure 1:**
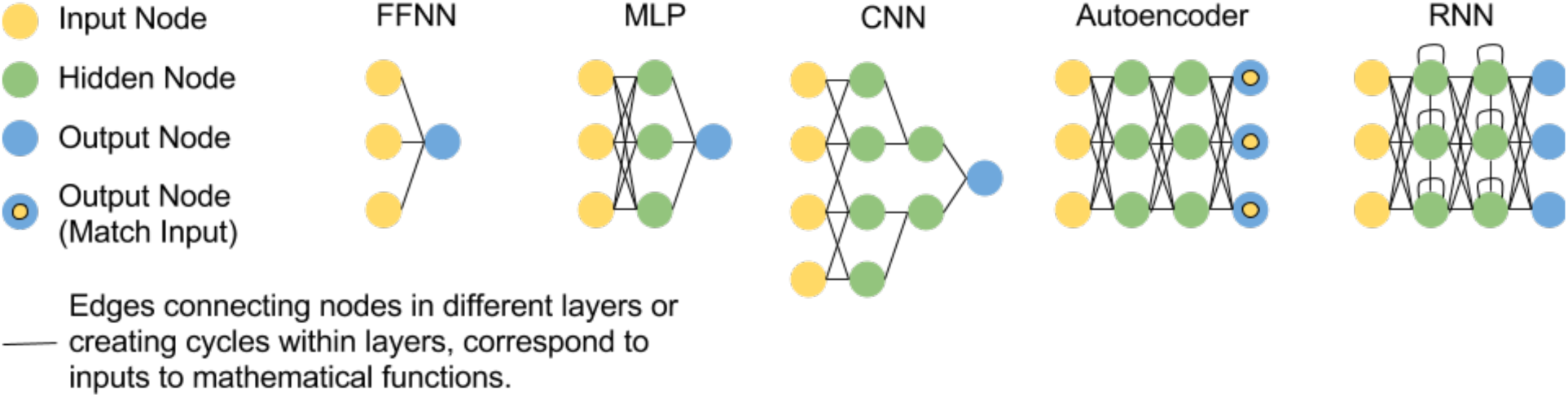
Neural networks come in many different forms. Left: a key for the various types of nodes used in neural networks. Simple FFNN: a feed forward neural network in which inputs are connected via some function to an output node and the model is trained to produce some output for a set of inputs. MLP: the multi-layer perceptron is a feed forward neural network in which there is at least one hidden layer between the input and output nodes. CNN: the convolutional neural network is a feed forward neural network in which the inputs are grouped spatially into hidden nodes. In the case of this example, each input node is only connected to hidden nodes alongside their neighboring input node. Autoencoder: a type of MLP in which the neural network is trained to produce an output that matches the input to the network. RNN: a deep recurrent neural network is used to allow the neural network to retain memory over time or sequential inputs. This figure was inspired by the Neural Network Zoo by Fjodor Van Veen.

**Table 1.** Glossary.

| Term | Definition | Example applications |
| --- | --- | --- |
| Supervised learning | Machine-learning approaches with goal of prediction of labels or outcomes |  |
| Unsupervised learning | Machine-learning approaches with goal of data summarization or pattern identification |  |
| Neural network (NN) | Machine-learning approach inspired by biological neurons where inputs are fed into one or more layers, producing an output layer |  |
| Deep neural network | NN with multiple hidden layers. Training happens over the network, and consequently such architectures allow for feature construction to occur alongside optimization of the overall training objective. |  |
| Feed-forward neural network (FFNN) | NN that does not have cycles between nodes in the same layer | Most of the examples below are special cases of FFNNs, except recurrent neural networks. |
| Multi-layer perceptron (MLP) | Type of FFNN with at least one hidden layer where each deeper layer is a nonlinear function of each earlier layer | MLPs do not impose structure and are frequently used when there is no natural ordering of the inputs (e.g. as with gene expression measurements). |
| Convolutional neural network (CNN) | A NN with layers in which connectivity preserves local structure. *If the data meet the underlying assumptions* performance is often good, and such networks can require fewer examples to train effectively because they have fewer parameters and also provide improved efficiency. | CNNs are used for sequence data—such as DNA sequences—or grid data—such as medical and microscopy images. |
| Recurrent neural network (RNN) | A neural network with cycles between nodes within a hidden layer. | The RNN architecture is used for sequential data—such as clinical time series and text or genome sequences. |
| Long short-term memory (LSTM) neural network | This special type of RNN has features that enable models to capture longer-term dependencies. | LSTMs are gaining a substantial foothold in the analysis of natural language, and may become more widely applied to biological sequence data. |
| Autoencoder (AE) | A NN where the training objective is to minimize the error between the output layer and the input layer. Such neural networks are unsupervised and are often used for dimensionality reduction. | Autoencoders have been used for unsupervised analysis of gene expression data as well as data extracted from the electronic health record. |
| Variational autoencoder (VAE) | This special type of AE has the added constraint that the model is trained to learn normally-distributed features. | VAEs have a track record of producing a valuable reduced representation in the imaging domain, and some early publications have used VAEs to analyze gene expression data. |
| Denoising autoencoder (DA) | This special type of AE includes a step where noise is added to the input during the training process. The denoising step acts as smoothing and may allow for effective use on input data that is inherently noisy. | Like AEs, DAs have been used for unsupervised analysis of gene expression data as well as data extracted from the electronic health record. |
| Generative neural network | Neural networks that fall into this class can be used to generate data similar to input data. These models can be sampled to produce hypothetical examples. | A number of the unsupervised learning neural network architectures that are summarized here can be used in a generative fashion. |
| Restricted Boltzmann machine (RBM) | A generative NN that forms the building block for many deep learning approaches, having a single input layer and a single hidden layer, with no connections between the nodes within each layer | RBMs have been applied to combine multiple types of omic data (e.g. DNA methylation, mRNA expression, and miRNA expression). |
| Deep belief network (DBN) | Generative NN with several hidden layers, which can be obtained from combining multiple RBMs | DBNs can be used to predict new relationships in a drug-target interaction network. |
| Generative adversarial network (GAN) | A generative NN approach where two neural networks are trained. One neural network, the generator, is provided with a set of randomly generated inputs and tasked with generating samples. The second, the discriminator, is trained to differentiate real and generated samples. After the two neural networks are trained against each other, the resulting generator can be used to produce new examples. | GANs can synthesize new examples with the same statistical properties of datasets that contain individual-level records and are subject to sharing restrictions. They have also been applied to generate microscopy images. |
| Adversarial training | A process by which artificial training examples are maliciously designed to fool a NN and then input as training examples to make the resulting NN robust (no relation to GANs) | Adversarial training has been used in image analysis. |
| Data augmentation | A process by which transformations that do not affect relevant properties of the input data (e.g. arbitrary rotations of histopathology images) are applied to training examples to increase the size of the training set. | Data augmentation is widely used in the analysis of images because rotation transformations for biomedical images often do not change relevant properties of the image. |

While deep learning shows increased flexibility over other machine learning approaches, as seen in the remainder of this review, it requires large training sets in order to fit the hidden layers, as well as accurate labels for the supervised learning applications. For these reasons, deep learning has recently become popular in some areas of biology and medicine, while having lower adoption in other areas. At the same time, this highlights the potentially even larger role that it may play in future research, given the increases in data in all biomedical fields. It is also important to see it as a branch of machine learning and acknowledge that it has the same limitations as other approaches in that field. In particular, the results are still dependent on the underlying study design and the usual caveats of correlation versus causation still apply—a more precise answer is only better than a less precise one if it answers the correct question.

## Will deep learning transform the study of human disease?

With this review, we ask the question: what is needed for deep learning to transform how we categorize, study, and treat individuals to maintain or restore health? We choose a high bar for “transform.” Andrew Grove, the former CEO of Intel, coined the term Strategic Inflection Point to refer to a change in technologies or environment that requires a business to be fundamentally reshaped [11]. Here, we seek to identify whether deep learning is an innovation that can induce a Strategic Inflection Point in the practice of biology or medicine.

There are already a number of reviews focused on applications of deep learning in biology [12–16], healthcare [17, 18], and drug discovery [19–22]. Under our guiding question, we sought to highlight cases where deep learning enabled researchers to solve challenges that were previously considered infeasible or makes difficult, tedious analyses routine. We also identified approaches that researchers are using to sidestep challenges posed by biomedical data. We find that domain-specific considerations have greatly influenced how to best harness the power and flexibility of deep learning. Model interpretability is often critical. Understanding the patterns in data may be just as important as fitting the data. In addition, there are important and pressing questions about how to build networks that efficiently represent the underlying structure and logic of the data. Domain experts can play important roles in designing networks to represent data appropriately, encoding the most salient prior knowledge and assessing success or failure. There is also great potential to create deep learning systems that augment biologists and clinicians by prioritizing experiments or streamlining tasks that do not require expert judgment. We have divided the large range of topics into three broad classes: Disease and Patient Categorization, Fundamental Biological Study, and Treatment of Patients. Below, we briefly introduce the types of questions, approaches and data that are typical for each class in the application of deep learning.

### Disease and patient categorization

A key challenge in biomedicine is the accurate classification of diseases and disease subtypes. In oncology, current “gold standard” approaches include histology, which requires interpretation by experts, or assessment of molecular markers such as cell surface receptors or gene expression. One example is the PAM50 approach to classifying breast cancer where the expression of 50 marker genes divides breast cancer patients into four subtypes. Substantial heterogeneity still remains within these four subtypes [23, 24]. Given the increasing wealth of molecular data available, a more comprehensive subtyping seems possible. Several studies have used deep learning methods to better categorize breast cancer patients: For instance, denoising autoencoders, an unsupervised approach, can be used to cluster breast cancer patients [25], and CNN can help count mitotic divisions, a feature that is highly correlated with disease outcome in histological images [26]. Despite these recent advances, a number of challenges exist in this area of research, most notably the integration of molecular and imaging data with other disparate types of data such as electronic health records (EHRs).

### Fundamental biological study

Deep learning can be applied to answer more fundamental biological questions; it is especially suited to leveraging large amounts of data from high-throughput “omics” studies. One classic biological problem where machine learning, and now deep learning, has been extensively applied is molecular target prediction. For example, deep recurrent neural networks (RNNs) have been used to predict gene targets of microRNAs [27], and CNNs have been applied to predict protein residue-residue contacts and secondary structure [28–30]. Other recent exciting applications of deep learning include recognition of functional genomic elements such as enhancers and promoters [31–33] and prediction of the deleterious effects of nucleotide polymorphisms [34].

### Treatment of patients

Although the application of deep learning to patient treatment is just beginning, we expect new methods to recommend patient treatments, predict treatment outcomes, and guide the development of new therapies. One type of effort in this area aims to identify drug targets and interactions or predict drug response. Another uses deep learning on protein structures to predict drug interactions and drug bioactivity [35]. Drug repositioning using deep learning on transcriptomic data is another exciting area of research [36]. Restricted Boltzmann machines (RBMs) can be combined into deep belief networks (DBNs) to predict novel drug-target interactions and formulate drug repositioning hypotheses [37, 38]. Finally, deep learning is also prioritizing chemicals in the early stages of drug discovery for new targets [22].

## Deep learning and patient categorization

In healthcare, individuals are diagnosed with a disease or condition based on symptoms, the results of certain diagnostic tests, or other factors. Once diagnosed with a disease, an individual might be assigned a stage based on another set of human-defined rules. While these rules are refined over time, the process is evolutionary and ad hoc, potentially impeding the identification of underlying biological mechanisms and their corresponding treatment interventions.

Deep learning methods applied to a large corpus of patient phenotypes may provide a meaningful and more data-driven approach to patient categorization. For example, they may identify new shared mechanisms that would otherwise be obscured due to ad hoc historical definitions of disease. Perhaps deep neural networks, by reevaluating data without the context of our assumptions, can reveal novel classes of treatable conditions.

In spite of such optimism, the ability of deep learning models to indiscriminately extract predictive signals must also be assessed and operationalized with care. Imagine a deep neural network is provided with clinical test results gleaned from electronic health records. Because physicians may order certain tests based on their suspected diagnosis, a deep neural network may learn to “diagnose” patients simply based on the tests that are ordered. For some objective functions, such as predicting an International Classification of Diseases (ICD) code, this may offer good performance even though it does not provide insight into the underlying disease beyond physician activity. This challenge is not unique to deep learning approaches; however, it is important for practitioners to be aware of these challenges and the possibility in this domain of constructing highly predictive classifiers of questionable actual utility.

Our goal in this section is to assess the extent to which deep learning is already contributing to the discovery of novel categories. Where it is not, we focus on barriers to achieving these goals. We also highlight approaches that researchers are taking to address challenges within the field, particularly with regards to data availability and labeling.

### Imaging applications in healthcare

Deep learning methods have transformed the analysis of natural images and video, and similar examples are beginning to emerge with medical images. Deep learning has been used to classify lesions and nodules; localize organs, regions, landmarks and lesions; segment organs, organ substructures and lesions; retrieve images based on content; generate and enhance images; and combine images with clinical reports [18, 39].

Though there are many commonalities with the analysis of natural images, there are also key differences. In all cases that we examined, fewer than one million images were available for training, and datasets are often many orders of magnitude smaller than collections of natural images. Researchers have developed subtask-specific strategies to address this challenge.

Data augmentation provides an effective strategy for working with small training sets. The practice is exemplified by a series of papers that analyze images from mammographies [40–44]. To expand the number and diversity of images, researchers constructed adversarial training examples [43]. Adversarial training examples are constructed by applying a transformation that changes training images but not their content—for example by rotating an image by a random amount. An alternative in the domain is to train towards human-created features before subsequent fine-tuning [41], which can help to sidestep this challenge though it does give up deep learning techniques’ strength as feature constructors.

A second strategy repurposes features extracted from natural images by deep learning models, such as ImageNet [45], for new purposes. Diagnosing diabetic retinopathy through color fundus images became an area of focus for deep learning researchers after a large labeled image set was made publicly available during a 2015 Kaggle competition [46]. Most participants trained neural networks from scratch [46–48], but Gulshan et al. [49] repurposed a 48-layer Inception-v3 deep architecture pre-trained on natural images and surpassed the state-of-the-art specificity and sensitivity. Such features were also repurposed to detect melanoma, the deadliest form of skin cancer, from dermoscopic [50, 51] and non-dermoscopic images of skin lesions [4,52,53] as well as age-related macular degeneration [54]. Pre-training on natural images can enable very deep networks to succeed without overfitting. For the melanoma task, reported performance was competitive with or better than a board of certified dermatologists [4, 50]. Reusing features from natural images is also an emerging approach for radiographic images, where datasets are often too small to train large deep neural networks without these techniques [55–58]. A deep CNN trained on natural images boosts performance in radiographic images [57]. However, the target task required either re-training the initial model from scratch with special pre-processing or fine-tuning of the whole network on radiographs with heavy data augmentation to avoid overfitting.

The technique of reusing features from a different task falls into the broader area of transfer learning (see Discussion). Though we’ve mentioned numerous successes for the transfer of natural image features to new tasks, we expect that a lower proportion of negative results have been published. The analysis of magnetic resonance images (MRIs) is also faced with the challenge of small training sets. In this domain, Amit et al. [59] investigated the tradeoff between pre-trained models from a different domain and a small CNN trained only with MRI images. In contrast with the other selected literature, they found a smaller network trained with data augmentation on few hundred images from a few dozen patients can outperform a pre-trained out-of-domain classifier.

Another way of dealing with limited training data is to divide rich data—e.g. 3D images—into numerous reduced projections. Shin et al. [56] compared various deep network architectures, dataset characteristics, and training procedures for computer tomography-based (CT) abnormality detection. They concluded that networks as deep as 22 layers could be useful for 3D data, despite the limited size of training datasets. However, they noted that choice of architecture, parameter setting, and model fine-tuning needed is very problem-and dataset-specific. Moreover, this type of task often depends on both lesion localization and appearance, which poses challenges for CNN-based approaches. Straightforward attempts to capture useful information from full-size images in all three dimensions simultaneously via standard neural network architectures were computationally unfeasible. Instead, two-dimensional models were used to either process image slices individually (2D), or aggregate information from a number of 2D projections in the native space (2.5D).

Roth et al. compared 2D, 2.5D, and 3D CNNs on a number of tasks for computer-aided detection from CT scans and showed that 2.5D CNNs performed comparably well to 3D analogs, while requiring much less training time, especially on augmented training sets [60]. Another advantage of 2D and 2.5D networks is the wider availability of pre-trained models. But reducing the dimensionality is not always helpful. Nie et al. [61] showed that multimodal, multi-channel 3D deep architecture was successful at learning high-level brain tumor appearance features jointly from MRI, functional MRI, and diffusion MRI images, outperforming single-modality or 2D models. Overall, the variety of modalities, properties and sizes of training sets, the dimensionality of input, and the importance of end goals in medical image analysis are provoking a development of specialized deep neural network architectures, training and validation protocols, and input representations that are not characteristic of widely-studied natural images.

Predictions from deep neural networks can be evaluated for use in workflows that also incorporate human experts. In a large dataset of mammography images, Kooi et al. [62] demonstrated that deep neural networks outperform the traditional computer-aided diagnosis system at low sensitivity and perform comparably at high sensitivity. They also compared network performance to certified screening radiologists on a patch level and found no significant difference between the network and the readers. However, using deep methods for clinical practice is challenged by the difficulty of assigning a level of confidence to each prediction. Leibig et al. [48] estimated the uncertainty of deep networks for diabetic retinopathy diagnosis by linking dropout networks with approximate Bayesian inference. Techniques that assign confidences to each prediction should aid pathologist-computer interactions and improve uptake by physicians.

Systems to aid in the analysis of histology slides are also promising use cases for deep learning [63]. Ciresan et al. [26] developed one of the earliest approaches for histology slides, winning the 2012 International Conference on Pattern Recognition’s Contest on Mitosis Detection while achieving human-competitive accuracy. In more recent work, Wang et al. [64] analyzed stained slides of lymph node slices to identify cancers. On this task a pathologist has about a 3% error rate. The pathologist did not produce any false positives, but did have a number of false negatives. The algorithm had about twice the error rate of a pathologist, but the errors were not strongly correlated. In this area, these algorithms may be ready to be incorporated into existing tools to aid pathologists and reduce the false negative rate. Ensembles of deep learning and human experts may help overcome some of the challenges presented by data limitations.

One source of training examples with rich phenotypical annotations is the EHR. Billing information in the form of ICD codes are simple annotations but phenotypic algorithms can combine laboratory tests, medication prescriptions, and patient notes to generate more reliable phenotypes. Recently, Lee et al. [65] developed an approach to distinguish individuals with age-related macular degeneration from control individuals. They trained a deep neural network on approximately 100,000 images extracted from structured electronic health records, reaching greater than 93% accuracy. The authors used their test set to evaluate when to stop training. In other domains, this has resulted in a minimal change in the estimated accuracy [66], but we recommend the use of an independent test set whenever feasible.

Rich clinical information is stored in EHRs. However, manually annotating a large set requires experts and is time consuming. For chest X-ray studies, a radiologist usually spends a few minutes per example. Generating the number of examples needed for deep learning is infeasibly expensive. Instead, researchers may benefit from using text mining to generate annotations [67], even if those annotations are of modest accuracy. Wang et al. [68] proposed to build predictive deep neural network models through the use of images with *weak labels*. Such labels are automatically generated and not verified by humans, so they may be noisy or incomplete. In this case, they applied a series of natural language processing (NLP) techniques to the associated chest X-ray radiological reports. They first extracted all diseases mentioned in the reports using a state-of-the-art NLP tool, then applied a new method, NegBio [69], to filter negative and equivocal findings in the reports. Evaluation on four independent datasets demonstrated that NegBio is highly accurate for detecting negative and equivocal findings (~90% in F₁ score, which balances precision and recall [70]). The resulting dataset [71] consisted of 112,120 frontal-view chest X-ray images from 30,805 patients, and each image was associated with one or more *text-mined* (weakly-labeled) pathology categories (e.g. pneumonia and cardiomegaly) or “no finding” otherwise. Further, Wang et al. [68] used this dataset with a unified weakly-supervised multi-label image classification framework, to detect common thoracic diseases. It showed superior performance over a benchmark using fully-labeled data.

With the exception of natural image-like problems (e.g. melanoma detection), biomedical imaging poses a number of challenges for deep learning. Datasets are typically small, annotations can be sparse, and images are often high-dimensional, multimodal, and multi-channel. Techniques like transfer learning, heavy dataset augmentation, and the use of multi-view and multi-stream architectures are more common than in the natural image domain. Furthermore, high model sensitivity and specificity can translate directly into clinical value. Thus, prediction evaluation, uncertainty estimation, and model interpretation methods are also of great importance in this domain (see Discussion). Finally, there is a need for better pathologist-computer interaction techniques that will allow combining the power of deep learning methods with human expertise and lead to better-informed decisions for patient treatment and care.

### Text applications in healthcare

Due to the rapid growth of scholarly publications and EHRs, biomedical text mining has become increasingly important in recent years. The main tasks in biological and clinical text mining include, but are not limited to, named entity recognition, relation/event extraction, and information retrieval (Figure 2). Deep learning is appealing in this domain because of its competitive performance versus traditional methods and ability to overcome challenges in feature engineering. Relevant applications can be stratified by the application domain (biomedical literature vs. clinical notes) and the actual task (e.g. concept or relation extraction).

**Figure 2:**
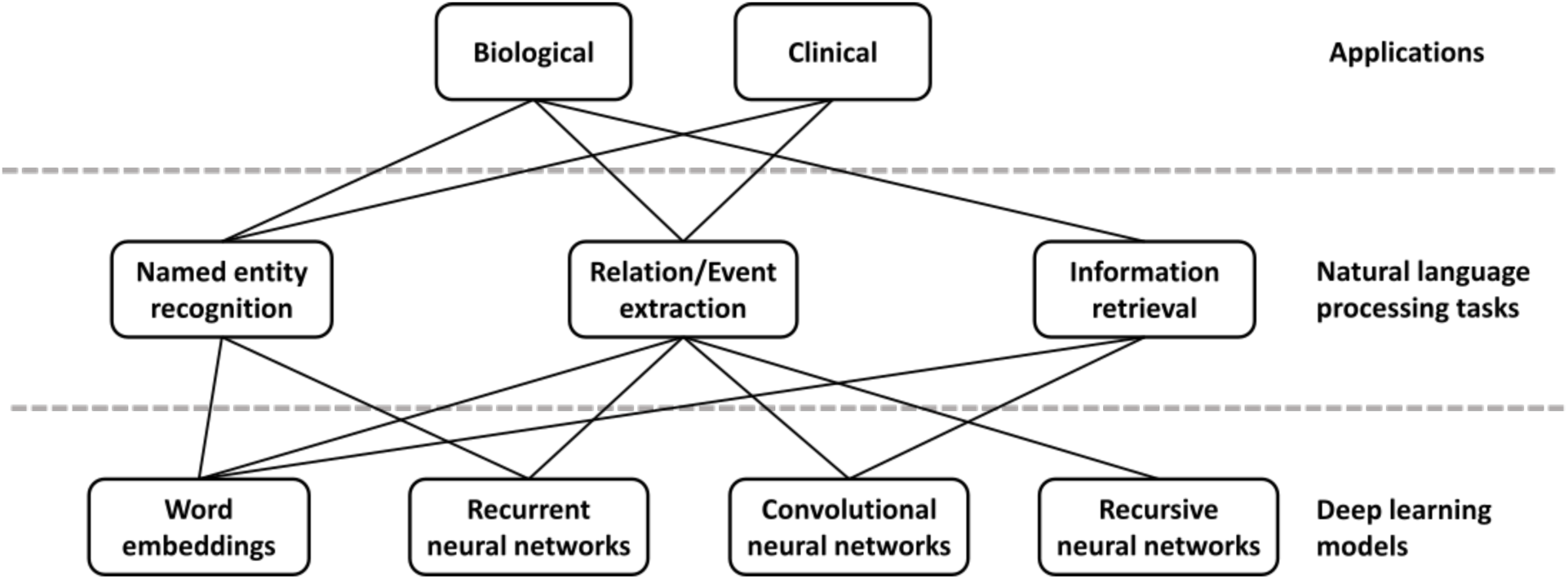
Deep learning applications, tasks, and models based on NLP perspectives.

Named entity recognition (NER) is a task of identifying text spans that refer to a biological concept of a specific class, such as disease or chemical, in a controlled vocabulary or ontology. NER is often needed as a first step in many complex text mining systems. The current state-of-the-art methods typically reformulate the task as a sequence labeling problem and use conditional random fields [72–74]. In recent years, word embeddings that contain rich latent semantic information of words have been widely used to improve the NER performance. Liu et al. studied the effect of word embeddings on drug name recognition and compared them with traditional semantic features [75]. Tang et al. investigated word embeddings in gene, DNA, and cell line mention detection tasks [76]. Moreover, Wu et al. examined the use of neural word embeddings for clinical abbreviation disambiguation [77]. Liu et al. exploited task-oriented resources to learn word embeddings for clinical abbreviation expansion [78].

Relation extraction involves detecting and classifying semantic relationships between entities from the literature. At present, kernel methods or feature-based approaches are commonly applied [79–81]. Deep learning can relieve the feature sparsity and engineering problems. Some studies focused on jointly extracting biomedical entities and relations simultaneously [82, 83], while others applied deep learning on relation classification given the relevant entities. For example, both multichannel dependency-based CNNs [84] and shortest path-based CNNs [85, 86] are well-suited for sentence-based protein-protein extraction. Jiang et al. proposed a biomedical domain-specific word embedding model to reduce the manual labor of designing semantic representation for the same task [87]. Gu et al. employed a maximum entropy model and a CNN model for chemical-induced disease relation extraction at the inter-and intra-sentence level, respectively [88]. For drug-drug interaction, Zhao et al. used a CNN that employs word embeddings with the syntactic information of a sentence as well as features of part-of-speech tags and dependency trees [89]. Asada et al. experimented with an attention CNN [90], and Yi et al. a RNN model with multiple attention layers [91]. In both cases, it is a single model with attention mechanism, which allows the decoder to focus on different parts of the source sentence. As a result, it does not require dependency parsing or training multiple models. Both attention CNN and RNN have comparable results, but the CNN model has an advantage in that it can be easily computed in parallel, hence making it faster with recent graphics processing units (GPUs).

For biotopes event extraction, Li et al. employed CNN and distributed representation [92] while Mehryary et al. used long short-term memory (LSTM) networks to extract complicated relations [93]. Li et al. applied word embedding to extract complete events from biomedical text and achieved results comparable to the state-of-the-art systems [94]. There are also approaches that identify event triggers rather than the complete event [95, 96]. Taken together, deep learning models outperform traditional kernel methods or feature-based approaches by 1–5% in f-score. Among various deep learning approaches, CNN stands out as the most popular model both in terms of computational complexity and performance, while RNN has achieved continuous progress.

Information retrieval is a task of finding relevant text that satisfies an information need from within a large document collection. While deep learning has not yet achieved the same level of success in this area as seen in others, the recent surge of interest and work suggest that this may be quickly changing. For example, Mohan et al. described a deep learning approach to modeling the relevance of a document’s text to a query, which they applied to the entire biomedical literature [97].

To summarize, deep learning has shown promising results in many biomedical text mining tasks and applications. But to realize its full potential in this domain, either large size of labeled data or technical advancements in current methods coping with limited labeled data are required.

### Electronic health records

EHR data include substantial amounts of free text, which remains challenging to approach [98]. Often, researchers developing algorithms that perform well on specific tasks must design and implement domain-specific features [99]. These features capture unique aspects of the literature being processed. Deep learning methods are natural feature constructors. In recent work, the authors evaluated the extent to which deep learning methods could be applied on top of generic features for domain-specific concept extraction [100]. They found that performance was in line with, but lower than the best domain-specific method [100]. This raises the possibility that deep learning may impact the field by reducing the researcher time and cost required to develop specific solutions, but it may not always lead to performance increases.

In recent work, Yoon et al.[101] analyzed simple features using deep neural networks and found that the patterns recognized by the algorithms could be re-used across tasks. Their aim was to analyze the free text portions of pathology reports to identify the primary site and laterality of tumors. The only features the authors supplied to the algorithms were unigrams (counts for single words) and bigrams (counts for two-word combinations) in a free text document. They subset the full set of words and word combinations to the 400 most common. The machine learning algorithms that they employed (naïve Bayes, logistic regression, and deep neural networks) all performed relatively similarly on the task of identifying the primary site. However, when the authors evaluated the more challenging task, evaluating the laterality of each tumor, the deep neural network outperformed the other methods. Of particular interest, when the authors first trained a neural network to predict primary site and then repurposed those features as a component of a secondary neural network trained to predict laterality, the performance was higher than a laterality-trained neural network. This demonstrates how deep learning methods can repurpose features across tasks, improving overall predictions as the field tackles new challenges. The Discussion further reviews this type of transfer learning.

Several authors have created reusable feature sets for medical terminologies using natural language processing and neural embedding models, as popularized by word2vec [102]. Minarro-Giménez et al. [103] applied the word2vec deep learning toolkit to medical corpora and evaluated the efficiency of word2vec in identifying properties of pharmaceuticals based on mid-sized, unstructured medical text corpora without any additional background knowledge. A goal of learning terminologies for different entities in the same vector space is to find relationships between different domains (e.g. drugs and the diseases they treat). It is difficult for us to provide a strong statement on the broad utility of these methods. Manuscripts in this area tend to compare algorithms applied to the same data but lack a comparison against overall best-practices for one or more tasks addressed by these methods. Techniques have been developed for free text medical notes [104], ICD and National Drug Codes [105, 106], and claims data [107]. Methods for neural embeddings learned from electronic health records have at least some ability to predict disease-disease associations and implicate genes with a statistical association with a disease [108], but the evaluations performed did not differentiate between simple predictions (i.e. the same disease in different sites of the body) and non-intuitive ones. Jagannatha and Yu [109] further employed a bidirectional LSTM structure to extract adverse drug events from electronic health records, and Lin et al. [110] investigated using CNN to extract temporal relations. While promising, a lack of rigorous evaluations of the real-world utility of these kinds of features makes current contributions in this area difficult to evaluate. Comparisons need to be performed to examine the true utility against leading approaches (i.e. algorithms and data) as opposed to simply evaluating multiple algorithms on the same potentially limited dataset.

Identifying consistent subgroups of individuals and individual health trajectories from clinical tests is also an active area of research. Approaches inspired by deep learning have been used for both unsupervised feature construction and supervised prediction. Early work by Lasko et al. [111], combined sparse autoencoders and Gaussian processes to distinguish gout from leukemia from uric acid sequences. Later work showed that unsupervised feature construction of many features via denoising autoencoder neural networks could dramatically reduce the number of labeled examples required for subsequent supervised analyses [112]. In addition, it pointed towards features learned during unsupervised training being useful for visualizing and stratifying subgroups of patients within a single disease. In a concurrent large-scale analysis of EHR data from 700,000 patients, Miotto et al. [113] used a deep denoising autoencoder architecture applied to the number and co-occurrence of clinical events to learn a representation of patients (DeepPatient). The model was able to predict disease trajectories within one year with over 90% accuracy and patient-level predictions were improved by up to 15% when compared to other methods. Choi et al. [114] attempted to model the longitudinal structure of EHRs with a RNN to predict future diagnosis and medication prescriptions on a cohort of 260,000 patients followed for 8 years (Doctor AI). Pham et al. [115] built upon this concept by using a RNN with a LSTM architecture enabling explicit modelling of patient trajectories through the use of memory cells. The method, DeepCare, performed better than shallow models or plain RNN when tested on two independent cohorts for its ability to predict disease progression, intervention recommendation and future risk prediction. Nguyen et al. [116] took a different approach and used word embeddings from EHRs to train a CNN that could detect and pool local clinical motifs to predict unplanned readmission after six months, with performance better than the baseline method (Deepr). Razavian et al. [117] used a set of 18 common lab tests to predict disease onset using both CNN and LSTM architectures and demonstrated an improvement over baseline regression models. However, numerous challenges including data integration (patient demographics, family history, laboratory tests, text-based patient records, image analysis, genomic data) and better handling of streaming temporal data with many features, will need to be overcome before we can fully assess the potential of deep learning for this application area.

Still, recent work has also revealed domains in which deep networks have proven superior to traditional methods. Survival analysis models the time leading to an event of interest from a shared starting point, and in the context of EHR data, often associates these events to subject covariates. Exploring this relationship is difficult, however, given that EHR data types are often heterogeneous, covariates are often missing, and conventional approaches require the covariate-event relationship be linear and aligned to a specific starting point [118]. Early approaches, such as the Faraggi-Simon feed-forward network, aimed to relax the linearity assumption, but performance gains were lacking [119]. Katzman et al. in turn developed a deep implementation of the Faraggi-Simon network that, in addition to outperforming Cox regression, was capable of comparing the risk between a given pair of treatments, thus potentially acting as recommender system [120]. To overcome the remaining difficulties, researchers have turned to deep exponential families, a class of latent generative models that are constructed from any type of exponential family distributions [121]. The result was a deep survival analysis model capable of overcoming challenges posed by missing data and heterogeneous data types, while uncovering nonlinear relationships between covariates and failure time. They showed their model more accurately stratified patients as a function of disease risk score compared to the current clinical implementation.

There is a computational cost for these methods, however, when compared to traditional, non-neural network approaches. For the exponential family models, despite their scalability [122], an important question for the investigator is whether he or she is interested in estimates of posterior uncertainty. Given that these models are effectively Bayesian neural networks, much of their utility simplifies to whether a Bayesian approach is warranted for a given increase in computational cost. Moreover, as with all variational methods, future work must continue to explore just how well the posterior distributions are approximated, especially as model complexity increases [123].

## Challenges and opportunities in patient categorization

### Generating ground-truth labels can be expensive or impossible

A dearth of true labels is perhaps among the biggest obstacles for EHR-based analyses that employ machine learning. Popular deep learning (and other machine learning) methods are often used to tackle classification tasks and thus require ground-truth labels for training. For EHRs this can mean that researchers must hire multiple clinicians to manually read and annotate individual patients’ records through a process called chart review. This allows researchers to assign “true” labels, i.e. those that match our best available knowledge. Depending on the application, sometimes the features constructed by algorithms also need to be manually validated and interpreted by clinicians. This can be time consuming and expensive [124]. Because of these costs, much of this research, including the work cited in this review, skips the process of expert review. Clinicians’ skepticism for research without expert review may greatly dampen their enthusiasm for the work and consequently reduce its impact. To date, even well-resourced large national consortia have been challenged by the task of acquiring enough expert-validated labeled data. For instance, in the eMERGE consortia and PheKB database [125], most samples with expert validation contain only 100 to 300 patients. These datasets are quite small even for simple machine learning algorithms. The challenge is greater for deep learning models with many parameters. While unsupervised and semi-supervised approaches can help with small sample sizes, the field would benefit greatly from large collections of anonymized records in which a substantial number of records have undergone expert review. This challenge is not unique to EHR-based studies. Work on medical images, omics data in applications for which detailed metadata are required, and other applications for which labels are costly to obtain will be hampered as long as abundant curated data are unavailable.

Successful approaches to date in this domain have sidestepped this challenge by making methodological choices that either reduce the need for labeled examples or that use transformations to training data to increase the number of times it can be used before overfitting occurs. For example, the unsupervised and semi-supervised methods that we have discussed reduce the need for labeled examples [112]. The anchor and learn framework [126] uses expert knowledge to identify high-confidence observations from which labels can be inferred. The strategies of adversarial training mentioned above can reduce overfitting, if transformations are available that preserve the meaningful content of the data while transforming irrelevant features [43]. While adversarial training examples can be easily imagined for certain methods that operate on images, it is more challenging to figure out what an equivalent transformation would be for a patient’s clinical test results. Consequently, it may be hard to employ adversarial training examples with other applications. Finally, approaches that transfer features can also help use valuable training data most efficiently. Rajkomar et al. trained a deep neural network using generic images before tuning using only radiology images [57]. Datasets that require many of the same types of features might be used for initial training, before fine tuning takes place with the more sparse biomedical examples. Though the analysis has not yet been attempted, it is possible that analogous strategies may be possible with electronic health records. For example, features learned from the electronic health record for one type of clinical test (e.g. a decrease over time in a lab value) may transfer across phenotypes. Methods to accomplish more with little high-quality labeled data arose in other domains and may also be adapted to this challenge, e.g. data programming [127]. In data programming, noisy automated labeling functions are integrated.

Numerous commentators have described data as the new oil [128, 129]. The idea behind this metaphor is that data are available in large quantities, valuable once refined, and this underlying resource will enable a data-driven revolution in how work is done. Contrasting with this perspective, Ratner, Bach, and Ré described labeled training data, instead of data, as “The *New* New Oil” [130]. In this framing, data are abundant and not a scarce resource. Instead, new approaches to solving problems arise when labeled training data become sufficient to enable them. Based on our review of research on deep learning methods to categorize disease, the latter framing rings true.

We expect improved methods for domains with limited data to play an important role if deep learning is going to transform how we categorize states of human health. We don’t expect that deep learning methods will replace expert review. We expect them to complement expert review by allowing more efficient use of the costly practice of manual annotation.

### Data sharing is hampered by standardization and privacy considerations

To construct the types of very large datasets that deep learning methods thrive on, we need robust sharing of large collections of data. This is in part a cultural challenge. We touch on this challenge in Discussion. Beyond the cultural hurdles around data sharing, there are also technological and legal hurdles related to sharing individual health records or deep models built from such records. This subsection deals primarily with these challenges.

EHRs are designed chiefly for clinical, administrative and financial purposes, such as patient care, insurance and billing [131]. Science is at best a tertiary priority, presenting challenges to EHR-based research in general and to deep learning research in particular. Although there is significant work in the literature around EHR data quality and the impact on research [132], we focus on three types of challenges: local bias, wider standards, and legal issues. Note these problems are not restricted to EHRs but can also apply to any large biomedical dataset, e.g. clinical trial data.

Even within the same healthcare system, EHRs can be used differently [133, 134]. Individual users have unique documentation and ordering patterns, with different departments and different hospitals having different priorities that code patients and introduce missing data in a non-random fashion [135]. Patient data may be kept across several “silos” within a single health system (e.g. separate nursing documentation, registries, etc.). Even the most basic task of matching patients across systems can be challenging due to data entry issues [136]. The situation is further exacerbated by the ongoing introduction, evolution, and migration of EHR systems, especially where reorganized and acquired healthcare facilities have to merge. Further, even the ostensibly least-biased data type, laboratory measurements, can be biased based by both the healthcare process and patient health state [137]. As a result, EHR data can be less complete and less objective than expected.

In the wider picture, standards for EHRs are numerous and evolving. Proprietary systems, indifferent and scattered use of health information standards, and controlled terminologies makes combining and comparison of data across systems challenging [138]. Further diversity arises from variation in languages, healthcare practices, and demographics. Merging EHRs gathered in different systems (and even under different assumptions) is challenging [139].

Combining or replicating studies across systems thus requires controlling for both the above biases and dealing with mismatching standards. This has the practical effect of reducing cohort size, limiting statistical significance, preventing the detection of weak effects [140], and restricting the number of parameters that can be trained in a model. Further, rules-based algorithms have been popular in EHR-based research, but because these are developed at a single institution and trained with a specific patient population, they do not transfer easily to other healthcare systems [141]. Genetic studies using EHR data are subject to even more bias, as the differences in population ancestry across health centers (e.g. proportion of patients with African or Asian ancestry) can affect algorithm performance. For example, Wiley et al. [142] showed that warfarin dosing algorithms often under-perform in African Americans, illustrating that some of these issues are unresolved even at a treatment best practices level. Lack of standardization also makes it challenging for investigators skilled in deep learning to enter the field, as numerous data processing steps must be performed before algorithms are applied.

Finally, even if data were perfectly consistent and compatible across systems, attempts to share and combine EHR data face considerable legal and ethical barriers. Patient privacy can severely restrict the sharing and use of EHR data [143]. Here again, standards are heterogeneous and evolving, but often EHR data can often not be exported or even accessed directly for research purposes without appropriate consent. In the United States, research use of EHR data is subject both to the Common Rule and the Health Insurance Portability and Accountability Act (HIPAA). Ambiguity in the regulatory language and individual interpretation of these rules can hamper use of EHR data [144]. Once again, this has the effect of making data gathering more laborious and expensive, reducing sample size and study power.

Several technological solutions have been proposed in this direction, allowing access to sensitive data satisfying privacy and legal concerns. Software like DataShield [145] and ViPAR [146], although not EHR-specific, allow querying and combining of datasets and calculation of summary statistics across remote sites by “taking the analysis to the data”. The computation is carried out at the remote site. Conversely, the EH4CR project [138] allows analysis of private data by use of an inter-mediation layer that interprets remote queries across internal formats and datastores and returns the results in a de-identified standard form, thus giving real-time consistent but secure access. Continuous Analysis [147] can allow reproducible computing on private data. Using such techniques, intermediate results can be automatically tracked and shared without sharing the original data. While none of these have been used in deep learning, the potential is there.

Even without sharing data, algorithms trained on confidential patient data may present security risks or accidentally allow for the exposure of individual level patient data. Tramer et al. [148] showed the ability to steal trained models via public application programming interfaces (APIs). Dwork and Roth [149] demonstrate the ability to expose individual level information from accurate answers in a machine learning model. Attackers can use similar attacks to find out if a particular data instance was present in the original training set for the machine learning model [150], in this case, whether a person’s record was present. To protect against these attacks, Simmons et al. [151] developed the ability to perform genome-wide association studies (GWASs) in a differentially private manner, and Abadi et al. [152] show the ability to train deep learning classifiers under the differential privacy framework.

These attacks also present a potential hazard for approaches that aim to generate data. Choi et al. propose generative adversarial neural networks (GANs) as a tool to make sharable EHR data [153], and Esteban et al. [154] showed that recurrent GANs could be used for time series data. However, in both cases the authors did not take steps to protect the model from such attacks. There are approaches to protect models, but they pose their own challenges. Training in a differentially private manner provides a limited guarantee that an algorithm’s output will be equally likely to occur regardless of the participation of any one individual. The limit is determined by parameters which provide a quantification of privacy. Beaulieu-Jones et al. demonstrated the ability to generate data that preserved properties of the SPRINT clinical trial with GANs under the differential privacy framework [155]. Both Beaulieu-Jones et al. and Esteban et al. train models on synthetic data generated under differentially private and observe performance from a transfer learning evaluation that is only slightly below models trained on the original, real data. Taken together, these results suggest that differentially private GANs may be an attractive way to generate sharable datasets for downstream reanalysis.

Federated learning [156] and secure aggregations [157] are complementary approaches that reinforce differential privacy. Both aim to maintain privacy by training deep learning models from decentralized data sources such as personal mobile devices without transferring actual training instances. This is becoming of increasing importance with the rapid growth of mobile health applications. However, the training process in these approaches places constraints on the algorithms used and can make fitting a model substantially more challenging. It can be trivial to train a model without differential privacy, but quite difficult to train one within the differential privacy framework [155]. This problem can be particularly pronounced with small sample sizes.

While none of these problems are insurmountable or restricted to deep learning, they present challenges that cannot be ignored. Technical evolution in EHRs and data standards will doubtless ease—although not solve—the problems of data sharing and merging. More problematic are the privacy issues. Those applying deep learning to the domain should consider the potential of inadvertently disclosing the participants’ identities. Techniques that enable training on data without sharing the raw data may have a part to play. Training within a differential privacy framework may often be warranted.

### Discrimination and “right to an explanation” laws

In April 2016, the European Union adopted new rules regarding the use of personal information, the General Data Protection Regulation [159]. A component of these rules can be summed up by the phrase “right to an explanation”. Those who use machine learning algorithms must be able to explain how a decision was reached. For example, a clinician treating a patient who is aided by a machine learning algorithm may be expected to explain decisions that use the patient’s data. The new rules were designed to target categorization or recommendation systems, which inherently profile individuals. Such systems can do so in ways that are discriminatory and unlawful.

As datasets become larger and more complex, we may begin to identify relationships in data that are important for human health but difficult to understand. The algorithms described in this review and others like them may become highly accurate and useful for various purposes, including within medical practice. However, to discover and avoid discriminatory applications it will be important to consider interpretability alongside accuracy. A number of properties of genomic and healthcare data will make this difficult.

First, research samples are frequently non-representative of the general population of interest; they tend to be disproportionately sick [160], male [161], and European in ancestry [162]. One well-known consequence of these biases in genomics is that penetrance is consistently lower in the general population than would be implied by case-control data, as reviewed in [160]. Moreover, real genetic associations found in one population may not hold in other populations with different patterns of linkage disequilibrium (even when population stratification is explicitly controlled for [163]). As a result, many genomic findings are of limited value for people of non-European ancestry [162] and may even lead to worse treatment outcomes for them. Methods have been developed for mitigating some of these problems in genomic studies [160, 163], but it is not clear how easily they can be adapted for deep models that are designed specifically to extract subtle effects from high-dimensional data. For example, differences in the equipment that tended to be used for cases versus controls have led to spurious genetic findings (e.g. Sebastiani et al.’s retraction [164]). In some contexts, it may not be possible to correct for all of these differences to the degree that a deep network is unable to use them. Moreover, the complexity of deep networks makes it difficult to determine when their predictions are likely to be based on such nominally-irrelevant features of the data (called “leakage” in other fields [165]). When we are not careful with our data and models, we may inadvertently say more about the way the data was collected (which may involve a history of unequal access and discrimination) than about anything of scientific or predictive value. This fact can undermine the privacy of patient data [165] or lead to severe discriminatory consequences [166].

There is a small but growing literature on the prevention and mitigation of data leakage [165], as well as a closely-related literature on discriminatory model behavior [167], but it remains difficult to predict when these problems will arise, how to diagnose them, and how to resolve them in practice. There is even disagreement about which kinds of algorithmic outcomes should be considered discriminatory [168]. Despite the difficulties and uncertainties, machine learning practitioners (and particularly those who use deep neural networks, which are challenging to interpret) must remain cognizant of these dangers and make every effort to prevent harm from discriminatory predictions. To reach their potential in this domain, deep learning methods will need to be interpretable (see Discussion). Researchers need to consider the extent to which biases may be learned by the model and whether or not a model is sufficiently interpretable to identify bias. We discuss the challenge of model interpretability more thoroughly in Discussion.

### Applications of deep learning to longitudinal analysis

Longitudinal analysis follows a population across time, for example, prospectively from birth or from the onset of particular conditions. In large patient populations, longitudinal analyses such as the Framingham Heart Study [169] and the Avon Longitudinal Study of Parents and Children [170] have yielded important discoveries about the development of disease and the factors contributing to health status. Yet, a common practice in EHR-based research is to take a snapshot at a point in time and convert patient data to a traditional vector for machine learning and statistical analysis. This results in loss of information as timing and order of events can provide insight into a patient’s disease and treatment [171]. Efforts to model sequences of events have shown promise [172] but require exceedingly large patient sizes due to discrete combinatorial bucketing. Lasko et al. [111] used autoencoders on longitudinal sequences of serum uric acid measurements to identify population subtypes. More recently, deep learning has shown promise working with both sequences (CNNs) [173] and the incorporation of past and current state (RNNs, LSTMs) [115]. This may be a particular area of opportunity for deep neural networks. The ability to recognize relevant sequences of events from a large number of trajectories requires powerful and flexible feature construction methods—an area in which deep neural networks excel.

## Deep learning to study the fundamental biological processes underlying human disease

The study of cellular structure and core biological processes—transcription, translation, signaling, metabolism, etc.—in humans and model organisms will greatly impact our understanding of human disease over the long horizon [174]. Predicting how cellular systems respond to environmental perturbations and are altered by genetic variation remain daunting tasks. Deep learning offers new approaches for modeling biological processes and integrating multiple types of omic data [175], which could eventually help predict how these processes are disrupted in disease. Recent work has already advanced our ability to identify and interpret genetic variants, study microbial communities, and predict protein structures, which also relates to the problems discussed in the drug development section. In addition, unsupervised deep learning has enormous potential for discovering novel cellular states from gene expression, fluorescence microscopy, and other types of data that may ultimately prove to be clinically relevant.

Progress has been rapid in genomics and imaging, fields where important tasks are readily adapted to well-established deep learning paradigms. One-dimensional convolutional and recurrent neural networks are well-suited for tasks related to DNA-and RNA-binding proteins, epigenomics, and RNA splicing. Two dimensional CNNs are ideal for segmentation, feature extraction, and classification in fluorescence microscopy images [16]. Other areas, such as cellular signaling, are biologically important but studied less-frequently to date, with some exceptions [176]. This may be a consequence of data limitations or greater challenges in adapting neural network architectures to the available data. Here, we highlight several areas of investigation and assess how deep learning might move these fields forward.

### Gene expression

Gene expression technologies characterize the abundance of many thousands of RNA transcripts within a given organism, tissue, or cell. This characterization can represent the underlying state of the given system and can be used to study heterogeneity across samples as well as how the system reacts to perturbation. While gene expression measurements were traditionally made by quantitative polymerase chain reaction (qPCR), low-throughput fluorescence-based methods, and microarray technologies, the field has shifted in recent years to primarily performing RNA sequencing (RNA-seq) to catalog whole transcriptomes. As RNA-seq continues to fall in price and rise in throughput, sample sizes will increase and training deep models to study gene expression will become even more useful.

Already several deep learning approaches have been applied to gene expression data with varying aims. For instance, many researchers have applied unsupervised deep learning models to extract meaningful representations of gene modules or sample clusters. Denoising autoencoders have been used to cluster yeast expression microarrays into known modules representing cell cycle processes [177] and to stratify yeast strains based on chemical and mutational perturbations [178]. Shallow (one hidden layer) denoising autoencoders have also been fruitful in extracting biological insight from thousands of *Pseudomonas aeruginosa* experiments [179, 180] and in aggregating features relevant to specific breast cancer subtypes [25]. These unsupervised approaches applied to gene expression data are powerful methods for identifying gene signatures that may otherwise be overlooked. An additional benefit of unsupervised approaches is that ground truth labels, which are often difficult to acquire or are incorrect, are nonessential. However, the genes that have been aggregated into features must be interpreted carefully. Attributing each node to a single specific biological function risks over-interpreting models. Batch effects could cause models to discover non-biological features, and downstream analyses should take this into consideration.

Deep learning approaches are also being applied to gene expression prediction tasks. For example, a deep neural network with three hidden layers outperformed linear regression in inferring the expression of over 20,000 target genes based on a representative, well-connected set of about 1,000 landmark genes [181]. However, while the deep learning model outperformed existing algorithms in nearly every scenario, the model still displayed poor performance. The paper was also limited by computational bottlenecks that required data to be split randomly into two distinct models and trained separately. It is unclear how much performance would have increased if not for computational restrictions.

Epigenomic data, combined with deep learning, may have sufficient explanatory power to infer gene expression. For instance, the DeepChrome CNN [182] improved prediction accuracy of high or low gene expression from histone modifications over existing methods. AttentiveChrome [183] added a deep attention model to further enhance DeepChrome. Deep learning can also integrate different data types. For example, Liang et al. combined RBMs to integrate gene expression, DNA methylation, and miRNA data to define ovarian cancer subtypes [184]. While these approaches are promising, many convert gene expression measurements to categorical or binary variables, thus ablating many complex gene expression signatures present in intermediate and relative numbers.

Deep learning applied to gene expression data is still in its infancy, but the future is bright. Many previously untestable hypotheses can now be interrogated as deep learning enables analysis of increasing amounts of data generated by new technologies. For example, the effects of cellular heterogeneity on basic biology and disease etiology can now be explored by single-cell RNA-seq and high-throughput fluorescence-based imaging, techniques we discuss below that will benefit immensely from deep learning approaches.

### Splicing

Pre-mRNA transcripts can be spliced into different isoforms by retaining or skipping subsets of exons or including parts of introns, creating enormous spatiotemporal flexibility to generate multiple distinct proteins from a single gene. This remarkable complexity can lend itself to defects that underlie many diseases. For instance, splicing mutations in the lamin A (*LMNA*) gene can lead to specific variants of dilated cardiomyopathy and limb girdle muscular dystrophy [185]. A recent study found that quantitative trait loci that affect splicing in lymphoblastoid cell lines are enriched within risk loci for schizophrenia, multiple sclerosis, and other immune diseases, implicating mis-splicing as a more widespread feature of human pathologies than previously thought [186]. Therapeutic strategies that aim to modulate splicing are also currently being considered for disorders such as Duchenne muscular dystrophy and spinal muscular atrophy [185].

Sequencing studies routinely return thousands of unannotated variants, but which cause functional changes in splicing and how are those changes manifested? Prediction of a “splicing code” has been a goal of the field for the past decade. Initial machine learning approaches used a naïve Bayes model and a 2-layer Bayesian neural network with thousands of hand-derived sequence-based features to predict the probability of exon skipping [187, 188]. With the advent of deep learning, more complex models provided better predictive accuracy [189, 190]. Importantly, these new approaches can take in multiple kinds of epigenomic measurements as well as tissue identity and RNA binding partners of splicing factors. Deep learning is critical in furthering these kinds of integrative studies where different data types and inputs interact in unpredictable (often nonlinear) ways to create higher-order features. Moreover, as in gene expression network analysis, interrogating the hidden nodes within neural networks could potentially illuminate important aspects of splicing behavior. For instance, tissue-specific splicing mechanisms could be inferred by training networks on splicing data from different tissues, then searching for common versus distinctive hidden nodes, a technique employed by Qin et al. for tissue-specific transcription factor (TF) binding predictions [191].

A parallel effort has been to use more data with simpler models. An exhaustive study using readouts of splicing for millions of synthetic intronic sequences uncovered motifs that influence the strength of alternative splice sites [192]. The authors built a simple linear model using hexamer motif frequencies that successfully generalized to exon skipping. In a limited analysis using single nucleotide polymorphisms (SNPs) from three genes, it predicted exon skipping with three times the accuracy of an existing deep learning-based framework [189]. This case is instructive in that clever sources of data, not just more descriptive models, are still critical.

We already understand how mis-splicing of a single gene can cause diseases such as limb girdle muscular dystrophy. The challenge now is to uncover how genome-wide alternative splicing underlies complex, non-Mendelian diseases such as autism, schizophrenia, Type 1 diabetes, and multiple sclerosis [193]. As a proof of concept, Xiong et al. [189] sequenced five autism spectrum disorder and 12 control samples, each with an average of 42,000 rare variants, and identified mis-splicing in 19 genes with neural functions. Such methods may one day enable scientists and clinicians to rapidly profile thousands of unannotated variants for functional effects on splicing and nominate candidates for further investigation. Moreover, these nonlinear algorithms can deconvolve the effects of multiple variants on a single splice event without the need to perform combinatorial *in vitro* experiments. The ultimate goal is to predict an individual’s tissue-specific, exon-specific splicing patterns from their genome sequence and other measurements to enable a new branch of precision diagnostics that also stratifies patients and suggests targeted therapies to correct splicing defects. However, to achieve this we expect that methods to interpret the “black box” of deep neural networks and integrate diverse data sources will be required.

### Transcription factors

Transcription factors are proteins that bind regulatory DNA in a sequence-specific manner to modulate the activation and repression of gene transcription. High-throughput *in vitro* experimental assays that quantitatively measure the binding specificity of a TF to a large library of short oligonucleotides [194] provide rich datasets to model the naked DNA sequence affinity of individual TFs in isolation. However, *in vivo* TF binding is affected by a variety of other factors beyond sequence affinity, such as competition and cooperation with other TFs, TF concentration, and chromatin state (chemical modifications to DNA and other packaging proteins that DNA is wrapped around) [194]. TFs can thus exhibit highly variable binding landscapes across the same genomic DNA sequence across diverse cell types and states. Several experimental approaches such as chromatin immunoprecipitation followed by sequencing (ChIP-seq) have been developed to profile *in vivo* binding maps of TFs [194]. Large reference compendia of ChIP-seq data are now freely available for a large collection of TFs in a small number of reference cell states in humans and a few other model organisms [195]. Due to fundamental material and cost constraints, it is infeasible to perform these experiments for all TFs in every possible cellular state and species. Hence, predictive computational models of TF binding are essential to understand gene regulation in diverse cellular contexts.

Several machine learning approaches have been developed to learn generative and discriminative models of TF binding from *in vitro* and *in vivo* TF binding datasets that associate collections of synthetic DNA sequences or genomic DNA sequences to binary labels (bound/unbound) or continuous measures of binding. The most common class of TF binding models in the literature are those that only model the DNA sequence affinity of TFs from *in vitro* and *in vivo* binding data. The earliest models were based on deriving simple, compact, interpretable sequence motif representations such as position weight matrices (PWMs) and other biophysically inspired models [196–198]. These models were outperformed by general k-mer based models including support vector machines (SVMs) with string kernels [199, 200].

In 2015, Alipanahi et al. developed DeepBind, the first CNN to classify bound DNA sequences based on *in vitro* and *in vivo* assays against random DNA sequences matched for dinucleotide sequence composition [201]. The convolutional layers learn pattern detectors reminiscent of PWMs from a one-hot encoding of the raw input DNA sequences. DeepBind outperformed several state-of-the-art methods from the DREAM5 *in vitro* TF-DNA motif recognition challenge [198]. Although DeepBind was also applied to RNA-binding proteins, in general RNA binding is a separate problem [202] and accurate models will need to account for RNA secondary structure. Following DeepBind, several optimized convolutional and recurrent neural network architectures as well as novel hybrid approaches that combine kernel methods with neural networks have been proposed that further improve performance [203–206]. Specialized layers and regularizers have also been proposed to reduce parameters and learn more robust models by taking advantage of specific properties of DNA sequences such as their reverse complement equivalence [207, 208].

While most of these methods learn independent models for different TFs, *in vivo* multiple TFs compete or cooperate to occupy DNA binding sites, resulting in complex combinatorial co-binding landscapes. To take advantage of this shared structure in *in vivo* TF binding data, multi-task neural network architectures have been developed that explicitly share parameters across models for multiple TFs [206,209,210]. Some of these multi-task models train and evaluate classification performance relative to an unbound background set of regulatory DNA sequences sampled from the genome rather than using synthetic background sequences with matched dinucleotide composition.

The above-mentioned TF binding prediction models that use only DNA sequences as inputs have a fundamental limitation. Because the DNA sequence of a genome is the same across different cell types and states, a sequence-only model of TF binding cannot predict different *in vivo* TF binding landscapes in new cell types not used during training. One approach for generalizing TF binding predictions to new cell types is to learn models that integrate DNA sequence inputs with other cell-type-specific data modalities that modulate *in vivo* TF binding such as surrogate measures of TF concentration (e.g. TF gene expression) and chromatin state. Arvey et al. showed that combining the predictions of SVMs trained on DNA sequence inputs and cell-type specific DNase-seq data, which measures genome-wide chromatin accessibility, improved *in vivo* TF binding prediction within and across cell types [211]. Several “footprinting” based methods have also been developed that learn to discriminate bound from unbound instances of known canonical motifs of a target TF based on high-resolution footprint patterns of chromatin accessibility that are specific to the target TF [212]. However, the genome-wide predictive performance of these methods in new cell types and states has not been evaluated.

Recently, a community challenge known as the “ENCODE-DREAM *in vivo* TF Binding Site Prediction Challenge” was introduced to systematically evaluate genome-wide performance of methods that can predict TF binding across cell states by integrating DNA sequence and *in vitro* DNA shape with cell-type-specific chromatin accessibility and gene expression [213]. A deep learning model called FactorNet was amongst the top three performing methods in the challenge [214]. FactorNet uses a multi-modal hybrid convolutional and recurrent architecture that integrates DNA sequence with chromatin accessibility profiles, gene expression, and evolutionary conservation of sequence. It is worth noting that FactorNet was slightly outperformed by an approach that does not use neural networks [215]. This top ranking approach uses an extensive set of curated features in a weighted variant of a discriminative maximum conditional likelihood model in combination with a novel iterative training strategy and model stacking. There appears to be significant room for improvement because none of the current approaches for cross cell type prediction explicitly account for the fact that TFs can co-bind with distinct co-factors in different cell states. In such cases, sequence features that are predictive of TF binding in one cell state may be detrimental to predicting binding in another.

Singh et al. developed transfer string kernels for SVMs for cross-context TF binding [216]. Domain adaptation methods that allow training neural networks which are transferable between differing training and test set distributions of sequence features could be a promising avenue going forward [217, 218]. These approaches may also be useful for transferring TF binding models across species.

Another class of imputation-based cross cell type *in vivo* TF binding prediction methods leverage the strong correlation between combinatorial binding landscapes of multiple TFs. Given a partially complete panel of binding profiles of multiple TFs in multiple cell types, a deep learning method called TFImpute learns to predict the missing binding profile of a target TF in some target cell type in the panel based on the binding profiles of other TFs in the target cell type and the binding profile of the target TF in other cell types in the panel [191]. However, TFImpute cannot generalize predictions beyond the training panel of cell types and requires TF binding profiles of related TFs.

It is worth noting that TF binding prediction methods in the literature based on neural networks and other machine learning approaches choose to sample the set of bound and unbound sequences in a variety of different ways. These choices and the choice of performance evaluation measures significantly confound systematic comparison of model performance (see Discussion).

Several methods have also been developed to interpret neural network models of TF binding. Alipanahi et al. visualize convolutional filters to obtain insights into the sequence preferences of TFs [201]. They also introduced *in silico* mutation maps for identifying important predictive nucleotides in input DNA sequences by exhaustively forward propagating perturbations to individual nucleotides to record the corresponding change in output prediction. Shrikumar et al. [219] proposed efficient backpropagation based approaches to simultaneously score the contribution of all nucleotides in an input DNA sequence to an output prediction. Lanchantin et al. [204] developed tools to visualize TF motifs learned from TF binding site classification tasks. These and other general interpretation techniques (see Discussion) will be critical to improve our understanding of the biologically meaningful patterns learned by deep learning models of TF binding.

## Promoters and enhancers

### From TF binding to promoters and enhancers

Multiple TFs act in concert to coordinate changes in gene regulation at the genomic regions known as promoters and enhancers. Each gene has an upstream promoter, essential for initiating that gene’s transcription. The gene may also interact with multiple enhancers, which can amplify transcription in particular cellular contexts. These contexts include different cell types in development or environmental stresses.

Promoters and enhancers provide a nexus where clusters of TFs and binding sites mediate downstream gene regulation, starting with transcription. The gold standard to identify an active promoter or enhancer requires demonstrating its ability to affect transcription or other downstream gene products. Even extensive biochemical TF binding data has thus far proven insufficient on its own to accurately and comprehensively locate promoters and enhancers. We lack sufficient understanding of these elements to derive a mechanistic “promoter code” or “enhancer code”. But extensive labeled data on promoters and enhancers lends itself to probabilistic classification. The complex interplay of TFs and chromatin leading to the emergent properties of promoter and enhancer activity seems particularly apt for representation by deep neural networks.

### Promoters

Despite decades of work, computational identification of promoters remains a stubborn problem [220]. Researchers have used neural networks for promoter recognition as early as 1996 [221]. Recently, a CNN recognized promoter sequences with sensitivity and specificity exceeding 90% [222]. Most activity in computational prediction of regulatory regions, however, has moved to enhancer identification. Because one can identify promoters with straightforward biochemical assays [223, 224], the direct rewards of promoter prediction alone have decreased. But the reliable ground truth provided by these assays makes promoter identification an appealing test bed for deep learning approaches that can also identify enhancers.

### Enhancers

Recognizing enhancers presents additional challenges. Enhancers may be up to 1,000,000 bp away from the affected promoter, and even within introns of other genes [225]. Enhancers do not necessarily operate on the nearest gene and may affect multiple genes. Their activity is frequently tissue-or context-specific. No biochemical assay can reliably identify all enhancers. Distinguishing them from other regulatory elements remains difficult, and some believe the distinction somewhat artificial [226]. While these factors make the enhancer identification problem more difficult, they also make a solution more valuable.

Several neural network approaches yielded promising results in enhancer prediction. Both Basset [227] and DeepEnhancer [228] used CNNs to predict enhancers. DECRES used a feed-forward neural network [229] to distinguish between different kinds of regulatory elements, such as active enhancers, and promoters. DECRES had difficulty distinguishing between inactive enhancers and promoters. They also investigated the power of sequence features to drive classification, finding that beyond CpG islands, few were useful.

Comparing the performance of enhancer prediction methods illustrates the problems in using metrics created with different benchmarking procedures. Both the Basset and DeepEnhancer studies include comparisons to a baseline SVM approach, gkm-SVM [200]. The Basset study reports gkm-SVM attains a mean area under the precision-recall curve (AUPR) of 0.322 over 164 cell types [227]. The DeepEnhancer study reports for gkm-SVM a dramatically different AUPR of 0.899 on nine cell types [228]. This large difference means it’s impossible to directly compare the performance of Basset and DeepEnhancer based solely on their reported metrics. DECRES used a different set of metrics altogether. To drive further progress in enhancer identification, we must develop a common and comparable benchmarking procedure (see Discussion).

### Promoter-enhancer interactions

In addition to the location of enhancers, identifying enhancer-promoter interactions in three-dimensional space will provide critical knowledge for understanding transcriptional regulation. SPEID used a CNN to predict these interactions with only sequence and the location of putative enhancers and promoters along a one-dimensional chromosome [230]. It compared well to other methods using a full complement of biochemical data from ChIP-seq and other epigenomic methods. Of course, the putative enhancers and promoters used were themselves derived from epigenomic methods. But one could easily replace them with the output of one of the enhancer or promoter prediction methods above.

### Micro-RNA binding

Prediction of microRNAs (miRNAs) and miRNA targets is of great interest, as they are critical components of gene regulatory networks and are often conserved across great evolutionary distance [231, 232]. While many machine learning algorithms have been applied to these tasks, they currently require extensive feature selection and optimization. For instance, one of the most widely adopted tools for miRNA target prediction, TargetScan, trained multiple linear regression models on 14 hand-curated features including structural accessibility of the target site on the mRNA, the degree of site conservation, and predicted thermodynamic stability of the miRNA-mRNA complex [233]. Some of these features, including structural accessibility, are imperfect or empirically derived. In addition, current algorithms suffer from low specificity [234].

As in other applications, deep learning promises to achieve equal or better performance in predictive tasks by automatically engineering complex features to minimize an objective function. Two recently published tools use different recurrent neural network-based architectures to perform miRNA and target prediction with solely sequence data as input [234, 235]. Though the results are preliminary and still based on a validation set rather than a completely independent test set, they were able to predict microRNA target sites with higher specificity and sensitivity than TargetScan. Excitingly, these tools seem to show that RNNs can accurately align sequences and predict bulges, mismatches, and wobble base pairing without requiring the user to input secondary structure predictions or thermodynamic calculations. Further incremental advances in deep learning for miRNA and target prediction will likely be sufficient to meet the current needs of systems biologists and other researchers who use prediction tools mainly to nominate candidates that are then tested experimentally.

### Protein secondary and tertiary structure

Proteins play fundamental roles in almost all biological processes, and understanding their structure is critical for basic biology and drug development. UniProt currently has about 94 million protein sequences, yet fewer than 100,000 proteins across all species have experimentally-solved structures in Protein Data Bank (PDB). As a result, computational structure prediction is essential for a majority of proteins. However, this is very challenging, especially when similar solved structures, called templates, are not available in PDB. Over the past several decades, many computational methods have been developed to predict aspects of protein structure such as secondary structure, torsion angles, solvent accessibility, inter-residue contact maps, disorder regions, and side-chain packing. In recent years, multiple deep learning architectures have been applied, including deep belief networks, LSTMs, CNNs, and deep convolutional neural fields (DeepCNFs) [30, 236].

Here we focus on deep learning methods for two representative sub-problems: secondary structure prediction and contact map prediction. Secondary structure refers to local conformation of a sequence segment, while a contact map contains information on all residue-residue contacts. Secondary structure prediction is a basic problem and an almost essential module of any protein structure prediction package. Contact prediction is much more challenging than secondary structure prediction, but it has a much larger impact on tertiary structure prediction. In recent years, the accuracy of contact prediction has greatly improved [28,237–239].

One can represent protein secondary structure with three different states (alpha helix, beta strand, and loop regions) or eight finer-grained states. Accuracy of a three-state prediction is called Q3, and accuracy of an 8-state prediction is called Q8. Several groups [29,240,241] applied deep learning to protein secondary structure prediction but were unable to achieve significant improvement over the *de facto* standard method PSIPRED [242], which uses two shallow feedforward neural networks. In 2014, Zhou and Troyanskaya demonstrated that they could improve Q8 accuracy by using a deep supervised and convolutional generative stochastic network [243]. In 2016 Wang et al. developed a DeepCNF model that improved Q3 and Q8 accuracy as well as prediction of solvent accessibility and disorder regions [30, 236]. DeepCNF achieved a higher Q3 accuracy than the standard maintained by PSIPRED for more than 10 years. This improvement may be mainly due to the ability of convolutional neural fields to capture long-range sequential information, which is important for beta strand prediction. Nevertheless, the improvements in secondary structure prediction from DeepCNF are unlikely to result in a commensurate improvement in tertiary structure prediction since secondary structure mainly reflects coarse-grained local conformation of a protein structure.

Protein contact prediction and contact-assisted folding (i.e. folding proteins using predicted contacts as restraints) represents a promising new direction for *ab initio* folding of proteins without good templates in PDB. Co-evolution analysis is effective for proteins with a very large number (>1000) of sequence homologs [239], but fares poorly for proteins without many sequence homologs. By combining co-evolution information with a few other protein features, shallow neural network methods such as MetaPSICOV [237] and CoinDCA-NN [244] have shown some advantage over pure co-evolution analysis for proteins with few sequence homologs, but their accuracy is still far from satisfactory. In recent years, deeper architectures have been explored for contact prediction, such as CMAPpro [245], DNCON [246] and PConsC [247]. However, blindly tested in the well-known CASP competitions, these methods did not show any advantage over MetaPSICOV [237].

Recently, Wang et al. proposed the deep learning method RaptorX-Contact [28], which significantly improves contact prediction over MetaPSICOV and pure co-evolution methods, especially for proteins without many sequence homologs. It employs a network architecture formed by one 1D residual neural network and one 2D residual neural network. Blindly tested in the latest CASP competition (i.e. CASP12 [248]), RaptorX-Contact ranked first in F₁ score on free-modeling targets as well as the whole set of targets. In CAMEO (which can be interpreted as a fully-automated CASP) [249], its predicted contacts were also able to fold proteins with a novel fold and only 65– 330 sequence homologs. This technique also worked well on membrane proteins even when trained on non-membrane proteins [250]. RaptorX-Contact performed better mainly due to introduction of residual neural networks and exploitation of contact occurrence patterns by simultaneously predicting all the contacts in a single protein.

Taken together, *ab initio* folding is becoming much easier with the advent of direct evolutionary coupling analysis and deep learning techniques. We expect further improvements in contact prediction for proteins with fewer than 1000 homologs by studying new deep network architectures. However, it is unclear if there is an effective way to use deep learning to improve prediction for proteins with few or no sequence homologs. Finally, the deep learning methods summarized above also apply to interfacial contact prediction for protein complexes but may be less effective since on average protein complexes have fewer sequence homologs.

### Structure determination and cryo-electron microscopy

Complementing computational prediction approaches, cryo-electron microscopy (cryo-EM) allows near-atomic resolution determination of protein models by comparing individual electron micrographs [251].

Detailed structures require tens of thousands of protein images [252]. Technological development has increased the throughput of image capture. New hardware, such as direct electron detectors, has made large-scale image production practical, while new software has focused on rapid, automated image processing.

Some components of cryo-EM image processing remain difficult to automate. For instance, in particle picking, micrographs are scanned to identify individual molecular images that will be used in structure refinement. In typical applications, hundreds of thousands of particles are necessary to determine a structure to near atomic resolution, making manual selection impractical [252]. Typical selection approaches are semi-supervised; a user will select several particles manually, and these selections will be used to train a classifier [253, 254]. Now CNNs are being used to select particles in tools like DeepPicker [255] and DeepEM [256]. In addition to addressing shortcomings from manual selection, such as selection bias and poor discrimination of low-contrast images, these approaches also provide a means of full automation. DeepPicker can be trained by reference particles from other experiments with structurally unrelated macromolecules, allowing for fully automated application to new samples.

Downstream of particle picking, deep learning is being applied to other aspects of cryo-EM image processing. Statistical manifold learning has been implemented in the software package ROME to classify selected particles and elucidate the different conformations of the subject molecule necessary for accurate 3D structures [257]. These recent tools highlight the general applicability of deep learning approaches for image processing to increase the throughput of high-resolution cryo-EM.

### Protein-protein interactions

Protein-protein interactions (PPIs) are highly specific and non-accidental physical contacts between proteins, which occur for purposes other than generic protein production or degradation [258]. Abundant interaction data have been generated in-part thanks to advances in high-throughput screening methods, such as yeast two-hybrid and affinity-purification with mass spectrometry. However, because many PPIs are transient or dependent on biological context, high-throughput methods can fail to capture a number of interactions. The imperfections and costs associated with many experimental PPI screening methods have motivated an interest in high-throughput computational prediction.

Many machine learning approaches to PPI have focused on text mining the literature [259, 260], but these approaches can fail to capture context-specific interactions, motivating *de novo* PPI prediction. Early *de novo* prediction approaches used a variety of statistical and machine learning tools on structural and sequential data, sometimes with reference to the existing body of protein structure knowledge. In the context of PPIs—as in other domains—deep learning shows promise both for exceeding current predictive performance and for circumventing limitations from which other approaches suffer.

One of the key difficulties in applying deep learning techniques to binding prediction is the task of representing peptide and protein sequences in a meaningful way. DeepPPI [261] made PPI predictions from a set of sequence and composition protein descriptors using a two-stage deep neural network that trained two subnetworks for each protein and combined them into a single network. Sun et al. [262] applied autocovariances, a coding scheme that returns uniform-size vectors describing the covariance between physicochemical properties of the protein sequence at various positions. Wang et al. [263] used deep learning as an intermediate step in PPI prediction. They examined 70 amino acid protein sequences from each of which they extracted 1260 features. A stacked sparse autoencoder with two hidden layers was then used to reduce feature dimensions and noisiness before a novel type of classification vector machine made PPI predictions.

Beyond predicting whether or not two proteins interact, Du et al. [264] employed a deep learning approach to predict the residue contacts between two interacting proteins. Using features that describe how similar a protein’s residue is relative to similar proteins at the same position, the authors extracted uniform-length features for each residue in the protein sequence. A stacked autoencoder took two such vectors as input for the prediction of contact between two residues. The authors evaluated the performance of this method with several classifiers and showed that a deep neural network classifier paired with the stacked autoencoder significantly exceeded classical machine learning accuracy.

Because many studies used predefined higher—level features, one of the benefits of deep learning — automatic feature extraction—is not fully leveraged. More work is needed to determine the best ways to represent raw protein sequence information so that the full benefits of deep learning as an automatic feature extractor can be realized.

### MHC-peptide binding

An important type of PPI involves the immune system’s ability to recognize the body’s own cells. The major histocompatibility complex (MHC) plays a key role in regulating this process by binding antigens and displaying them on the cell surface to be recognized by T cells. Due to its importance in immunity and immune response, peptide-MHC binding prediction is a useful problem in computational biology, and one that must account for the allelic diversity in MHC-encoding gene region.

Shallow, feed-forward neural networks are competitive methods and have made progress toward pan-allele and pan-length peptide representations. Sequence alignment techniques are useful for representing variable-length peptides as uniform-length features [265, 266]. For pan-allelic prediction, NetMHCpan [267, 268] used a pseudo-sequence representation of the MHC class I molecule, which included only polymorphic peptide contact residues. The sequences of the peptide and MHC were then represented using both sparse vector encoding and Blosum encoding, in which amino acids are encoded by matrix score vectors. A comparable method to the NetMHC tools is MHCflurry [269], a method which shows superior performance on peptides of lengths other than nine. MHCflurry adds placeholder amino acids to transform variable-length peptides to length 15 peptides. In training the MHCflurry feed-forward neural network [270], the authors imputed missing MHC-peptide binding affinities using a Gibbs sampling method, showing that imputation improves performance for data-sets with roughly 100 or fewer training examples. MHCflurry’s imputation method increases its performance on poorly characterized alleles, making it competitive with NetMHCpan for this task. Kuksa et al. [271] developed a shallow, higher-order neural network (HONN) comprised of both mean and covariance hidden units to capture some of the higher-order dependencies between amino acid locations. Pretraining this HONN with a semi-restricted Boltzmann machine, the authors found that the performance of the HONN exceeded that of a simple deep neural network, as well as that of NetMHC.

Deep learning’s unique flexibility was recently leveraged by Bhattacharya et al. [272], who used a gated RNN method called MHCnuggets to overcome the difficulty of multiple length peptides. Under this framework, they used smoothed sparse encoding to represent amino acids individually. Because MHCnuggets had to be trained for every MHC allele, performance was far better for alleles with abundant, balanced training data. Vang et al. [273] developed HLA-CNN, a method which maps amino acids onto a 15-dimensional vector space based on their context relation to other amino acids before making predictions with a CNN. In a comparison of several current methods, Bhattacharya et al. found that the top methods—NetMHC, NetMHCpan, MHCflurry, and MHCnuggets—showed comparable performance, but large differences in speed. Convolutional neural networks (in this case, HLA-CNN) showed comparatively poor performance, while shallow and recurrent neural networks performed the best. They found that MHCnuggets—the recurrent neural network—was by far the fastest-training among the top performing methods.

### PPI networks and graph analysis

Because interacting proteins are more likely to share a similar function, the connectivity of a PPI network itself can be a valuable information source for the prediction of protein function [274]. To incorporate higher-order network information, it is necessary to find a lower-level embedding of network structure that preserves this higher-order structure. Rather than use hand-crafted network features, deep learning shows promise for the automatic discovery of predictive features within networks. For example, Navlakha [275] showed that a deep autoencoder was able to compress a graph to 40% of its original size, while being able to reconstruct 93% of the original graph’s edges, improving upon standard dimension reduction methods. To achieve this, each graph was represented as an adjacency matrix with rows sorted in descending node degree order, then flattened into a vector and given as input to the autoencoder. While the activity of some hidden layers correlated with several popular hand-crafted network features such as k-core size and graph density, this work showed that deep learning can effectively reduce graph dimensionality while retaining much of its structural information.

An important challenge in PPI network prediction is the task of combining different networks and types of networks. Gligorijevic et al. [276] developed a multimodal deep autoencoder, deepNF, to find a feature representation common among several different PPI networks. This common lower-level representation allows for the combination of various PPI data sources towards a single predictive task. An SVM classifier trained on the compressed features from the middle layer of the autoencoder outperformed previous methods in predicting protein function.

Hamilton et al. addressed the issue of large, heterogeneous, and changing networks with an inductive approach called GraphSAGE [277]. By finding node embeddings through learned aggregator functions that describe the node and its neighbors in the network, the GraphSAGE approach allows for the generalization of the model to new graphs. In a classification task for the prediction of protein function, Chen and Zhu [278] optimized this approach and enhanced the graph convolutional network with a preprocessing step that uses an approximation to the dropout operation. This preprocessing effectively reduces the number of graph convolutional layers and it significantly improves both training time and prediction accuracy.

### Morphological phenotypes

A field poised for dramatic revolution by deep learning is bioimage analysis. Thus far, the primary use of deep learning for biological images has been for segmentation—that is, for the identification of biologically relevant structures in images such as nuclei, infected cells, or vasculature—in fluorescence or even brightfield channels [279]. Once so-called regions of interest have been identified, it is often straightforward to measure biological properties of interest, such as fluorescence intensities, textures, and sizes. Given the dramatic successes of deep learning in biological imaging, we simply refer to articles that review recent advancements [16,279,280]. For deep learning to become commonplace for biological image segmentation, we need user-friendly tools.

We anticipate an additional paradigm shift in bioimaging that will be brought about by deep learning: what if images of biological samples, from simple cell cultures to three-dimensional organoids and tissue samples, could be mined for much more extensive biologically meaningful information than is currently standard? For example, a recent study demonstrated the ability to predict lineage fate in hematopoietic cells up to three generations in advance of differentiation [281]. In biomedical research, most often biologists decide in advance what feature to measure in images from their assay system. Although classical methods of segmentation and feature extraction can produce hundreds of metrics per cell in an image, deep learning is unconstrained by human intuition and can in theory extract more subtle features through its hidden nodes. Already, there is evidence deep learning can surpass the efficacy of classical methods [282], even using generic deep convolutional networks trained on natural images [283], known as transfer learning. Recent work by Johnson et al. [284] demonstrated how the use of a conditional adversarial autoencoder allows for a probabilistic interpretation of cell and nuclear morphology and structure localization from fluorescence images. The proposed model is able to generalize well to a wide range of subcellular localizations. The generative nature of the model allows it to produce high-quality synthetic images predicting localization of subcellular structures by directly modeling the localization of fluorescent labels. Notably, this approach reduces the modeling time by omitting the subcellular structure segmentation step.

The impact of further improvements on biomedicine could be enormous. Comparing cell population morphologies using conventional methods of segmentation and feature extraction has already proven useful for functionally annotating genes and alleles, identifying the cellular target of small molecules, and identifying disease-specific phenotypes suitable for drug screening [285–287]. Deep learning would bring to these new kinds of experiments—known as image-based profiling or morphological profiling—a higher degree of accuracy, stemming from the freedom from human-tuned feature extraction strategies.

### Single-cell data

Single-cell methods are generating excitement as biologists characterize the vast heterogeneity within unicellular species and between cells of the same tissue type in the same organism [288]. For instance, tumor cells and neurons can both harbor extensive somatic variation [289].

Understanding single-cell diversity in all its dimensions—genetic, epigenomic, transcriptomic, proteomic, morphologic, and metabolic—is key if treatments are to be targeted not only to a specific individual, but also to specific pathological subsets of cells. Single-cell methods also promise to uncover a wealth of new biological knowledge. A sufficiently large population of single cells will have enough representative “snapshots” to recreate timelines of dynamic biological processes. If tracking processes over time is not the limiting factor, single-cell techniques can provide maximal resolution compared to averaging across all cells in bulk tissue, enabling the study of transcriptional bursting with single-cell fluorescence *in situ* hybridization or the heterogeneity of epigenomic patterns with single-cell Hi-C or ATAC-seq [290, 291]. Joint profiling of single-cell epigenomic and transcriptional states provides unprecedented views of regulatory processes [292].

However, large challenges exist in studying single cells. Relatively few cells can be assayed at once using current droplet, imaging, or microwell technologies, and low-abundance molecules or modifications may not be detected by chance due to a phenomenon known as dropout, not to be confused with the dropout layer of deep learning. To solve this problem, Angermueller et al. [293] trained a neural network to predict the presence or absence of methylation of a specific CpG site in single cells based on surrounding methylation signal and underlying DNA sequence, achieving several percentage points of improvement compared to random forests or deep networks trained only on CpG or sequence information. Similar deep learning methods have been applied to impute low-resolution ChIP-seq signal from bulk tissue with great success, and they could easily be adapted to single-cell data [191, 294]. Deep learning has also been useful for dealing with batch effects [295].

Examining populations of single cells can reveal biologically meaningful subsets of cells as well as their underlying gene regulatory networks [296]. Unfortunately, machine learning methods generally struggle with imbalanced data—when there are many more examples of class 1 than class 2—because prediction accuracy is usually evaluated over the entire dataset. To tackle this challenge, Arvaniti et al. [297] classified healthy and cancer cells expressing 25 markers by using the most discriminative filters from a CNN trained on the data as a linear classifier. They achieved impressive performance, even for cell types where the subset percentage ranged from 0.1 to 1%, significantly outperforming logistic regression and distance-based outlier detection methods. However, they did not benchmark against random forests, which tend to work better for imbalanced data, and their data was relatively low dimensional.

Neural networks can also learn low-dimensional representations of single-cell gene expression data for visualization, clustering, and other tasks. Both scvis [298] and scVI [299] are unsupervised approaches based on VAEs. Whereas scvis primarily focuses on single-cell visualization as a replacement for t-Distributed Stochastic Neighbor Embedding [300], the scVI model accounts for zero-inflated expression distributions and can impute zero values that are due to technical effects. Beyond VAEs, Lin et al. developed a supervised model to predict cell type [301]. Similar to transfer learning approaches for microscopy images [283], they demonstrated that the hidden layer representations were informative in general and could be used to identify cellular subpopulations or match new cells to known cell types. The supervised neural network’s representation was better overall at retrieving cell types than alternatives, but all methods struggled to recover certain cell types such as hematopoietic stem cells and inner cell mass cells. As the Human Cell Atlas [302] and related efforts generate more single-cell expression data, there will be opportunities to assess how well these low-dimensional representations generalize to new cell types as well as abundant training data to learn broadly-applicable representations.

The sheer quantity of omic information that can be obtained from each cell, as well as the number of cells in each dataset, uniquely position single-cell data to benefit from deep learning. In the future, lineage tracing could be revolutionized by using autoencoders to reduce the feature space of transcriptomic or variant data followed by algorithms to learn optimal cell differentiation trajectories [303] or by feeding cell morphology and movement into neural networks [281]. Reinforcement learning algorithms [304] could be trained on the evolutionary dynamics of cancer cells or bacterial cells undergoing selection pressure and reveal whether patterns of adaptation are random or deterministic, allowing us to develop therapeutic strategies that forestall resistance. We are excited to see the creative applications of deep learning to single-cell biology that emerge over the next few years.

### Metagenomics

Metagenomics, which refers to the study of genetic material—16S rRNA or whole-genome shotgun DNA—from microbial communities, has revolutionized the study of micro-scale ecosystems within and around us. In recent years, machine learning has proved to be a powerful tool for metagenomic analysis. 16S rRNA has long been used to deconvolve mixtures of microbial genomes, yet this ignores more than 99% of the genomic content. Subsequent tools aimed to classify 300–3000 bp reads from complex mixtures of microbial genomes based on tetranucleotide frequencies, which differ across organisms [305], using supervised [306, 307] or unsupervised methods [308]. Then, researchers began to use techniques that could estimate relative abundances from an entire sample faster than classifying individual reads [309–312]. There is also great interest in identifying and annotating sequence reads [313, 314]. However, the focus on taxonomic and functional annotation is just the first step. Several groups have proposed methods to determine host or environment phenotypes from the organisms that are identified [315–318] or overall sequence composition [319]. Also, researchers have looked into how feature selection can improve classification [318, 320], and techniques have been proposed that are classifier-independent [321, 322].

Most neural networks are used for phylogenetic classification or functional annotation from sequence data where there is ample data for training. Neural networks have been applied successfully to gene annotation (e.g. Orphelia [323] and FragGeneScan [324]). Representations (similar to Word2Vec [102] in natural language processing) for protein family classification have been introduced and classified with a skip-gram neural network [325]. Recurrent neural networks show good performance for homology and protein family identification [326, 327].

One of the first techniques of *de novo* genome binning used self-organizing maps, a type of neural network [308]. Essinger et al. [328] used Adaptive Resonance Theory to cluster similar genomic fragments and showed that it had better performance than k-means. However, other methods based on interpolated Markov models [329] have performed better than these early genome binners. Neural networks can be slow and therefore have had limited use for reference-based taxonomic classification, with TAC-ELM [330] being the only neural network-based algorithm to taxonomically classify massive amounts of metagenomic data. An initial study successfully applied neural networks to taxonomic classification of 16S rRNA genes, with convolutional networks providing about 10% accuracy genus-level improvement over RNNs and random forests [331]. However, this study evaluated only 3000 sequences.

Neural network uses for classifying phenotype from microbial composition are just beginning. A simple multi-layer perceptron (MLP) was able to classify wound severity from microbial species present in the wound [332]. Recently, Ditzler et al. associated soil samples with pH level using MLPs, DBNs, and RNNs [333]. Besides classifying samples appropriately, internal phylogenetic tree nodes inferred by the networks represented features for low and high pH. Thus, hidden nodes might provide biological insight as well as new features for future metagenomic sample comparison. Also, an initial study has shown promise of these networks for diagnosing disease [334].

Challenges remain in applying deep neural networks to metagenomics problems. They are not yet ideal for phenotype classification because most studies contain tens of samples and hundreds or thousands of features (species). Such underdetermined, or ill-conditioned, problems are still a challenge for deep neural networks that require many training examples. Also, due to convergence issues [335], taxonomic classification of reads from whole genome sequencing seems out of reach at the moment for deep neural networks. There are only thousands of full-sequenced genomes as compared to hundreds of thousands of 16S rRNA sequences available for training.

However, because RNNs have been applied to base calls for the Oxford Nanopore long-read sequencer with some success [336] (discussed below), one day the entire pipeline, from denoising to functional classification, may be combined into one step using powerful LSTMs [337]. For example, metagenomic assembly usually requires binning then assembly, but could deep neural nets accomplish both tasks in one network? We believe the greatest potential in deep learning is to learn the complete characteristics of a metagenomic sample in one complex network.

### Sequencing and variant calling

While we have so far primarily discussed the role of deep learning in analyzing genomic data, deep learning can also substantially improve our ability to obtain the genomic data itself. We discuss two specific challenges: calling SNPs and indels (insertions and deletions) with high specificity and sensitivity and improving the accuracy of new types of data such as nanopore sequencing. These two tasks are critical for studying rare variation, allele-specific transcription and translation, and splice site mutations. In the clinical realm, sequencing of rare tumor clones and other genetic diseases will require accurate calling of SNPs and indels.

Current methods achieve relatively high (>99%) precision at 90% recall for SNPs and indel calls from Illumina short-read data [338], yet this leaves a large number of potentially clinically-important remaining false positives and false negatives. These methods have so far relied on experts to build probabilistic models that reliably separate signal from noise. However, this process is time consuming and fundamentally limited by how well we understand and can model the factors that contribute to noise. Recently, two groups have applied deep learning to construct data-driven unbiased noise models. One of these models, DeepVariant, leverages Inception, a neural network trained for image classification by Google Brain, by encoding reads around a candidate SNP as a 221x100 bitmap image, where each column is a nucleotide and each row is a read from the sample library [338]. The top 5 rows represent the reference, and the bottom 95 rows represent randomly sampled reads that overlap the candidate variant. Each RGBA (red/green/blue/alpha) image pixel encodes the base (A, C, G, T) as a different red value, quality score as a green value, strand as a blue value, and variation from the reference as the alpha value. The neural network outputs genotype probabilities for each candidate variant. They were able to achieve better performance than GATK [339], a leading genotype caller, even when GATK was given information about population variation for each candidate variant. Another method, still in its infancy, hand-developed 62 features for each candidate variant and fed these vectors into a fully connected deep neural network [340]. Unfortunately, this feature set required at least 15 iterations of software development to fine-tune, which suggests that these models may not generalize.

Variant calling will benefit more from optimizing neural network architectures than from developing features by hand. An interesting and informative next step would be to rigorously test if encoding raw sequence and quality data as an image, tensor, or some other mixed format produces the best variant calls. Because many of the latest neural network architectures (ResNet, Inception, Xception, and others) are already optimized for and pre-trained on generic, large-scale image datasets [341], encoding genomic data as images could prove to be a generally effective and efficient strategy.

In limited experiments, DeepVariant was robust to sequencing depth, read length, and even species [338]. However, a model built on Illumina data, for instance, may not be optimal for Pacific Biosciences long-read data or MinION nanopore data, which have vastly different specificity and sensitivity profiles and signal-to-noise characteristics. Recently, Boza et al. used bidirectional recurrent neural networks to infer the *E. coli* sequence from MinION nanopore electric current data with higher per-base accuracy than the proprietary hidden Markov model-based algorithm Metrichor [336]. Unfortunately, training any neural network requires a large amount of data, which is often not available for new sequencing technologies. To circumvent this, one very preliminary study simulated mutations and spiked them into somatic and germline RNA-seq data, then trained and tested a neural network on simulated paired RNA-seq and exome sequencing data [342]. However, because this model was not subsequently tested on ground-truth datasets, it is unclear whether simulation can produce sufficiently realistic data to produce reliable models.

Method development for interpreting new types of sequencing data has historically taken two steps: first, easily implemented hard cutoffs that prioritize specificity over sensitivity, then expert development of probabilistic models with hand-developed inputs [342]. We anticipate that these steps will be replaced by deep learning, which will infer features simply by its ability to optimize a complex model against data.

### Neuroscience

Artificial neural networks were originally conceived as a model for computation in the brain [6]. Although deep neural networks have evolved to become a workhorse across many fields, there is still a strong connection between deep networks and the study of the brain. The rich parallel history of artificial neural networks in computer science and neuroscience is reviewed in [343–345].

Convolutional neural networks were originally conceived as faithful models of visual information processing in the primate visual system, and are still considered so [346]. The activations of hidden units in consecutive layers of deep convolutional networks have been found to parallel the activity of neurons in consecutive brain regions involved in processing visual scenes. Such models of neural computation are called “encoding” models, as they predict how the nervous system might encode sensory information in the world.

Even when they are not directly modeling biological neurons, deep networks have been a useful computational tool in neuroscience. They have been developed as statistical time series models of neural activity in the brain. And in contrast to the encoding models described earlier, these models are used for decoding neural activity, for instance in brain machine interfaces [347]. They have been crucial to the field of connectomics, which is concerned with mapping the connectivity of biological neural networks in the brain. In connectomics, deep networks are used to segment the shapes of individual neurons and to infer their connectivity from 3D electron microscopic images [348], and they have been also been used to infer causal connectivity from optical measurement and perturbation of neural activity [349].

It is an exciting time for neuroscience. Recent rapid progress in deep networks continues to inspire new machine learning based models of brain computation [343]. And neuroscience continues to inspire new models of artificial intelligence [345].

## The impact of deep learning in treating disease and developing new treatments

Given the need to make better, faster interventions at the point of care—incorporating the complex calculus of a patients symptoms, diagnostics, and life history—there have been many attempts to apply deep learning to patient treatment. Success in this area could help to enable personalized healthcare or precision medicine [350, 351]. Earlier, we reviewed approaches for patient categorization. Here, we examine the potential for better treatment, which broadly, may be divided into methods for improved choices of interventions for patients and those for development of new interventions.

### Clinical decision making

In 1996, Tu [352] compared the effectiveness of artificial neural networks and logistic regression, questioning whether these techniques would replace traditional statistical methods for predicting medical outcomes such as myocardial infarction [353] or mortality [354]. He posited that while neural networks have several advantages in representational power, the difficulties in interpretation may limit clinical applications, a limitation that still remains today. In addition, the challenges faced by physicians parallel those encountered by deep learning. For a given patient, the number of possible diseases is very large, with a long tail of rare diseases and patients are highly heterogeneous and may present with very different signs and symptoms for the same disease. Still, in 2006 Lisboa and Taktak [355] examined the use of artificial neural networks in medical journals, concluding that they improved healthcare relative to traditional screening methods in 21 of 27 studies.

While further progress has been made in using deep learning for clinical decision making, it is hindered by a challenge common to many deep learning applications: it is much easier to predict an outcome than to suggest an action to change the outcome. Several attempts [118, 120] at recasting the clinical decision-making problem into a prediction problem (i.e. prediction of which treatment will most improve the patient’s health) have accurately predicted survival patterns, but technical and medical challenges remain for clinical adoption (similar to those for categorization). In particular, remaining barriers include actionable interpretability of deep learning models, fitting deep models to limited and heterogeneous data, and integrating complex predictive models into a dynamic clinical environment.

A critical challenge in providing treatment recommendations is identifying a causal relationship for each recommendation. Causal inference is often framed in terms of counterfactual question [356]. Johansson et al. [357] use deep neural networks to create representation models for covariates that capture nonlinear effects and show significant performance improvements over existing models. In a less formal approach, Kale et al. [358] first create a deep neural network to model clinical time series and then analyze the relationship of the hidden features to the output using a causal approach.

A common challenge for deep learning is the interpretability of the models and their predictions. The task of clinical decision making is necessarily risk-averse, so model interpretability is key. Without clear reasoning, it is difficult to establish trust in a model. As described above, there has been some work to directly assign treatment plans without interpretability; however, the removal of human experts from the decision-making loop make the models difficult to integrate with clinical practice. To alleviate this challenge, several studies have attempted to create more interpretable deep models, either specifically for healthcare or as a general procedure for deep learning (see Discussion).

### Predicting patient trajectories

A common application for deep learning in this domain is the temporal structure of healthcare records. Many studies [359–362] have used RNNs to categorize patients, but most stop short of suggesting clinical decisions. Nemati et al. [363] used deep reinforcement learning to optimize a heparin dosing policy for intensive care patients. However, because the ideal dosing policy is unknown, the model’s predictions must be evaluated on counter-factual data. This represents a common challenge when bridging the gap between research and clinical practice. Because the ground-truth is unknown, researchers struggle to evaluate model predictions in the absence of interventional data, but clinical application is unlikely until the model has been shown to be effective. The impressive applications of deep reinforcement learning to other domains [304] have relied on knowledge of the underlying processes (e.g. the rules of the game). Some models have been developed for targeted medical problems [364], but a generalized engine is beyond current capabilities.

### Clinical trial efficiency

A clinical deep learning task that has been more successful is the assignment of patients to clinical trials. Ithapu et al. [365] used a randomized denoising autoencoder to learn a multimodal imaging marker that predicts future cognitive and neural decline from positron emission tomography (PET), amyloid florbetapir PET, and structural magnetic resonance imaging. By accurately predicting which cases will progress to dementia, they were able to efficiently assign patients to a clinical trial and reduced the required sample sizes by a factor of five. Similarly, Artemov et al. [366] applied deep learning to predict which clinical trials were likely to fail and which were likely to succeed. By predicting the side effects and pathway activations of each drug and translating these activations to a success probability, their deep learning-based approach was able to significantly outperform a random forest classifier trained on gene expression changes. These approaches suggest promising directions to improve the efficiency of clinical trials and accelerate drug development.

### Drug repositioning

Drug repositioning (or repurposing) is an attractive option for delivering new drugs to the market because of the high costs and failure rates associated with more traditional drug discovery approaches [367, 368]. A decade ago, the Connectivity Map [369] had a sizeable impact. Reverse matching disease gene expression signatures with a large set of reference compound profiles allowed researchers to formulate repurposing hypotheses at scale using a simple non-parametric test. Since then, several advanced computational methods have been applied to formulate and validate drug repositioning hypotheses [370–372]. Using supervised learning and collaborative filtering to tackle this type of problem is proving successful, especially when coupling disease or compound omic data with topological information from protein-protein or protein-compound interaction networks [373–375].

For example, Menden et al. [376] used a shallow neural network to predict sensitivity of cancer cell lines to drug treatment using both cell line and drug features, opening the door to precision medicine and drug repositioning opportunities in cancer. More recently, Aliper et al. [36] used gene-and pathway-level drug perturbation transcriptional profiles from the Library of Network-Based Cellular Signatures [377] to train a fully connected deep neural network to predict drug therapeutic uses and indications. By using confusion matrices and leveraging misclassification, the authors formulated a number of interesting hypotheses, including repurposing cardiovascular drugs such as otenzepad and pinacidil for neurological disorders.

Drug repositioning can also be approached by attempting to predict novel drug-target interactions and then repurposing the drug for the associated indication [378, 379]. Wang et al. [380] devised a pairwise input neural network with two hidden layers that takes two inputs, a drug and a target binding site, and predicts whether they interact. Wang et al. [37] trained individual RBMs for each target in a drug-target interaction network and used these models to predict novel interactions pointing to new indications for existing drugs. Wen et al. [38] extended this concept to deep learning by creating a DBN called DeepDTIs, which predicts interactions using chemical structure and protein sequence features.

Drug repositioning appears an obvious candidate for deep learning both because of the large amount of high-dimensional data available and the complexity of the question being asked. However, perhaps the most promising piece of work in this space [36] is more of a proof of concept than a real-world hypothesis-generation tool; notably, deep learning was used to predict drug indications but not for the actual repositioning. At present, some of the most popular state-of-the-art methods for signature-based drug repurposing [381] do not use predictive modeling. A mature and production-ready framework for drug repositioning via deep learning is currently missing.

### Drug development

#### Ligand-based prediction of bioactivity

High-throughput chemical screening in biomedical research aims to improve therapeutic options over a long term horizon [21]. The objective is to discover which small molecules (also referred to as chemical compounds or ligands) specifically affect the activity of a target, such as a kinase, protein-protein interaction, or broader cellular phenotype. This screening is often one of the first steps in a long drug discovery pipeline, where novel molecules are pursued for their ability to inhibit or enhance disease-relevant biological mechanisms [382]. Initial hits are confirmed to eliminate false positives and proceed to the lead generation stage [383], where they are evaluated for absorption, distribution, metabolism, excretion, and toxicity (ADMET) and other properties. It is desirable to advance multiple lead series, clusters of structurally-similar active chemicals, for further optimization by medicinal chemists to protect against unexpected failures in the later stages of drug discovery [382].

Computational work in this domain aims to identify sufficient candidate active compounds without exhaustively screening libraries of hundreds of thousands or millions of chemicals. Predicting chemical activity computationally is known as virtual screening. An ideal algorithm will rank a sufficient number of active compounds before the inactives, but the rankings of actives relative to other actives and inactives are less important [384]. Computational modeling also has the potential to predict ADMET traits for lead generation [385] and how drugs are metabolized [386].

Ligand-based approaches train on chemicals’ features without modeling target features (e.g. protein structure). Neural networks have a long history in this domain [19, 22], and the 2012 Merck Molecular Activity Challenge on Kaggle generated substantial excitement about the potential for high-parameter deep learning approaches. The winning submission was an ensemble that included a multi-task multi-layer perceptron network [387]. The sponsors noted drastic improvements over a random forest baseline, remarking “we have seldom seen any method in the past 10 years that could consistently outperform [random forest] by such a margin” [388], but not all outside experts were convinced [389]. Subsequent work (reviewed in more detail by Goh et al. [20]) explored the effects of jointly modeling far more targets than the Merck challenge [390, 391], with Ramsundar et al. [391] showing that the benefits of multi-task networks had not yet saturated even with 259 targets. Although DeepTox [392], a deep learning approach, won another competition, the Toxicology in the 21st Century (Tox21) Data Challenge, it did not dominate alternative methods as thoroughly as in other domains. DeepTox was the top performer on 9 of 15 targets and highly competitive with the top performer on the others. However, for many targets there was little separation between the top two or three methods.

The nuanced Tox21 performance may be more reflective of the practical challenges encountered in ligand-based chemical screening than the extreme enthusiasm generated by the Merck competition. A study of 22 ADMET tasks demonstrated that there are limitations to multi-task transfer learning that are in part a consequence of the degree to which tasks are related [385]. Some of the ADMET datasets showed superior performance in multi-task models with only 22 ADMET tasks compared to multi-task models with over 500 less-similar tasks. In addition, the training datasets encountered in practical applications may be tiny relative to what is available in public datasets and organized competitions. A study of BACE-1 inhibitors included only 1547 compounds [393]. Machine learning models were able to train on this limited dataset, but overfitting was a challenge and the differences between random forests and a deep neural network were negligible, especially in the classification setting. Overfitting is still a problem in larger chemical screening datasets with tens or hundreds of thousands of compounds because the number of active compounds can be very small, on the order of 0.1% of all tested chemicals for a typical target [394]. This has motivated low-parameter neural networks that emphasize compound-compound similarity, such as influence-relevance voter [384, 395], instead of predicting compound activity directly from chemical features.

#### Chemical featurization and representation learning

Much of the recent excitement in this domain has come from what could be considered a creative experimentation phase, in which deep learning has offered novel possibilities for feature representation and modeling of chemical compounds. A molecular graph, where atoms are labeled nodes and bonds are labeled edges, is a natural way to represent a chemical structure. Chemical features can be represented as a list of molecular descriptors such as molecular weight, atom counts, functional groups, charge representations, summaries of atom-atom relationships in the molecular graph, and more sophisticated derived properties [396]. Traditional machine learning approaches relied on preprocessing the graph into a feature vector of molecular descriptors or a fixed-width bit vector known as a fingerprint [397]. The same fingerprints have been used by some drug-target interaction methods discussed above [38]. An overly simplistic but approximately correct view of chemical fingerprints is that each bit represents the presence or absence of a particular chemical substructure in the molecular graph. Instead of using molecular descriptors or fingerprints as input, modern neural networks can represent chemicals as textual strings [398] or images [399] or operate directly on the molecular graph, which has enabled strategies for learning novel chemical representations.

Virtual screening and chemical property prediction have emerged as one of the major applications areas for graph-based neural networks. Duvenaud et al. [400] generalized standard circular fingerprints by substituting discrete operations in the fingerprinting algorithm with operations in a neural network, producing a real-valued feature vector instead of a bit vector. Other approaches offer trainable networks that can learn chemical feature representations that are optimized for a particular prediction task. Lusci et al. [401] applied recursive neural networks for directed acyclic graphs to undirected molecular graphs by creating an ensemble of directed graphs in which one atom is selected as the root node. Graph convolutions on undirected molecular graphs have eliminated the need to enumerate artificially directed graphs, learning feature vectors for atoms that are a function of the properties of neighboring atoms and local regions on the molecular graph [402 –404]. More sophisticated graph algorithms [405, 406] addressed limitations of standard graph convolutions that primarily operate on each node’s local neighborhood. We anticipate that these graph-based neural networks could also be applicable in other types of biological networks, such as the PPI networks we discussed previously.

Advances in chemical representation learning have also enabled new strategies for learning chemical-chemical similarity functions. Altae-Tran et al. developed a one-shot learning network [403] to address the reality that most practical chemical screening studies are unable to provide the thousands or millions of training compounds that are needed to train larger multi-task networks. Using graph convolutions to featurize chemicals, the network learns an embedding from compounds into a continuous feature space such that compounds with similar activities in a set of training tasks have similar embeddings. The approach is evaluated in an extremely challenging setting. The embedding is learned from a subset of prediction tasks (e.g. activity assays for individual proteins), and only one to ten labeled examples are provided as training data on a new task. On Tox21 targets, even when trained with *one* task-specific active compound and *one* inactive compound, the model is able to generalize reasonably well because it has learned an informative embedding function from the related tasks. Random forests, which cannot take advantage of the related training tasks, trained in the same setting are only slightly better than a random classifier. Despite the success on Tox21, performance on MUV datasets, which contains assays designed to be challenging for chemical informatics algorithms, is considerably worse. The authors also demonstrate the limitations of transfer learning as embeddings learned from the Tox21 assays have little utility for a drug adverse reaction dataset.

These novel, learned chemical feature representations may prove to be essential for accurately predicting why some compounds with similar structures yield similar target effects and others produce drastically different results. Currently, these methods are enticing but do not necessarily outperform classic approaches by a large margin. The neural fingerprints [400] were narrowly beaten by regression using traditional circular fingerprints on a drug efficacy prediction task but were superior for predicting solubility or photovoltaic efficiency. In the original study, graph convolutions [402] performed comparably to a multi-task network using standard fingerprints and slightly better than the neural fingerprints [400] on the drug efficacy task but were slightly worse than the influence-relevance voter method on an HIV dataset [384]. Broader recent benchmarking has shown that relative merits of these methods depends on the dataset and cross validation strategy [407], though evaluation in this domain often uses area under the receiver operating characteristic curve (AUROC) [408], which has limited utility due to the large class imbalance (see Discussion).

We remain optimistic for the potential of deep learning and specifically representation learning in drug discovery. Rigorous benchmarking on broad and diverse prediction tasks will be as important as novel neural network architectures to advance the state of the art and convincingly demonstrate superiority over traditional cheminformatics techniques. Fortunately, there has recently been much progress in this direction. The DeepChem software [403, 409] and MoleculeNet benchmarking suite [407] built upon it contain chemical bioactivity and toxicity prediction datasets, multiple compound featurization approaches including graph convolutions, and various machine learning algorithms ranging from standard baselines like logistic regression and random forests to recent neural network architectures. Independent research groups have already contributed additional datasets and prediction algorithms to DeepChem. Adoption of common benchmarking evaluation metrics, datasets, and baseline algorithms has the potential to establish the practical utility of deep learning in chemical bioactivity prediction and lower the barrier to entry for machine learning researchers without biochemistry expertise.

One open question in ligand-based screening pertains to the benefits and limitations of transfer learning. Multi-task neural networks have shown the advantages of jointly modeling many targets [390, 391]. Other studies have shown the limitations of transfer learning when the prediction tasks are insufficiently related [385, 403]. This has important implications for representation learning. The typical approach to improve deep learning models by expanding the dataset size may not be applicable if only “related” tasks are beneficial, especially because task-task relatedness is ill-defined. The massive chemical state space will also influence the development of unsupervised representation learning methods [398, 410]. Future work will establish whether it is better to train on massive collections of diverse compounds, drug-like small molecules, or specialized subsets.

#### Structure-based prediction of bioactivity

When protein structure is available, virtual screening has traditionally relied on docking programs to predict how a compound best fits in the target’s binding site and score the predicted ligand-target complex [411]. Recently, deep learning approaches have been developed to model protein structure, which is expected to improve upon the simpler drug-target interaction algorithms described above that represent proteins with feature vectors derived from amino acid sequences [38, 380].

Structure-based deep learning methods differ in whether they use experimentally-derived or predicted ligand-target complexes and how they represent the 3D structure. The Atomic CNN [412] and TopologyNet [413] models take 3D structures from PDBBind [414] as input, ensuring the ligand-target complexes are reliable. AtomNet [35] samples multiple ligand poses within the target binding site, and DeepVS [415] and Ragoza et al. [416] use a docking program to generate protein-compound complexes. If they are sufficiently accurate, these latter approaches would have wider applicability to a much larger set of compounds and proteins. However, incorrect ligand poses will be misleading during training, and the predictive performance is sensitive to the docking quality [415].

There are two established options for representing a protein-compound complex. One option, a 3D grid, can featurize the input complex [35, 416]. Each entry in the grid tracks the types of protein and ligand atoms in that region of the 3D space or descriptors derived from those atoms. Alternatively, DeepVS [415] and atomic convolutions [412] offer greater flexibility in their convolutions by eschewing the 3D grid. Instead, they each implement techniques for executing convolutions over atoms’ neighboring atoms in the 3D space. Gomes et al. demonstrate that currently random forest on a 1D feature vector that describes the 3D ligand-target structure generally outperforms neural networks on the same feature vector as well as atomic convolutions and ligand-based neural networks when predicting the continuous-valued inhibition constant on the PDBBind refined dataset [412]. However, in the long term, atomic convolutions may ultimately overtake grid-based methods, as they provide greater freedom to model atom-atom interactions and the forces that govern binding affinity.

#### *De novo* drug design

*De novo* drug design attempts to model the typical design-synthesize-test cycle of drug discovery [417, 418]. It explores an estimated 10^60^ synthesizable organic molecules with drug-like properties without explicit enumeration [394]. To test or score structures, algorithms like those discussed earlier are used. To “design” and “synthesize”, traditional *de novo* design software relied on classical optimizers such as genetic algorithms. Unfortunately, this often leads to overfit, “weird” molecules, which are difficult to synthesize in the lab. Current programs have settled on rule-based virtual chemical reactions to generate molecular structures [418]. Deep learning models that generate realistic, synthesizable molecules have been proposed as an alternative. In contrast to the classical, symbolic approaches, generative models learned from data would not depend on laboriously encoded expert knowledge. The challenge of generating molecules has parallels to the generation of syntactically and semantically correct text [419].

As deep learning models that directly output (molecular) graphs remain under-explored, generative neural networks for drug design typically represent chemicals with the simplified molecular-input line-entry system (SMILES), a standard string-based representation with characters that represent atoms, bonds, and rings [420]. This allows treating molecules as sequences and leveraging recent progress in recurrent neural networks. Gómez-Bombarelli et al. designed a SMILES-to-SMILES autoencoder to learn a continuous latent feature space for chemicals [398]. In this learned continuous space it was possible to interpolate between continuous representations of chemicals in a manner that is not possible with discrete (e.g. bit vector or string) features or in symbolic, molecular graph space. Even more interesting is the prospect of performing gradient-based or Bayesian optimization of molecules within this latent space. The strategy of constructing simple, continuous features before applying supervised learning techniques is reminiscent of autoencoders trained on high-dimensional EHR data [112]. A drawback of the SMILES-to-SMILES autoencoder is that not all SMILES strings produced by the autoencoder’s decoder correspond to valid chemical structures. Recently, the Grammar Variational Autoencoder, which takes the SMILES grammar into account and is guaranteed to produce syntactically valid SMILES, has been proposed to alleviate this issue [421].

Another approach to *de novo* design is to train character-based RNNs on large collections of molecules, for example, ChEMBL [422], to first obtain a generic generative model for drug-like compounds [420]. These generative models successfully learn the grammar of compound representations, with 94% [423] or nearly 98% [420] of generated SMILES corresponding to valid molecular structures. The initial RNN is then fine-tuned to generate molecules that are likely to be active against a specific target by either continuing training on a small set of positive examples [420] or adopting reinforcement learning strategies [423, 424]. Both the fine-tuning and reinforcement learning approaches can rediscover known, held-out active molecules. The great flexibility of neural networks, and progress in generative models offers many opportunities for deep architectures in *de novo* design (e.g. the adaptation of GANs for molecules).

## Discussion

Despite the disparate types of data and scientific goals in the learning tasks covered above, several challenges are broadly important for deep learning in the biomedical domain. Here we examine these factors that may impede further progress, ask what steps have already been taken to overcome them, and suggest future research directions.

### Customizing deep learning models reflects a tradeoff between bias and variance

Some of the challenges in applying deep learning are shared with other machine learning methods. In particular, many problem-specific optimizations described in this review reflect a recurring universal tradeoff—controlling the flexibility of a model in order to maximize predictivity. Methods for adjusting the flexibility of deep learning models include dropout, reduced data projections, and transfer learning (described below). One way of understanding such model optimizations is that they incorporate external information to limit model flexibility and thereby improve predictions. This balance is formally described as a tradeoff between “bias and variance” [10].

Although the bias-variance tradeoff is common to all machine learning applications, recent empirical and theoretical observations suggest that deep learning models may have uniquely advantageous generalization properties [425, 426]. Nevertheless, additional advances will be needed to establish a coherent theoretical foundation that enables practitioners to better reason about their models from first principles.

### Evaluation metrics for imbalanced classification

Making predictions in the presence of high class imbalance and differences between training and generalization data is a common feature of many large biomedical datasets, including deep learning models of genomic features, patient classification, disease detection, and virtual screening. Prediction of transcription factor binding sites exemplifies the difficulties with learning from highly imbalanced data. The human genome has 3 billion base pairs, and only a small fraction of them are implicated in specific biochemical activities. Less than 1% of the genome can be confidently labeled as bound for most transcription factors.

Estimating the false discovery rate (FDR) is a standard method of evaluation in genomics that can also be applied to deep learning model predictions of genomic features. Using deep learning predictions for targeted validation experiments of specific biochemical activities necessitates a more stringent FDR (typically 5–25%). However, when predicted biochemical activities are used as features in other models, such as gene expression models, a low FDR may not be necessary.

What is the correspondence between FDR metrics and commonly used classification metrics such as AUPR and AUROC? AUPR evaluates the average precision, or equivalently, the average FDR across all recall thresholds. This metric provides an overall estimate of performance across all possible use cases, which can be misleading for targeted validation experiments. For example, classification of TF binding sites can exhibit a recall of 0% at 10% FDR and AUPR greater than 0.6. In this case, the AUPR may be competitive, but the predictions are ill-suited for targeted validation that can only examine a few of the highest-confidence predictions. Likewise, AUROC evaluates the average recall across all false positive rate (FPR) thresholds, which is often a highly misleading metric in class-imbalanced domains [70, 427]. Consider a classification model with recall of 0% at FDR less than 25% and 100% recall at FDR greater than 25%. In the context of TF binding predictions where only 1% of genomic regions are bound by the TF, this is equivalent to a recall of 100% for FPR greater than 0.33%. In other words, the AUROC would be 0.9967, but the classifier would be useless for targeted validation. It is not unusual to obtain a chromosome-wide AUROC greater than 0.99 for TF binding predictions but a recall of 0% at 10% FDR. Consequently, practitioners must select the metric most tailored to their subsequent use case to use these methods most effectively.

### Formulation of classification labels

Genome-wide continuous signals are commonly formulated into classification labels through signal peak detection. ChIP-seq peaks are used to identify locations of TF binding and histone modifications. Such procedures rely on thresholding criteria to define what constitutes a peak in the signal. This inevitably results in a set of signal peaks that are close to the threshold, not sufficient to constitute a positive label but too similar to positively labeled examples to constitute a negative label. To avoid an arbitrary label for these examples they may be labeled as “ambiguous”. Ambiguously labeled examples can then be ignored during model training and evaluation of recall and FDR. The correlation between model predictions on these examples and their signal values can be used to evaluate if the model correctly ranks these examples between positive and negative examples.

### Formulation of a performance upper bound

In assessing the upper bound on the predictive performance of a deep learning model, it is necessary to incorporate inherent between-study variation inherent to biomedical research [428]. Study-level variability limits classification performance and can lead to underestimating prediction error if the generalization error is estimated by splitting a single dataset. Analyses can incorporate data from multiple labs and experiments to capture between-study variation within the prediction model mitigating some of these issues.

### Uncertainty quantification

Deep learning based solutions for biomedical applications could substantially benefit from guarantees on the reliability of predictions and a quantification of uncertainty. Due to biological variability and precision limits of equipment, biomedical data do not consist of precise measurements but of estimates with noise. Hence, it is crucial to obtain uncertainty measures that capture how noise in input values propagate through deep neural networks. Such measures can be used for reliability assessment of automated decisions in clinical and public health applications, and for guarding against model vulnerabilities in the face of rare or adversarial cases [429]. Moreover, in fundamental biological research, measures of uncertainty help researchers distinguish between true regularities in the data and patterns that are false or merely anecdotal. There are two main uncertainties that one can calculate: epistemic and aleatoric [430]. Epistemic uncertainty describes uncertainty about the model, its structure, or its parameters. This uncertainty is caused by insufficient training data or by a difference in the training set and testing set distributions, so it vanishes in the limit of infinite data. On the other hand, aleatoric uncertainty describes uncertainty inherent in the observations. This uncertainty is due to noisy or missing data, so it vanishes with the ability to observe all independent variables with infinite precision. A good way to represent aleatoric uncertainty is to design an appropriate loss function with an uncertainty variable. In the case of data-dependent aleatoric uncertainty, one can train the model to increase its uncertainty when it is incorrect due to noisy or missing data, and in the case of task-dependent aleatoric uncertainty, one can optimize for the best uncertainty parameter for each task [431]. Meanwhile, there are various methods for modeling epistemic uncertainty, outlined below.

In classification tasks, confidence calibration is the problem of using classifier scores to predict class membership probabilities that match the true membership likelihoods. These membership probabilities can be used to assess the uncertainty associated with assigning the example to each of the classes. Guo et al. [432] observed that contemporary neural networks are poorly calibrated and provided a simple recommendation for calibration: temperature scaling, a single parameter special case of Platt scaling [433]. In addition to confidence calibration, there is early work from Chryssolouris et al. [434] that described a method for obtaining confidence intervals with the assumption of normally distributed error for the neural network. More recently, Hendrycks and Gimpel discovered that incorrect or out-of-distribution examples usually have lower maximum softmax probabilities than correctly classified examples, allowing for effective detection of misclassified examples [435]. Liang et al. used temperature scaling and small perturbations to further separate the softmax scores of correctly classified examples and the scores of out-of-distribution examples, allowing for more effective detection [436]. This approach outperformed the baseline approaches by a large margin, establishing a new state-of-the-art performance.

An alternative approach for obtaining principled uncertainty estimates from deep learning models is to use Bayesian neural networks. Deep learning models are usually trained to obtain the most likely parameters given the data. However, choosing the single most likely set of parameters ignores the uncertainty about which set of parameters (among the possible models that explain the given dataset) should be used. This sometimes leads to uncertainty in predictions when the chosen likely parameters produce high-confidence but incorrect results. On the other hand, the parameters of Bayesian neural networks are modeled as full probability distributions. This Bayesian approach comes with a whole host of benefits, including better calibrated confidence estimates [437] and more robustness to adversarial and out-of-distribution examples [438]. Unfortunately, modeling the full posterior distribution for the model’s parameters given the data is usually computationally intractable. One popular method for circumventing this high computational cost is called test-time dropout [439], where an approximate posterior distribution is obtained using variational inference. Gal and Ghahramani showed that a stack of fully connected layers with dropout between the layers is equivalent to approximate inference in a Gaussian process model [439]. The authors interpret dropout as a variational inference method and apply their method to convolutional neural networks. This is simple to implement and preserves the possibility of obtaining cheap samples from the approximate posterior distribution. Operationally, obtaining model uncertainty for a given case becomes as straightforward as leaving dropout turned on and predicting multiple times. The spread of the different predictions is a reasonable proxy for model uncertainty. This technique has been successfully applied in an automated system for detecting diabetic retinopathy [440], where uncertainty-informed referrals improved diagnostic performance and allowed the model to meet the National Health Service recommended levels of sensitivity and specificity. The authors also found that entropy performs comparably to the spread obtained via test-time dropout for identifying uncertain cases, and therefore it can be used instead for automated referrals.

Several other techniques have been proposed for effectively estimating predictive uncertainty as uncertainty quantification for neural networks continues to be an active research area. Recently, McClure and Kriegeskorte observed that test-time sampling improved calibration of the probabilistic predictions, sampling weights led to more robust uncertainty estimates than sampling units, and spike-and-slab sampling is superior to Gaussian dropconnect and Bernoulli dropout [441]. Krueger et al. introduced Bayesian hypernetworks [442] as another framework for approximate Bayesian inference in deep learning, where an invertible generative hypernetwork maps isotropic Gaussian noise to parameters of the primary network allowing for computationally cheap sampling and efficient estimation of the posterior. Meanwhile, Lakshminarayanan et al. proposed using deep ensembles, which are traditionally used for boosting predictive performance, on standard (non-Bayesian) neural networks to obtain well-calibrated uncertainty estimates that are comparable to those obtained by Bayesian neural networks [443]. In cases where model uncertainty is known to be caused by a difference in training and testing distributions, domain adaptation based techniques can help mitigate the problem [218].

Despite the success and popularity of deep learning, some deep learning models can be surprisingly brittle. Researchers are actively working on modifications to deep learning frameworks to enable them to handle probability and embrace uncertainty. Most notably, Bayesian modeling and deep learning are being integrated with renewed enthusiasm. As a result, several opportunities for innovation arise: understanding the causes of model uncertainty can lead to novel optimization and regularization techniques, assessing the utility of uncertainty estimation techniques on various model architectures and structures can be very useful to practitioners, and extending Bayesian deep learning to unsupervised settings can be a significant breakthrough [444]. Unfortunately, uncertainty quantification techniques are underutilized in the computational biology communities and largely ignored in the current deep learning for biomedicine literature. Thus, the practical value of uncertainty quantification in biomedical domains is yet to be appreciated.

### Interpretation

As deep learning models achieve state-of-the-art performance in a variety of domains, there is a growing need to make the models more interpretable. Interpretability matters for two main reasons. First, a model that achieves breakthrough performance may have identified patterns in the data that practitioners in the field would like to understand. However, this would not be possible if the model is a black box. Second, interpretability is important for trust. If a model is making medical diagnoses, it is important to ensure the model is making decisions for reliable reasons and is not focusing on an artifact of the data. A motivating example of this can be found in Ba and Caruana [445], where a model trained to predict the likelihood of death from pneumonia assigned lower risk to patients with asthma, but only because such patients were treated as higher priority by the hospital. In the context of deep learning, understanding the basis of a model’s output is particularly important as deep learning models are unusually susceptible to adversarial examples [446] and can output confidence scores over 99.99% for samples that resemble pure noise.

As the concept of interpretability is quite broad, many methods described as improving the interpretability of deep learning models take disparate and often complementary approaches.

### Assigning example-specific importance scores

Several approaches ascribe importance on an example-specific basis to the parts of the input that are responsible for a particular output. These can be broadly divided into perturbation-based approaches and backpropagation-based approaches.

Perturbation-based approaches change parts of the input and observe the impact on the output of the network. Alipanahi et al. [201] and Zhou & Troyanskaya [209] scored genomic sequences by introducing virtual mutations at individual positions in the sequence and quantifying the change in the output. Umarov et al. [222] used a similar strategy, but with sliding windows where the sequence within each sliding window was substituted with a random sequence. Kelley et al. [227] inserted known protein-binding motifs into the centers of sequences and assessed the change in predicted accessibility. Ribeiro et al. [447] introduced LIME, which constructs a linear model to locally approximate the output of the network on perturbed versions of the input and assigns importance scores accordingly. For analyzing images, Zeiler and Fergus [448] applied constant-value masks to different input patches. More recently, marginalizing over the plausible values of an input has been suggested as a way to more accurately estimate contributions [449].

A common drawback to perturbation-based approaches is computational efficiency: each perturbed version of an input requires a separate forward propagation through the network to compute the output. As noted by Shrikumar et al. [219], such methods may also underestimate the impact of features that have saturated their contribution to the output, as can happen when multiple redundant features are present. To reduce the computational overhead of perturbation-based approaches, Fong and Vedaldi [450] solve an optimization problem using gradient descent to discover a minimal subset of inputs to perturb in order to decrease the predicted probability of a selected class. Their method converges in many fewer iterations but requires the perturbation to have a differentiable form.

Backpropagation-based methods, in which the signal from a target output neuron is propagated backwards to the input layer, are another way to interpret deep networks that sidestep inefficiencies of the perturbation-based methods. A classic example of this is calculating the gradients of the output with respect to the input [451] to compute a “saliency map”. Bach et al. [452] proposed a strategy called Layerwise Relevance Propagation, which was shown to be equivalent to the element-wise product of the gradient and input [219, 453]. Networks with Rectified Linear Units (ReLUs) create nonlinearities that must be addressed. Several variants exist for handling this [448, 454]. Backpropagation-based methods are a highly active area of research. Researchers are still actively identifying weaknesses [455], and new methods are being developed to address them [219,456,457]. Lundberg and Lee [458] noted that several importance scoring methods including integrated gradients and LIME could all be considered approximations to Shapely values [459], which have a long history in game theory for assigning contributions to players in cooperative games.

### Matching or exaggerating the hidden representation

Another approach to understanding the network’s predictions is to find artificial inputs that produce similar hidden representations to a chosen example. This can elucidate the features that the network uses for prediction and drop the features that the network is insensitive to. In the context of natural images, Mahendran and Vedaldi [460] introduced the “inversion” visualization, which uses gradient descent and backpropagation to reconstruct the input from its hidden representation. The method required placing a prior on the input to favor results that resemble natural images. For genomic sequence, Finnegan and Song [461] used a Markov chain Monte Carlo algorithm to find the maximum-entropy distribution of inputs that produced a similar hidden representation to the chosen input.

A related idea is “caricaturization”, where an initial image is altered to exaggerate patterns that the network searches for [462]. This is done by maximizing the response of neurons that are active in the network, subject to some regularizing constraints. Mordvintsev et al. [463] leveraged caricaturization to generate aesthetically pleasing images using neural networks.

### Activation maximization

Activation maximization can reveal patterns detected by an individual neuron in the network by generating images which maximally activate that neuron, subject to some regularizing constraints. This technique was first introduced in Ehran et al. [464] and applied in subsequent work [451,462,463,465]. Lanchantin et al. [204] applied class-based activation maximization to genomic sequence data. One drawback of this approach is that neural networks often learn highly distributed representations where several neurons cooperatively describe a pattern of interest. Thus, visualizing patterns learned by individual neurons may not always be informative.

### RNN-specific approaches

Several interpretation methods are specifically tailored to recurrent neural network architectures. The most common form of interpretability provided by RNNs is through attention mechanisms, which have been used in diverse problems such as image captioning and machine translation to select portions of the input to focus on generating a particular output [466, 467]. Deming et al. [468] applied the attention mechanism to models trained on genomic sequence. Attention mechanisms provide insight into the model’s decision-making process by revealing which portions of the input are used by different outputs. Singh et al. used a hierarchy of attention layers to locate important genome positions and signals for predicting gene expression from histone modifications [183]. In the clinical domain, Choi et al. [469] leveraged attention mechanisms to highlight which aspects of a patient’s medical history were most relevant for making diagnoses. Choi et al. [470] later extended this work to take into account the structure of disease ontologies and found that the concepts represented by the model aligned with medical knowledge. Note that interpretation strategies that rely on an attention mechanism do not provide insight into the logic used by the attention layer.

Visualizing the activation patterns of the hidden state of a recurrent neural network can also be instructive. Early work by Ghosh and Karamcheti [471] used cluster analysis to study hidden states of comparatively small networks trained to recognize strings from a finite state machine. More recently, Karpathy et al. [472] showed the existence of individual cells in LSTMs that kept track of quotes and brackets in character-level language models. To facilitate such analyses, LSTMVis [473] allows interactive exploration of the hidden state of LSTMs on different inputs.

Another strategy, adopted by Lanchatin et al. [204] looks at how the output of a recurrent neural network changes as longer and longer subsequences are supplied as input to the network, where the subsequences begin with just the first position and end with the entire sequence. In a binary classification task, this can identify those positions which are responsible for flipping the output of the network from negative to positive. If the RNN is bidirectional, the same process can be repeated on the reverse sequence. As noted by the authors, this approach was less effective at identifying motifs compared to the gradient-based backpropagation approach of Simonyan et al. [451], illustrating the need for more sophisticated strategies to assign importance scores in recurrent neural networks.

Murdoch and Szlam [474] showed that the output of an LSTM can be decomposed into a product of factors, where each factor can be interpreted as the contribution at a particular timestep. The contribution scores were then used to identify key phrases from a model trained for sentiment analysis and obtained superior results compared to scores derived via a gradient-based approach.

### Latent space manipulation

Interpretation of embedded or latent space features learned through generative unsupervised models can reveal underlying patterns otherwise masked in the original input. Embedded feature interpretation has been emphasized mostly in image and text based applications [102, 475], but applications to genomic and biomedical domains are increasing.

For example, Way and Greene trained a variational autoencoder (VAE) on gene expression from The Cancer Genome Atlas (TCGA) [476] and use latent space arithmetic to rapidly isolate and interpret gene expression features descriptive of high grade serous ovarian cancer subtypes [477]. The most differentiating VAE features were representative of biological processes that are known to distinguish the subtypes. Latent space arithmetic with features derived using other compression algorithms were not as informative in this context [478]. Embedding discrete chemical structures with autoencoders and interpreting the learned continuous representations with latent space arithmetic has also facilitated predicting drug-like compounds [398]. Furthermore, embedding biomedical text into lower dimensional latent spaces have improved name entity recognition in a variety of tasks including annotating clinical abbreviations, genes, cell lines, and drug names [75–78].

Other approaches have used interpolation through latent space embeddings learned by GANs to interpret unobserved intermediate states. For example, Osokin et al. trained GANs on two-channel fluorescent microscopy images to interpret intermediate states of protein localization in yeast cells [479]. Goldsborough et al. trained a GAN on fluorescent microscopy images and used latent space interpolation and arithmetic to reveal underlying responses to small molecule perturbations in cell lines [480].

### Miscellaneous approaches

It can often be informative to understand how the training data affects model learning. Toward this end, Koh and Liang [481] used influence functions, a technique from robust statistics, to trace a model’s predictions back through the learning algorithm to identify the datapoints in the training set that had the most impact on a given prediction. A more free-form approach to interpretability is to visualize the activation patterns of the network on individual inputs and on subsets of the data. ActiVis and CNNvis [482, 483] are two frameworks that enable interactive visualization and exploration of large-scale deep learning models. An orthogonal strategy is to use a knowledge distillation approach to replace a deep learning model with a more interpretable model that achieves comparable performance. Towards this end, Che et al. [484] used gradient boosted trees to learn interpretable healthcare features from trained deep models.

Finally, it is sometimes possible to train the model to provide justifications for its predictions. Lei et al. [485] used a generator to identify “rationales”, which are short and coherent pieces of the input text that produce similar results to the whole input when passed through an encoder. The authors applied their approach to a sentiment analysis task and obtained substantially superior results compared to an attention-based method.

### Future outlook

While deep learning lags behind most Bayesian models in terms of interpretability, the interpretability of deep learning is comparable to or exceeds that of many other widely-used machine learning methods such as random forests or SVMs. While it is possible to obtain importance scores for different inputs in a random forest, the same is true for deep learning. Similarly, SVMs trained with a nonlinear kernel are not easily interpretable because the use of the kernel means that one does not obtain an explicit weight matrix. Finally, it is worth noting that some simple machine learning methods are less interpretable in practice than one might expect. A linear model trained on heavily engineered features might be difficult to interpret as the input features themselves are difficult to interpret. Similarly, a decision tree with many nodes and branches may also be difficult for a human to make sense of.

There are several directions that might benefit the development of interpretability techniques. The first is the introduction of gold standard benchmarks that different interpretability approaches could be compared against, similar in spirit to how the ImageNet [45] and CIFAR [486] datasets spurred the development of deep learning for computer vision. It would also be helpful if the community placed more emphasis on domains outside of computer vision. Computer vision is often used as the example application of interpretability methods, but it is not the domain with the most pressing need. Finally, closer integration of interpretability approaches with popular deep learning frameworks would make it easier for practitioners to apply and experiment with different approaches to understanding their deep learning models.

### Data limitations

A lack of large-scale, high-quality, correctly labeled training data has impacted deep learning in nearly all applications we have discussed. The challenges of training complex, high-parameter neural networks from few examples are obvious, but uncertainty in the labels of those examples can be just as problematic. In genomics labeled data may be derived from an experimental assay with known and unknown technical artifacts, biases, and error profiles. It is possible to weight training examples or construct Bayesian models to account for uncertainty or non-independence in the data, as described in the TF binding example above. As another example, Park et al. [487] estimated shared non-biological signal between datasets to correct for non-independence related to assay platform or other factors in a Bayesian integration of many datasets. However, such techniques are rarely placed front and center in any description of methods and may be easily overlooked.

For some types of data, especially images, it is straightforward to augment training datasets by splitting a single labeled example into multiple examples. For example, an image can easily be rotated, flipped, or translated and retain its label [42]. 3D MRI and 4D fMRI (with time as a dimension) data can be decomposed into sets of 2D images [488]. This can greatly expand the number of training examples but artificially treats such derived images as independent instances and sacrifices the structure inherent in the data. CellCnn trains a model to recognize rare cell populations in single-cell data by creating training instances that consist of subsets of cells that are randomly sampled with replacement from the full dataset [297].

Simulated or semi-synthetic training data has been employed in multiple biomedical domains, though many of these ideas are not specific to deep learning. Training and evaluating on simulated data, for instance, generating synthetic TF binding sites with position weight matrices [207] or RNA-seq reads for predicting mRNA transcript boundaries [489], is a standard practice in bioinformatics. This strategy can help benchmark algorithms when the available gold standard dataset is imperfect, but it should be paired with an evaluation on real data, as in the prior examples [207, 489]. In rare cases, models trained on simulated data have been successfully applied directly to real data [489].

Data can be simulated to create negative examples when only positive training instances are available. DANN [34] adopts this approach to predict the pathogenicity of genetic variants using semi-synthetic training data from Combined Annotation-Dependent Depletion (CADD) [490]. Though our emphasis here is on the training strategy, it should be noted that logistic regression outperformed DANN when distinguishing known pathogenic mutations from likely benign variants in real data. Similarly, a somatic mutation caller has been trained by injecting mutations into real sequencing datasets [342]. This method detected mutations in other semi-synthetic datasets but was not validated on real data.

In settings where the experimental observations are biased toward positive instances, such as MHC protein and peptide ligand binding affinity [270], or the negative instances vastly outnumber the positives, such as high-throughput chemical screening [395], training datasets have been augmented by adding additional instances and assuming they are negative. There is some evidence that this can improve performance [395], but in other cases it was only beneficial when the real training datasets were extremely small [270]. Overall, training with simulated and semi-simulated data is a valuable idea for overcoming limited sample sizes but one that requires more rigorous evaluation on real ground-truth datasets before we can recommend it for widespread use. There is a risk that a model will easily discriminate synthetic examples but not generalize to real data.

Multimodal, multi-task, and transfer learning, discussed in detail below, can also combat data limitations to some degree. There are also emerging network architectures, such as Diet Networks for high-dimensional SNP data [491]. These use multiple networks to drastically reduce the number of free parameters by first flipping the problem and training a network to predict parameters (weights) for each input (SNP) to learn a feature embedding. This embedding (e.g. from principal component analysis, per class histograms, or a Word2vec [102] generalization) can be learned directly from input data or take advantage of other datasets or domain knowledge. Additionally, in this task the features are the examples, an important advantage when it is typical to have 500 thousand or more SNPs and only a few thousand patients. Finally, this embedding is of a much lower dimension, allowing for a large reduction in the number of free parameters. In the example given, the number of free parameters was reduced from 30 million to 50 thousand, a factor of 600.

### Hardware limitations and scaling

Efficiently scaling deep learning is challenging, and there is a high computational cost (e.g. time, memory, and energy) associated with training neural networks and using them to make predictions. This is one of the reasons why neural networks have only recently found widespread use [492].

Many have sought to curb these costs, with methods ranging from the very applied (e.g. reduced numerical precision [493–496]) to the exotic and theoretic (e.g. training small networks to mimic large networks and ensembles [445, 497]). The largest gains in efficiency have come from computation with GPUs [492,498–502], which excel at the matrix and vector operations so central to deep learning. The massively parallel nature of GPUs allows additional optimizations, such as accelerated mini-batch gradient descent [499,500,503,504]. However, GPUs also have limited memory, making networks of useful size and complexity difficult to implement on a single GPU or machine [66, 498]. This restriction has sometimes forced computational biologists to use workarounds or limit the size of an analysis. Chen et al. [181] inferred the expression level of all genes with a single neural network, but due to memory restrictions they randomly partitioned genes into two separately analyzed halves. In other cases, researchers limited the size of their neural network [28] or the total number of training instances [398]. Some have also chosen to use standard central processing unit (CPU) implementations rather than sacrifice network size or performance [505].

While steady improvements in GPU hardware may alleviate this issue, it is unclear whether advances will occur quickly enough to keep pace with the growing biological datasets and increasingly complex neural networks. Much has been done to minimize the memory requirements of neural networks [445,493–496,506,507], but there is also growing interest in specialized hardware, such as field-programmable gate arrays (FPGAs) [502, 508] and application-specific integrated circuits (ASICs) [509]. Less software is available for such highly specialized hardware [508]. But specialized hardware promises improvements in deep learning at reduced time, energy, and memory [502]. Specialized hardware may be a difficult investment for those not solely interested in deep learning, but for those with a deep learning focus these solutions may become popular.

Distributed computing is a general solution to intense computational requirements and has enabled many large-scale deep learning efforts. Some types of distributed computation [510, 511] are not suitable for deep learning [512], but much progress has been made. There now exist a number of algorithms [495,512,513], tools [514–516], and high-level libraries [517, 518] for deep learning in a distributed environment, and it is possible to train very complex networks with limited infrastructure [519]. Besides handling very large networks, distributed or parallelized approaches offer other advantages, such as improved ensembling [520] or accelerated hyperparameter optimization [521, 522].

Cloud computing, which has already seen wide adoption in genomics [523], could facilitate easier sharing of the large datasets common to biology [524, 525], and may be key to scaling deep learning. Cloud computing affords researchers flexibility, and enables the use of specialized hardware (e.g. FPGAs, ASICs, GPUs) without major investment. As such, it could be easier to address the different challenges associated with the multitudinous layers and architectures available [526]. Though many are reluctant to store sensitive data (e.g. patient electronic health records) in the cloud, secure, regulation-compliant cloud services do exist [527].

### Data, code, and model sharing

A robust culture of data, code, and model sharing would speed advances in this domain. The cultural barriers to data sharing in particular are perhaps best captured by the use of the term “research parasite” to describe scientists who use data from other researchers [528]. A field that honors only discoveries and not the hard work of generating useful data will have difficulty encouraging scientists to share their hard-won data. It’s precisely those data that would help to power deep learning in the domain. Efforts are underway to recognize those who promote an ecosystem of rigorous sharing and analysis [529].

The sharing of high-quality, labeled datasets will be especially valuable. In addition, researchers who invest time to preprocess datasets to be suitable for deep learning can make the preprocessing code (e.g. Basset [227] and variationanalysis [340]) and cleaned data (e.g. MoleculeNet [407]) publicly available to catalyze further research. However, there are complex privacy and legal issues involved in sharing patient data that cannot be ignored. Solving these issues will require increased understanding of privacy risks and standards specifying acceptable levels. In some domains high-quality training data has been generated privately, i.e. high-throughput chemical screening data at pharmaceutical companies. One perspective is that there is little expectation or incentive for this private data to be shared. However, data are not inherently valuable. Instead, the insights that we glean from them are where the value lies. Private companies may establish a competitive advantage by releasing data sufficient for improved methods to be developed. Recently, Ramsundar et al. did this with an open source platform DeepChem, where they released four privately generated datasets [530].

Code sharing and open source licensing is essential for continued progress in this domain. We strongly advocate following established best practices for sharing source code, archiving code in repositories that generate digital object identifiers, and open licensing [531] regardless of the minimal requirements, or lack thereof, set by journals, conferences, or preprint servers. In addition, it is important for authors to share not only code for their core models but also scripts and code used for data cleaning (see above) and hyperparameter optimization. These improve reproducibility and serve as documentation of the detailed decisions that impact model performance but may not be exhaustively captured in a manuscript’s methods text.

Because many deep learning models are often built using one of several popular software frameworks, it is also possible to directly share trained predictive models. The availability of pre-trained models can accelerate research, with image classifiers as an apt example. A pre-trained neural network can be quickly fine-tuned on new data and used in transfer learning, as discussed below. Taking this idea to the extreme, genomic data has been artificially encoded as images in order to benefit from pre-trained image classifiers [338]. “Model zoos”—collections of pre—trained models—are not yet common in biomedical domains but have started to appear in genomics applications [293, 532]. However, it is important to note that sharing models trained on individual data requires great care because deep learning models can be attacked to identify examples used in training. One possible solution to protect individual samples includes training models under differential privacy [152], which has been used in the biomedical domain [155]. We discussed this issue as well as recent techniques to mitigate these concerns in the patient categorization section.

DeepChem [403,407,409] and DragoNN [532] exemplify the benefits of sharing pre-trained models and code under an open source license. DeepChem, which targets drug discovery and quantum chemistry, has actively encouraged and received community contributions of learning algorithms and benchmarking datasets. As a consequence, it now supports a large suite of machine learning approaches, both deep learning and competing strategies, that can be run on diverse test cases.

This realistic, continual evaluation will play a critical role in assessing which techniques are most promising for chemical screening and drug discovery. Like formal, organized challenges such as the ENCODE-DREAM *in vivo* Transcription Factor Binding Site Prediction Challenge [213], DeepChem provides a forum for the fair, critical evaluations that are not always conducted in individual methodological papers, which can be biased toward favoring a new proposed algorithm. Likewise DragoNN (Deep RegulAtory GenOmic Neural Networks) offers not only code and a model zoo but also a detailed tutorial and partner package for simulating training data. These resources, especially the ability to simulate datasets that are sufficiently complex to demonstrate the challenges of training neural networks but small enough to train quickly on a CPU, are important for training students and attracting machine learning researchers to problems in genomics and healthcare.

### Multimodal, multi-task, and transfer learning

The fact that biomedical datasets often contain a limited number of instances or labels can cause poor performance of deep learning algorithms. These models are particularly prone to overfitting due to their high representational power. However, transfer learning techniques, also known as domain adaptation, enable transfer of extracted patterns between different datasets and even domains. This approach consists of training a model for the base task and subsequently reusing the trained model for the target problem. The first step allows a model to take advantage of a larger amount of data and/or labels to extract better feature representations. Transferring learned features in deep neural networks improves performance compared to randomly initialized features even when pre-training and target sets are dissimilar. However, transferability of features decreases as the distance between the base task and target task increases [533].

In image analysis, previous examples of deep transfer learning applications proved large-scale natural image sets [45] to be useful for pre-training models that serve as generic feature extractors for various types of biological images [14,283,534,535]. More recently, deep learning models predicted protein sub-cellular localization for proteins not originally present in a training set [536]. Moreover, learned features performed reasonably well even when applied to images obtained using different fluorescent labels, imaging techniques, and different cell types [537]. However, there are no established theoretical guarantees for feature transferability between distant domains such as natural images and various modalities of biological imaging. Because learned patterns are represented in deep neural networks in a layer-wise hierarchical fashion, this issue is usually addressed by fixing an empirically chosen number of layers that preserve generic characteristics of both training and target datasets. The model is then fine-tuned by re-training top layers on the specific dataset in order to re-learn domain-specific high level concepts (e.g. fine-tuning for radiology image classification [57]). Fine-tuning on specific biological datasets enables more focused predictions.

In genomics, the Basset package [227] for predicting chromatin accessibility was shown to rapidly learn and accurately predict on new data by leveraging a model pre-trained on available public data. To simulate this scenario, authors put aside 15 of 164 cell type datasets and trained the Basset model on the remaining 149 datasets. Then, they fine-tuned the model with one training pass of each of the remaining datasets and achieved results close to the model trained on all 164 datasets together. In another example, Min et al. [228] demonstrated how training on the experimentally-validated FANTOM5 permissive enhancer dataset followed by fine-tuning on ENCODE enhancer datasets improved cell type-specific predictions, outperforming state-of-the-art results. In drug design, general RNN models trained to generate molecules from the ChEMBL database have been fine-tuned to produce drug-like compounds for specific targets [420, 423].

Related to transfer learning, multimodal learning assumes simultaneous learning from various types of inputs, such as images and text. It can capture features that describe common concepts across input modalities. Generative graphical models like RBMs, deep Boltzmann machines, and DBNs, demonstrate successful extraction of more informative features for one modality (images or video) when jointly learned with other modalities (audio or text) [538]. Deep graphical models such as DBNs are well-suited for multimodal learning tasks because they learn a joint probability distribution from inputs. They can be pre-trained in an unsupervised fashion on large unlabeled data and then fine-tuned on a smaller number of labeled examples. When labels are available, convolutional neural networks are ubiquitously used because they can be trained end-to-end with backpropagation and demonstrate state-of-the-art performance in many discriminative tasks [14].

Jha et al. [190] showed that integrated training delivered better performance than individual networks. They compared a number of feed-forward architectures trained on RNA-seq data with and without an additional set of CLIP-seq, knockdown, and over-expression based input features. The integrative deep model generalized well for combined data, offering a large performance improvement for alternative splicing event estimation. Chaudhary et al. [539] trained a deep autoencoder model jointly on RNA-seq, miRNA-seq, and methylation data from TCGA to predict survival subgroups of hepatocellular carcinoma patients. This multimodal approach that treated different omic data types as different modalities outperformed both traditional methods (principal component analysis) and single-omic models. Interestingly, multi-omic model performance did not improve when combined with clinical information, suggesting that the model was able to capture redundant contributions of clinical features through their correlated genomic features. Chen et al. [176] used deep belief networks to learn phosphorylation states of a common set of signaling proteins in primary cultured bronchial cells collected from rats and humans treated with distinct stimuli. By interpreting species as different modalities representing similar high-level concepts, they showed that DBNs were able to capture cross-species representation of signaling mechanisms in response to a common stimuli. Another application used DBNs for joint unsupervised feature learning from cancer datasets containing gene expression, DNA methylation, and miRNA expression data [184]. This approach allowed for the capture of intrinsic relationships in different modalities and for better clustering performance over conventional k-means.

Multimodal learning with CNNs is usually implemented as a collection of individual networks in which each learns representations from single data type. These individual representations are further concatenated before or within fully-connected layers. FIDDLE [540] is an example of a multimodal CNN that represents an ensemble of individual networks that take NET-seq, MNase-seq, ChIP-seq, RNA-seq, and raw DNA sequence as input to predict transcription start sites. The combined model radically improves performance over separately trained datatype-specific networks, suggesting that it learns the synergistic relationship between datasets.

Multi-task learning is an approach related to transfer learning. In a multi-task learning framework, a model learns a number of tasks simultaneously such that features are shared across them. DeepSEA [209] implemented multi-task joint learning of diverse chromatin factors from raw DNA sequence. This allowed a sequence feature that was effective in recognizing binding of a specific TF to be simultaneously used by another predictor for a physically interacting TF. Similarly, TFImpute [191] learned information shared across transcription factors and cell lines to predict cell-specific TF binding for TF-cell line combinations. Yoon et al. [101] demonstrated that predicting the primary cancer site from cancer pathology reports together with its laterality substantially improved the performance for the latter task, indicating that multi-task learning can effectively leverage the commonality between two tasks using a shared representation. Many studies employed multi-task learning to predict chemical bioactivity [387, 391] and drug toxicity [392, 541]. Kearnes et al. [385] systematically compared single-task and multi-task models for ADMET properties and found that multi-task learning generally improved performance. Smaller datasets tended to benefit more than larger datasets.

Multi-task learning is complementary to multimodal and transfer learning. All three techniques can be used together in the same model. For example, Zhang et al. [534] combined deep model-based transfer and multi-task learning for cross-domain image annotation. One could imagine extending that approach to multimodal inputs as well. A common characteristic of these methods is better generalization of extracted features at various hierarchical levels of abstraction, which is attained by leveraging relationships between various inputs and task objectives.

Despite demonstrated improvements, transfer learning approaches pose challenges. There are no theoretically sound principles for pre-training and fine-tuning. Best practice recommendations are heuristic and must account for additional hyper-parameters that depend on specific deep architectures, sizes of the pre-training and target datasets, and similarity of domains. However, similarity of datasets and domains in transfer learning and relatedness of tasks in multi-task learning is difficult to access. Most studies address these limitations by empirical evaluation of the model. Unfortunately, negative results are typically not reported. A deep CNN trained on natural images boosts performance in radiographic images [57]. However, due to differences in imaging domains, the target task required either re-training the initial model from scratch with special preprocessing or fine-tuning of the whole network on radiographs with heavy data augmentation to avoid overfitting. Exclusively fine-tuning top layers led to much lower validation accuracy (81.4 versus 99.5). Fine-tuning the aforementioned Basset model with more than one pass resulted in overfitting [227]. DeepChem successfully improved results for low-data drug discovery with one-shot learning for related tasks. However, it clearly demonstrated the limitations of cross-task generalization across unrelated tasks in one-shot models, specifically nuclear receptor assays and patient adverse reactions [403].

In the medical domain, multimodal, multi-task and transfer learning strategies not only inherit most methodological issues from natural image, text, and audio domains, but also pose domain-specific challenges. There is a compelling need for the development of privacy-preserving transfer learning algorithms, such as Private Aggregation of Teacher Ensembles [158]. We suggest that these types of models deserve deeper investigation to establish sound theoretical guarantees and determine limits for the transferability of features between various closely related and distant learning tasks.

## Conclusions

Deep learning-based methods now match or surpass the previous state of the art in a diverse array of tasks in patient and disease categorization, fundamental biological study, genomics, and treatment development. Returning to our central question: given this rapid progress, has deep learning transformed the study of human disease? Though the answer is highly dependent on the specific domain and problem being addressed, we conclude that deep learning has not yet realized its transformative potential or induced a strategic inflection point. Despite its dominance over competing machine learning approaches in many of the areas reviewed here and quantitative improvements in predictive performance, deep learning has not yet definitively “solved” these problems.

As an analogy, consider recent progress in conversational speech recognition. Since 2009 there have been drastic performance improvements with error rates dropping from more than 20% to less than 6% [542] and finally approaching or exceeding human performance in the past year [543, 544]. The phenomenal improvements on benchmark datasets are undeniable, but greatly reducing the error rate on these benchmarks did not fundamentally transform the domain. Widespread adoption of conversational speech technologies will require solving the problem, i.e. methods that surpass human performance, and persuading users to adopt them [542]. We see parallels in healthcare, where achieving the full potential of deep learning will require outstanding predictive performance as well as acceptance and adoption by biologists and clinicians. These experts will rightfully demand rigorous evidence that deep learning has impacted their respective disciplines—elucidated new biological mechanisms and improved patient outcomes—to be convinced that the promises of deep learning are more substantive than those of previous generations of artificial intelligence.

Some of the areas we have discussed are closer to surpassing this lofty bar than others, generally those that are more similar to the non-biomedical tasks that are now monopolized by deep learning. In medical imaging, diabetic retinopathy [49], diabetic macular edema [49], tuberculosis [58], and skin lesion [4] classifiers are highly accurate and comparable to clinician performance.

In other domains, perfect accuracy will not be required because deep learning will primarily prioritize experiments and assist discovery. For example, in chemical screening for drug discovery, a deep learning system that successfully identifies dozens or hundreds of target-specific, active small molecules from a massive search space would have immense practical value even if its overall precision is modest. In medical imaging, deep learning can point an expert to the most challenging cases that require manual review [58], though the risk of false negatives must be addressed. In protein structure prediction, errors in individual residue-residue contacts can be tolerated when using the contacts jointly for 3D structure modeling. Improved contact map predictions [28] have led to notable improvements in fold and 3D structure prediction for some of the most challenging proteins, such as membrane proteins [250].

Conversely, the most challenging tasks may be those in which predictions are used directly for downstream modeling or decision-making, especially in the clinic. As an example, errors in sequence variant calling will be amplified if they are used directly for GWAS. In addition, the stochasticity and complexity of biological systems implies that for some problems, for instance predicting gene regulation in disease, perfect accuracy will be unattainable.

We are witnessing deep learning models achieving human-level performance across a number of biomedical domains. However, machine learning algorithms, including deep neural networks, are also prone to mistakes that humans are much less likely to make, such as misclassification of adversarial examples [545, 546], a reminder that these algorithms do not understand the semantics of the objects presented. It may be impossible to guarantee that a model is not susceptible to adversarial examples, but work in this area is continuing [547, 548]. Cooperation between human experts and deep learning algorithms addresses many of these challenges and can achieve better performance than either individually [64]. For sample and patient classification tasks, we expect deep learning methods to augment clinicians and biomedical researchers.

We are optimistic about the future of deep learning in biology and medicine. It is by no means inevitable that deep learning will revolutionize these domains, but given how rapidly the field is evolving, we are confident that its full potential in biomedicine has not been explored. We have highlighted numerous challenges beyond improving training and predictive accuracy, such as preserving patient privacy and interpreting models. Ongoing research has begun to address these problems and shown that they are not insurmountable. Deep learning offers the flexibility to model data in its most natural form, for example, longer DNA sequences instead of k-mers for transcription factor binding prediction and molecular graphs instead of pre-computed bit vectors for drug discovery. These flexible input feature representations have spurred creative modeling approaches that would be infeasible with other machine learning techniques. Unsupervised methods are currently less-developed than their supervised counterparts, but they may have the most potential because of how expensive and time-consuming it is to label large amounts of biomedical data. If future deep learning algorithms can summarize very large collections of input data into interpretable models that spur scientists to ask questions that they did not know how to ask, it will be clear that deep learning has transformed biology and medicine.

## Methods

### Continuous collaborative manuscript drafting

We recognized that deep learning in precision medicine is a rapidly developing area. Hence, diverse expertise was required to provide a forward-looking perspective. Accordingly, we collaboratively wrote this review in the open, enabling anyone with expertise to contribute. We wrote the manuscript in markdown and tracked changes using git. Contributions were handled through GitHub, with individuals submitting “pull requests” to suggest additions to the manuscript.

To facilitate citation, we defined a markdown citation syntax. We supported citations to the following identifier types (in order of preference): DOIs, PubMed Central IDs, PubMed IDs, arXiv IDs, and URLs. References were automatically generated from citation metadata by querying APIs to generate Citation Style Language (CSL) JSON items for each reference. Pandoc and pandocciteproc converted the markdown to HTML and PDF, while rendering the formatted citations and references. In total, referenced works consisted of 369 DOIs, 6 PubMed Central records, 129 arXiv manuscripts, and 48 URLs (webpages as well as manuscripts lacking standardized identifiers).

We implemented continuous analysis so the manuscript was automatically regenerated whenever
the source changed [147]. We configured Travis CI—a continuous integration service—to fetch new citation metadata and rebuild the manuscript for every commit. Accordingly, formatting or citation errors in pull requests would cause the Travis CI build to fail, automating quality control. In addition, the build process renders templated variables, such as the reference counts mentioned above, to automate the updating of dynamic content. When contributions were merged into the master branch, Travis CI deployed the built manuscript by committing back to the GitHub repository. As a result, the latest manuscript version is always available at https://greenelab.github.io/deep-review. To ensure a consistent software environment, we defined a versioned conda environment of the software dependencies

In addition, we instructed the Travis CI deployment script to perform blockchain timestamping [549,550]. Using OpenTimestamps, we submitted hashes for the manuscript and the source git commit for timestamping in the Bitcoin blockchain [551]. These timestamps attest that a given version of this manuscript (and its history) existed at a given point in time. The ability to irrefutably prove manuscript existence at a past time could be important to establish scientific precedence and enforce an immutable record of authorship.

## Author contributions

We created an open repository on the GitHub version control platform `greenelab/deep-review`) [552]. Here, we engaged with numerous authors from papers within and outside of the area. The manuscript was drafted via GitHub commits by 36 individuals who met the ICMJE standards of authorship. These were individuals who contributed to the review of the literature; drafted the manuscript or provided substantial critical revisions; approved the final manuscript draft; and agreed to be accountable in all aspects of the work. Individuals who did not contribute in all of these ways, but who did participate, are acknowledged below. We grouped authors into the following four classes of approximately equal contributions and randomly ordered authors within each contribution class. Drafted multiple sub-sections along with extensive editing, pull request reviews, or discussion: A.A.K., B.K.B., B.T.D., D.S.H., E.F., G.P.W., M.M.H., M.Z., P.A., T.C. Drafted one or more sub-sections: A.E.C., A.M.A., A.S., B.J.L., C.A.L., E.M.C., G.L.R., J.I., J.L, J.X., S.C.T., S.W., W.X., Z.L. Revised specific sub-sections or supervised drafting one or more subsections: A.H., A.K., D.D., D.J.H., L.K.W., M.H.S.S., S.J.S., S.M.B., Y.P, Y.Q. Drafted sub-sections, edited the manuscript, reviewed pull requests, and coordinated co-authors: A.G., C.S.G.

## Acknowledgements

We gratefully acknowledge Christof Angermueller, Kumardeep Chaudhary, Gökcen Eraslan, Mikael Huss, Bharath Ramsundar and Xun Zhu for their discussion of the manuscript and reviewed papers on GitHub. We would like to thank Aaron Sheldon, who contributed text but did not formally approve the manuscript. We would like to thank Anna Greene for a careful proofreading of the manuscript in advance of the first submission. We would like to thank Robert Gieseke, Ruibang Luo, Sourav Singh, and GitHub user snikumbh for correcting typos, formatting, and references. Finally, we acknowledge funding from the Gordon and Betty Moore Foundation awards GBMF4552 (C.S.G. and D.S.H.) and GBMF4563 (D.J.H.); the Howard Hughes Medical Institute (S.C.T.); the National Institutes of Health awards DP2GM123485 (A.K.), P30CA051008 (S.M.B.), R01AI116794 (B.K.B.), R01GM089652 (A.E.C.), R01GM089753 (J.X.), R01LM012222 (S.J.S.), R01LM012482 (S.J.S.), R21CA220398 (S.M.B.), T32GM007753 (B.T.D.), T32HG000046 (G.P.W.), and U54AI117924 (A.G.); the National Institutes of Health Intramural Research Program and National Library of Medicine (Y.P. and Z.L.); the National Science Foundation awards 1245632 (G.L.R.), 1531594 (E.M.C.), and 1564955 (J.X.); the Natural Sciences and Engineering Research Council of Canada award RGPIN-2015-3948 (M.M.H.); and the Roy and Diana Vagelos Scholars Program in the Molecular Life Sciences (M.Z.).

## References

1. Big Data: Astronomical or Genomical? Zachary D. Stephens, Skylar Y. Lee, Faraz Faghri, Roy H. Campbell, Chengxiang Zhai, Miles J. Efron, Ravishankar Iyer, Michael C. Schatz, Saurabh Sinha, Gene E. Robinson PLOS Biology (2015-07-07) https://doi.org/10.1371/journal.pbio.1002195

2. Deep learning Yann LeCun, Yoshua Bengio, Geoffrey Hinton Nature (2015-05-27) https://doi.org/10.1038/nature14539

3. Searching for exotic particles in high-energy physics with deep learning P. Baldi, P. Sadowski, D. Whiteson Nature Communications (2014-07-02) https://doi.org/10.1038/ncomms5308

4. Dermatologist-level classification of skin cancer with deep neural networks Andre Esteva, Brett Kuprel, Roberto A. Novoa, Justin Ko, Susan M. Swetter, Helen M. Blau, Sebastian Thrun Nature (2017-01-25) https://doi.org/10.1038/nature21056

5. Google’s Neural Machine Translation System: Bridging the Gap between Human and Machine Translation Yonghui Wu, Mike Schuster, Zhifeng Chen, Quoc V. Le, Mohammad Norouzi, Wolfgang Macherey, Maxim Krikun, Yuan Cao, Qin Gao, Klaus Macherey, … Jeffrey Dean arXiv (2016-09-26) https://arxiv.org/abs/1609.08144v2

6. A logical calculus of the ideas immanent in nervous activity Warren S. McCulloch, Walter Pitts The Bulletin of Mathematical Biophysics (1943-12) https://doi.org/10.1007/bf02478259

7. Analysis of a Four-Layer Series-Coupled Perceptron. II H. D. Block, B. W. Knight, F. Rosenblatt Reviews of Modern Physics (1962-01-01) https://doi.org/10.1103/revmodphys.34.135

8. Google Research Publication: Building High-level Features Using Large Scale Unsupervised Learning(2016-12-15) http://research.google.com/archive/unsupervised_icml2012.html

9. HOGWILD!: A Lock-Free Approach to Parallelizing Stochastic Gradient Descent Feng Niu, Benjamin Recht, Christopher Re, Stephen J. Wright arXiv (2011-06-28) https://arxiv.org/abs/1106.5730v2

10. Deep Learning Ian Goodfellow, Yoshua Bengio, Aaron Courville (2016) http://www.deeplearningbook.org/

11. Academy of Management: Andrew S. Grove Andrew S. Grove (1998-08-09) http://www.intel.com/pressroom/archive/speeches/ag080998.htm

12. Deep learning for regulatory genomics Yongjin Park, Manolis Kellis Nature Biotechnology (2015-08-07) https://doi.org/10.1038/nbt.3313

13. Applications of Deep Learning in Biomedicine Polina Mamoshina, Armando Vieira, Evgeny Putin, Alex Zhavoronkov Molecular Pharmaceutics (2016-05-02) https://doi.org/10.1021/acs.molpharmaceut.5b00982

14. Deep learning for computational biology Christof Angermueller, Tanel Pärnamaa, Leopold Parts, Oliver Stegle Molecular Systems Biology (2016-07) https://doi.org/10.15252/msb.20156651

15. Deep learning in bioinformatics Seonwoo Min, Byunghan Lee, Sungroh Yoon Briefings in Bioinformatics (2016-07-29) https://doi.org/10.1093/bib/bbw068

16. Computer vision for high content screening Oren Z. Kraus, Brendan J. Frey Critical Reviews in Biochemistry and Molecular Biology (2016-01-24) https://doi.org/10.3109/10409238.2015.1135868

17. Deep learning for healthcare:review>, opportunities and challenges Riccardo Miotto, Fei Wang, Shuang Wang, Xiaoqian Jiang, Joel T. Dudley Briefings in Bioinformatics (2017-05-06) https://doi.org/10.1093/bib/bbx044

18. A survey on deep learning in medical image analysis Geert Litjens, Thijs Kooi, Babak Ehteshami Bejnordi, Arnaud Arindra Adiyoso Setio, Francesco Ciompi, Mohsen Ghafoorian, Jeroen A.W.M. van der Laak, Bram van Ginneken, Clara I. Sánchez Medical Image Analysis (2017-12) https://doi.org/10.1016/j.media.2017.07.005

19. Deep Learning in Drug Discovery Erik Gawehn, Jan A. Hiss, Gisbert Schneider Molecular Informatics (2015-12-30) https://doi.org/10.1002/minf.201501008

20. Deep learning for computational chemistry Garrett B. Goh, Nathan O. Hodas, Abhinav Vishnu Journal of Computational Chemistry (2017-03-08) https://doi.org/10.1002/jcc.24764

21. Virtual Screening: A Challenge for Deep Learning Javier Pérez-Sianes, Horacio Pérez-Sánchez, Fernando Díaz Advances in Intelligent Systems and Computing (2016) https://doi.org/10.1007/978-3-319-40126-3_2

22. A renaissance of neural networks in drug discovery Igor I. Baskin, David Winkler, Igor V. Tetko Expert Opinion on Drug Discovery (2016-07-04) https://doi.org/10.1080/17460441.2016.1201262

23. Supervised Risk Predictor of Breast Cancer Based on Intrinsic Subtypes Joel S. Parker, Michael Mullins, Maggie C.U. Cheang, Samuel Leung, David Voduc, Tammi Vickery, Sherri Davies, Christiane Fauron, Xiaping He, Zhiyuan Hu, … Philip S. Bernard Journal of Clinical Oncology (2009-03-10) https://doi.org/10.1200/jco.2008.18.1370

24. New Strategies for Triple-Negative Breast Cancer–Deciphering the Heterogeneity I. A. Mayer, V. G. Abramson, B. D. Lehmann, J. A. Pietenpol Clinical Cancer Research (2014-02-15) https://doi.org/10.1158/1078-0432.ccr-13-0583

25. UNSUPERVISED FEATURE CONSTRUCTION AND KNOWLEDGE EXTRACTION FROM GENOME-WIDE ASSAYS OF BREAST CANCER WITH DENOISING AUTOENCODERS Jie Tan, Matthew Ung, Chao Cheng, Casey S Greene Biocomputing 2015 (2014-11) https://doi.org/10.1142/9789814644730_0014

26. Mitosis Detection in Breast Cancer Histology Images with Deep Neural Networks Dan C. Cireşan, Alessandro Giusti, Luca M. Gambardella, Jürgen Schmidhuber Medical Image Computing and Computer-Assisted Intervention – MICCAI 2013 (2013) https://doi.org/10.1007/978-3-642-40763-5_51

27. End effector target position learning using feedforward with error back-propagation and recurrent neural networks J. Zurada Proceedings of 1994 IEEE International Conference on Neural Networks (ICNN’94) (1994) https://doi.org/10.1109/icnn.1994.374637

28. Accurate De Novo Prediction of Protein Contact Map by Ultra-Deep Learning Model Sheng Wang, Siqi Sun, Zhen Li, Renyu Zhang, Jinbo Xu PLOS Computational Biology (2017-01-05) https://doi.org/10.1371/journal.pcbi.1005324

29. A Deep Learning Network Approach to ab initio Protein Secondary Structure Prediction Matt Spencer, Jesse Eickholt, Jianlin Cheng IEEE/ACM Transactions on Computational Biology and Bioinformatics (2015-01-01) https://doi.org/10.1109/tcbb.2014.2343960

30. Protein Secondary Structure Prediction Using Deep Convolutional Neural Fields Sheng Wang, Jian Peng, Jianzhu Ma, Jinbo Xu Scientific Reports (2016-01-11) https://doi.org/10.1038/srep18962

31. PEDLA: predicting enhancers with a deep learning-based algorithmic framework Feng Liu, Hao Li, Chao Ren, Xiaochen Bo, Wenjie Shu Cold Spring Harbor Laboratory (2016-01-07) https://doi.org/10.1101/036129

32. Deep Feature Selection: Theory and Application to Identify Enhancers and Promoters Yifeng Li, Chih-Yu Chen, Wyeth W. Wasserman Lecture Notes in Computer Science (2015) https://doi.org/10.1007/978-3-319-16706-0_20

33. DEEP: a general computational framework for predicting enhancers Dimitrios Kleftogiannis, Panos Kalnis, Vladimir B. Bajic Nucleic Acids Research (2014-11-05) https://doi.org/10.1093/nar/gku1058

34. DANN: a deep learning approach for annotating the pathogenicity of genetic variants Daniel Quang, Yifei Chen, Xiaohui Xie Bioinformatics (2014-10-22) https://doi.org/10.1093/bioinformatics/btu703

35. AtomNet: A Deep Convolutional Neural Network for Bioactivity Prediction in Structure-based Drug Discovery Izhar Wallach, Michael Dzamba, Abraham Heifets arXiv (2015-10-10) https://arxiv.org/abs/1510.02855v1

36. Deep Learning Applications for Predicting Pharmacological Properties of Drugs and Drug Repurposing Using Transcriptomic Data Alexander Aliper, Sergey Plis, Artem Artemov, Alvaro Ulloa, Polina Mamoshina, Alex Zhavoronkov Molecular Pharmaceutics (2016-07-05) https://doi.org/10.1021/acs.molpharmaceut.6b00248

37. Predicting drug-target interactions using restricted Boltzmann machines Yuhao Wang, Jianyang Zeng Bioinformatics (2013-06-19) https://doi.org/10.1093/bioinformatics/btt234

38. Deep-Learning-Based Drug–Target Interaction Prediction Ming Wen, Zhimin Zhang, Shaoyu Niu, Haozhi Sha, Ruihan Yang, Yonghuan Yun, Hongmei Lu Journal of Proteome Research (2017-03-13) https://doi.org/10.1021/acs.jproteome.6b00618

39. Deep Learning in Medical Image Analysis Dinggang Shen, Guorong Wu, Heung-Il Suk Annual Review of Biomedical Engineering (2017-06-21) https://doi.org/10.1146/annurev-bioeng-071516-044442

40. Deep Learning and Structured Prediction for the Segmentation of Mass in Mammograms Neeraj Dhungel, Gustavo Carneiro, Andrew P. Bradley Lecture Notes in Computer Science (2015) https://doi.org/10.1007/978-3-319-24553-9_74

41. The Automated Learning of Deep Features for Breast Mass Classification from Mammograms Neeraj Dhungel, Gustavo Carneiro, Andrew P. Bradley Medical Image Computing and Computer-Assisted Intervention – MICCAI 2016 (2016) https://doi.org/10.1007/978-3-319-46723-8_13

42. Deep Multi-instance Networks with Sparse Label Assignment for Whole Mammogram Classification Wentao Zhu, Qi Lou, Yeeleng Scott Vang, Xiaohui Xie Cold Spring Harbor Laboratory (2016-12-20) https://doi.org/10.1101/095794

43. Adversarial Deep Structural Networks for Mammographic Mass Segmentation Wentao Zhu, Xiaohui Xie Cold Spring Harbor Laboratory (2016-12-20) https://doi.org/10.1101/095786

44. A deep learning approach for the analysis of masses in mammograms with minimal user intervention Neeraj Dhungel, Gustavo Carneiro, Andrew P. Bradley Medical Image Analysis (2017-04) https://doi.org/10.1016/j.media.2017.01.009

45. ImageNet Large Scale Visual Recognition Challenge Olga Russakovsky, Jia Deng, Hao Su, Jonathan Krause, Sanjeev Satheesh, Sean Ma, Zhiheng Huang, Andrej Karpathy, Aditya Khosla, Michael Bernstein, … Li Fei-Fei International Journal of Computer Vision (2015-04-11) https://doi.org/10.1007/s11263-015-0816-y

46. Convolutional Neural Networks for Diabetic Retinopathy Harry Pratt, Frans Coenen, Deborah M. Broadbent, Simon P. Harding, Yalin Zheng Procedia Computer Science (2016) https://doi.org/10.1016/j.procs.2016.07.014

47. Improved Automated Detection of Diabetic Retinopathy on a Publicly Available Dataset Through Integration of Deep Learning Michael David Abràmoff, Yiyue Lou, Ali Erginay, Warren Clarida, Ryan Amelon, James C. Folk, Meindert Niemeijer Investigative Opthalmology & Visual Science (2016-10-04) https://doi.org/10.1167/iovs.16-19964

48. Leveraging uncertainty information from deep neural networks for disease detection Christian Leibig, Vaneeda Allken, Murat Seckin Ayhan, Philipp Berens, Siegfried Wahl Cold Spring Harbor Laboratory (2016-10-28) https://doi.org/10.1101/084210

49. Development and Validation of a Deep Learning Algorithm for Detection of Diabetic Retinopathy in Retinal Fundus Photographs Varun Gulshan, Lily Peng, Marc Coram, Martin C. Stumpe, Derek Wu, Arunachalam Narayanaswamy, Subhashini Venugopalan, Kasumi Widner, Tom Madams, Jorge Cuadros, … Dale R. Webster JAMA (2016-12-13) https://doi.org/10.1001/jama.2016.17216

50. Deep Learning Ensembles for Melanoma Recognition in Dermoscopy Images Noel Codella, Quoc-Bao Nguyen, Sharath Pankanti, David Gutman, Brian Helba, Allan Halpern, John R. Smith arXiv (2016-10-14) https://arxiv.org/abs/1610.04662v2

51. Automated Melanoma Recognition in Dermoscopy Images via Very Deep Residual Networks Lequan Yu, Hao Chen, Qi Dou, Jing Qin, Pheng-Ann Heng IEEE Transactions on Medical Imaging (2017-04) https://doi.org/10.1109/tmi.2016.2642839

52. Extraction of skin lesions from non-dermoscopic images for surgical excision of melanoma M. Hossein Jafari, Ebrahim Nasr-Esfahani, Nader Karimi, S. M. Reza Soroushmehr, Shadrokh Samavi, Kayvan Najarian International Journal of Computer Assisted Radiology and Surgery (2017-03-24) https://doi.org/10.1007/s11548-017-1567-8

53. Melanoma detection by analysis of clinical images using convolutional neural network E. Nasr-Esfahani, S. Samavi, N. Karimi, S.M.R. Soroushmehr, M.H. Jafari, K. Ward, K. Najarian 2016 38th Annual International Conference of the IEEE Engineering in Medicine and Biology Society (EMBC) (2016-08) https://doi.org/10.1109/embc.2016.7590963

54. Detection of age-related macular degeneration via deep learning P. Burlina, D. E. Freund, N. Joshi, Y. Wolfson, N. M. Bressler 2016 IEEE 13th International Symposium on Biomedical Imaging (ISBI) (2016-04) https://doi.org/10.1109/isbi.2016.7493240

55. Deep learning with non-medical training used for chest pathology identification Yaniv Bar, Idit Diamant, Lior Wolf, Hayit Greenspan Medical Imaging 2015: Computer-Aided Diagnosis (2015-03-20) https://doi.org/10.1117/12.2083124

56. Deep Convolutional Neural Networks for Computer-Aided Detection: CNN Architectures, Dataset Characteristics and Transfer Learning Hoo-Chang Shin, Holger R. Roth, Mingchen Gao, Le Lu, Ziyue Xu, Isabella Nogues, Jianhua Yao, Daniel Mollura, Ronald M. Summers IEEE Transactions on Medical Imaging (2016-05) https://doi.org/10.1109/tmi.2016.2528162

57. High-Throughput Classification of Radiographs Using Deep Convolutional Neural Networks Alvin Rajkomar, Sneha Lingam, Andrew G. Taylor, Michael Blum, John Mongan Journal of Digital Imaging (2016-10-11) https://doi.org/10.1007/s10278-016-9914-9

58. Deep Learning at Chest Radiography: Automated Classification of Pulmonary Tuberculosis by Using Convolutional Neural Networks Paras Lakhani, Baskaran Sundaram Radiology (2017-08) https://doi.org/10.1148/radiol.2017162326

59. Classification of breast MRI lesions using small-size training sets: comparison of deep learning approaches Guy Amit, Rami Ben-Ari, Omer Hadad, Einat Monovich, Noa Granot, Sharbell Hashoul Medical Imaging 2017: Computer-Aided Diagnosis (2017-03-03) https://doi.org/10.1117/12.2249981

60. Improving Computer-Aided Detection Using_newlineConvolutional Neural Networks and Random View Aggregation Holger R. Roth, Le Lu, Jiamin Liu, Jianhua Yao, Ari Seff, Kevin Cherry, Lauren Kim, Ronald M. Summers IEEE Transactions on Medical Imaging (2016-05) https://doi.org/10.1109/tmi.2015.2482920

61. 3D Deep Learning for Multi-modal Imaging-Guided Survival Time Prediction of Brain Tumor Patients Dong Nie, Han Zhang, Ehsan Adeli, Luyan Liu, Dinggang Shen Medical Image Computing and Computer-Assisted Intervention – MICCAI 2016 (2016) https://doi.org/10.1007/978-3-319-46723-8_25

62. Large scale deep learning for computer aided detection of mammographic lesions Thijs Kooi, Geert Litjens, Bram van Ginneken, Albert Gubern-Mérida, Clara I. Sánchez, Ritse Mann, Ard den Heeten, Nico Karssemeijer Medical Image Analysis (2017-01) https://doi.org/10.1016/j.media.2016.07.007

63. Deep learning as a tool for increased accuracy and efficiency of histopathological diagnosis Geert Litjens, Clara I. Sánchez, Nadya Timofeeva, Meyke Hermsen, Iris Nagtegaal, Iringo Kovacs, Christina Hulsbergen - van de Kaa, Peter Bult, Bram van Ginneken, Jeroen van der Laak Scientific Reports (2016-05-23) https://doi.org/10.1038/srep26286

64. Deep Learning for Identifying Metastatic Breast Cancer Dayong Wang, Aditya Khosla, Rishab Gargeya, Humayun Irshad, Andrew H. Beck arXiv (2016-06-18) https://arxiv.org/abs/1606.05718v1

65. Deep learning is effective for the classification of OCT images of normal versus Age-related Macular Degeneration Cecilia S Lee, Doug M Baughman, Aaron Y Lee Cold Spring Harbor Laboratory (2016-12-14) https://doi.org/10.1101/094276

66. ImageNet Classification with Deep Convolutional Neural Networks Alex Krizhevsky, Ilya Sutskever, Geoffrey E. Hinton Proceedings of the 25th International Conference on Neural Information Processing Systems (2012) http://dl.acm.org/citation.cfm?id=2999134.2999257

67. A shared task involving multi-label classification of clinical free text John P. Pestian, Christopher Brew, Paweł Matykiewicz, D. J. Hovermale, Neil Johnson, K. Bretonnel Cohen, Włodzisław Duch Proceedings of the Workshop on BioNLP 2007 Biological, Translational, and Clinical Language Processing - BioNLP’07 (2007) https://doi.org/10.3115/1572392.1572411

68. ChestX-ray8: Hospital-scale Chest X-ray Database and Benchmarks on Weakly-Supervised Classification and Localization of Common Thorax Diseases Xiaosong Wang, Yifan Peng, Le Lu, Zhiyong Lu, Mohammadhadi Bagheri, Ronald M. Summers arXiv (2017-05-05) https://arxiv.org/abs/1705.02315v4

69. NegBio: a high-performance tool for negation and uncertainty detection in radiology reports Yifan Peng, Xiaosong Wang, Le Lu, Mohammadhadi Bagheri, Ronald Summers, Zhiyong Lu arXiv (2017-12-16) https://arxiv.org/abs/1712.05898v2

70. Points of Significance: Classification evaluation Jake Lever, Martin Krzywinski, Naomi Altman Nature Methods (2016-07-28) https://doi.org/10.1038/nmeth.3945

71. NIH Chest X-ray Dataset NIH Clinical Center (2017-09-07) https://nihcc.app.box.com/v/ChestXray-NIHCC

72. TaggerOne: joint named entity recognition and normalization with semi-Markov Models Robert Leaman, Zhiyong Lu Bioinformatics (2016-06-09) https://doi.org/10.1093/bioinformatics/btw343

73. tmVar: a text mining approach for extracting sequence variants in biomedical literature C.-H. Wei, B. R. Harris, H.-Y. Kao, Z. Lu Bioinformatics (2013-04-05) https://doi.org/10.1093/bioinformatics/btt156

74. DNorm: disease name normalization with pairwise learning to rank R. Leaman, R. Islamaj Dogan, Z. Lu Bioinformatics (2013-08-21) https://doi.org/10.1093/bioinformatics/btt474

75. Effects of Semantic Features on Machine Learning-Based Drug Name Recognition Systems: Word Embeddings vs. Manually Constructed Dictionaries Shengyu Liu, Buzhou Tang, Qingcai Chen, Xiaolong Wang Information (2015-12-11) https://doi.org/10.3390/info6040848

76. Evaluating Word Representation Features in Biomedical Named Entity Recognition Tasks Buzhou Tang, Hongxin Cao, Xiaolong Wang, Qingcai Chen, Hua Xu BioMed Research International (2014) https://doi.org/10.1155/2014/240403

77. Clinical Abbreviation Disambiguation Using Neural Word Embeddings yonghui wu, Jun Xu, Yaoyun Zhang, Hua Xu Proceedings of BioNLP 15 (2015) https://doi.org/10.18653/v1/w15-3822

78. Exploiting Task-Oriented Resources to Learn Word Embeddings for Clinical Abbreviation Expansion Yue Liu, Tao Ge, Kusum Mathews, Heng Ji, Deborah McGuinness Proceedings of BioNLP 15 (2015) https://doi.org/10.18653/v1/w15-3810

79. A Comprehensive Benchmark of Kernel Methods to Extract Protein–Protein Interactions from Literature Domonkos Tikk, Philippe Thomas, Peter Palaga, Jörg Hakenberg, Ulf Leser PLoS Computational Biology (2010-07-01) https://doi.org/10.1371/journal.pcbi.1000837

80. Improving chemical disease relation extraction with rich features and weakly labeled data Yifan Peng, Chih-Hsuan Wei, Zhiyong Lu Journal of Cheminformatics (2016-10-07) https://doi.org/10.1186/s13321-016-0165-z

81. Evaluation of linguistic features useful in extraction of interactions from PubMed; Application to annotating known, high-throughput and predicted interactions in I2D Yun Niu, David Otasek, Igor Jurisica Bioinformatics (2009-10-22) https://doi.org/10.1093/bioinformatics/btp602

82. Joint Models for Extracting Adverse Drug Events from Biomedical Text Fei Li, Yue Zhang, Meishan Zhang, Donghong Ji Proceedings of the Twenty-Fifth International Joint Conference on Artificial Intelligence (2016) http://dl.acm.org/citation.cfm?id=3060832.3061018

83. A neural joint model for entity and relation extraction from biomedical text Fei Li, Meishan Zhang, Guohong Fu, Donghong Ji BMC Bioinformatics (2017-03-31) https://doi.org/10.1186/s12859-017-1609-9

84. Deep learning for extracting protein-protein interactions from biomedical literature Yifan Peng, Zhiyong Lu BioNLP 2017 (2017) https://doi.org/10.18653/v1/w17-2304

85. A Shortest Dependency Path Based Convolutional Neural Network for Protein-Protein Relation Extraction Lei Hua, Chanqin Quan BioMed Research International (2016) https://doi.org/10.1155/2016/8479587

86. Multichannel Convolutional Neural Network for Biological Relation Extraction Chanqin Quan, Lei Hua, Xiao Sun, Wenjun Bai BioMed Research International (2016) https://doi.org/10.1155/2016/1850404

87. A general protein-protein interaction extraction architecture based on word representation and feature selection Zhenchao Jiang, shuang Li, Degen Huang International Journal of Data Mining and Bioinformatics (2016) https://doi.org/10.1504/ijdmb.2016.074878

88. Chemical-induced disease relation extraction via convolutional neural network Jinghang Gu, Fuqing Sun, Longhua Qian, Guodong Zhou Database (2017-01) https://doi.org/10.1093/database/bax024

89. Drug drug interaction extraction from biomedical literature using syntax convolutional neural network Zhehuan Zhao, Zhihao Yang, Ling Luo, Hongfei Lin, Jian Wang Bioinformatics (2016-07-27) https://doi.org/10.1093/bioinformatics/btw486

90. Extracting Drug-Drug Interactions with Attention CNNs Masaki Asada, Makoto Miwa, Yutaka Sasaki BioNLP 2017 (2017) https://doi.org/10.18653/v1/w17-2302

91. Drug-drug Interaction Extraction via Recurrent Neural Network with Multiple Attention Layers Zibo Yi, Shasha Li, Jie Yu, Qingbo Wu arXiv (2017-05-09) https://arxiv.org/abs/1705.03261v2

92. DUTIR in BioNLP-ST 2016: Utilizing Convolutional Network and Distributed Representation to Extract Complicate Relations Honglei Li, Jianhai Zhang, Jian Wang, Hongfei Lin, Zhihao Yang Proceedings of the 4th BioNLP Shared Task Workshop (2016) https://doi.org/10.18653/v1/w16-3012

93. Deep Learning with Minimal Training Data: TurkuNLP Entry in the BioNLP Shared Task 2016 Farrokh Mehryary, Jari Björne, Sampo Pyysalo, Tapio Salakoski, Filip Ginter Proceedings of the 4th BioNLP Shared Task Workshop (2016) https://doi.org/10.18653/v1/w16-3009

94. Using word embedding for bio-event extraction Chen Li, Runqing Song, Maria Liakata, Andreas Vlachos, Stephanie Seneff, Xiangrong Zhang Proceedings of BioNLP 15 (2015) https://doi.org/10.18653/v1/w15-3814

95. Embedding assisted prediction architecture for event trigger identification Yifan Nie, Wenge Rong, Yiyuan Zhang, Yuanxin Ouyang, Zhang Xiong Journal of Bioinformatics and Computational Biology (2015-06) https://doi.org/10.1142/s0219720015410012

96. Biomedical Event Trigger Identification Using Bidirectional Recurrent Neural Network Based Models Patchigolla V S S Rahul, Sunil Kumar Sahu, Ashish Anand arXiv (2017-05-26) https://arxiv.org/abs/1705.09516v1

97. Deep Learning for Biomedical Information Retrieval: Learning Textual Relevance from Click Logs Sunil Mohan, Nicolas Fiorini, Sun Kim, Zhiyong Lu BioNLP 2017 (2017) https://doi.org/10.18653/v1/w17-2328

98. Realizing the full potential of electronic health records: the role of natural language processing Lucila Ohno-Machado Journal of the American Medical Informatics Association (2011-09) https://doi.org/10.1136/amiajnl-2011-000501

99. Machine-learned solutions for three stages of clinical information extraction: the state of the art at i2b2 2010 Berry de Bruijn, Colin Cherry, Svetlana Kiritchenko, Joel Martin, Xiaodan Zhu Journal of the American Medical Informatics Association (2011-09) https://doi.org/10.1136/amiajnl-2011-000150

100. Bidirectional LSTM-CRF for Clinical Concept Extraction Raghavendra Chalapathy, Ehsan Zare Borzeshi, Massimo Piccardi arXiv (2016-11-25) https://arxiv.org/abs/1611.08373v1

101. Multi-task Deep Neural Networks for Automated Extraction of Primary Site and Laterality Information from Cancer Pathology Reports Hong-Jun Yoon, Arvind Ramanathan, Georgia Tourassi Advances in Big Data (2016-10-08) https://doi.org/10.1007/978-3-319-47898-2_21

102. Efficient Estimation of Word Representations in Vector Space Tomas Mikolov, Kai Chen, Greg Corrado, Jeffrey Dean arXiv (2013-01-16) https://arxiv.org/abs/1301.3781v3

103. Exploring the Application of Deep Learning Techniques on Medical Text Corpora Minarro-Giménez José Antonio, Marín-Alonso Oscar, Samwald Matthias Studies in Health Technology and Informatics (2014) https://doi.org/10.3233/978-1-61499-432-9-584

104. Medical Semantic Similarity with a Neural Language Model Lance De Vine, Guido Zuccon, Bevan Koopman, Laurianne Sitbon, Peter Bruza Proceedings of the 23rd ACM International Conference on Conference on Information and Knowledge Management-CIKM’14 (2014) https://doi.org/10.1145/2661829.2661974

105. Automatic Diagnosis Coding of Radiology Reports: A Comparison of Deep Learning and Conventional Classification Methods Sarvnaz Karimi, Xiang Dai, Hamedh Hassanzadeh, Anthony Nguyen BioNLP 2017 (2017) https://doi.org/10.18653/v1/w17-2342

106. International Classification of Diseases(2017-07-07) http://www.who.int/classifications/icd/en/

107. Multi-layer Representation Learning for Medical Concepts Edward Choi, Mohammad Taha Bahadori, Elizabeth Searles, Catherine Coffey, Michael Thompson, James Bost, Javier Tejedor-Sojo, Jimeng Sun Proceedings of the 22nd ACM SIGKDD International Conference on Knowledge Discovery and Data Mining - KDD’16 (2016) https://doi.org/10.1145/2939672.2939823

108. Large-Scale Discovery of Disease-Disease and Disease-Gene Associations Djordje Gligorijevic, Jelena Stojanovic, Nemanja Djuric, Vladan Radosavljevic, Mihajlo Grbovic, Rob J. Kulathinal, Zoran Obradovic Scientific Reports (2016-08-31) https://doi.org/10.1038/srep32404

109. Bidirectional RNN for Medical Event Detection in Electronic Health Records Abhyuday N Jagannatha, Hong Yu Proceedings of the conference. Association for Computational Linguistics. North American Chapter. Meeting (2016-06) https://www.ncbi.nlm.nih.gov/pmc/articles/PMC5119627/

110. Representations of Time Expressions for Temporal Relation Extraction with Convolutional Neural Networks Chen Lin, Timothy Miller, Dmitriy Dligach, Steven Bethard, Guergana Savova BioNLP 2017 (2017) https://doi.org/10.18653/v1/w17-2341

111. Computational Phenotype Discovery Using Unsupervised Feature Learning over Noisy, Sparse, and Irregular Clinical Data Thomas A. Lasko, Joshua C. Denny, Mia A. Levy PLoS ONE (2013-06-24) https://doi.org/10.1371/journal.pone.0066341

112. Semi-supervised learning of the electronic health record for phenotype stratification Brett K. Beaulieu-Jones, Casey S. Greene Journal of Biomedical Informatics (2016-12) https://doi.org/10.1016/j.jbi.2016.10.007

113. Deep Patient: An Unsupervised Representation to Predict the Future of Patients from the Electronic Health Records Riccardo Miotto, Li Li, Brian A. Kidd, Joel T. Dudley Scientific Reports (2016-05-17) https://doi.org/10.1038/srep26094

114. Doctor AI: Predicting Clinical Events via Recurrent Neural Networks Edward Choi, Mohammad Taha Bahadori, Andy Schuetz, Walter F. Stewart, Jimeng Sun arXiv (2015-11-18) https://arxiv.org/abs/1511.05942v11

115. DeepCare: A Deep Dynamic Memory Model for Predictive Medicine Trang Pham, Truyen Tran, Dinh Phung, Svetha Venkatesh arXiv (2016-02-01) https://arxiv.org/abs/1602.00357v2

116. Deepr: A Convolutional Net for Medical Records Phuoc Nguyen, Truyen Tran, Nilmini Wickramasinghe, Svetha Venkatesh IEEE Journal of Biomedical and Health Informatics (2017-01) https://doi.org/10.1109/jbhi.2016.2633963

117. Multi-task Prediction of Disease Onsets from Longitudinal Lab Tests Narges Razavian, Jake Marcus, David Sontag arXiv (2016-08-02) https://arxiv.org/abs/1608.00647v3

118. Deep Survival Analysis Rajesh Ranganath, Adler Perotte, Noémie Elhadad, David Blei arXiv (2016-08-06) https://arxiv.org/abs/1608.02158v2

119. Comparison of the performance of neural network methods and Cox regression for censored survival data Anny Xiang, Pablo Lapuerta, Alex Ryutov, Jonathan Buckley, Stanley Azen Computational Statistics & Data Analysis (2000-08) https://doi.org/10.1016/s0167-9473(99)00098-5

120. DeepSurv: Personalized Treatment Recommender System Using A Cox Proportional Hazards Deep Neural Network Jared Katzman, Uri Shaham, Jonathan Bates, Alexander Cloninger, Tingting Jiang, Yuval Kluger arXiv (2016-06-02) https://arxiv.org/abs/1606.00931v3

121. Deep Exponential Families Rajesh Ranganath, Linpeng Tang, Laurent Charlin, David M. Blei arXiv (2014-11-10) https://arxiv.org/abs/1411.2581v1

122. Stochastic Variational Inference Matt Hoffman, David M. Blei, Chong Wang, John Paisley arXiv (2012-06-29) https://arxiv.org/abs/1206.7051v3

123. Hierarchical Variational Models Rajesh Ranganath, Dustin Tran, David M. Blei arXiv (2015-11-07) https://arxiv.org/abs/1511.02386v2

124. A machine learning-based framework to identify type 2 diabetes through electronic health records Tao Zheng, Wei Xie, Liling Xu, Xiaoying He, Ya Zhang, Mingrong You, Gong Yang, You Chen International Journal of Medical Informatics (2017-01) https://doi.org/10.1016/j.ijmedinf.2016.09.014

125. Implementations by Phenotype | PheKB(2017-05-17) https://phekb.org/implementations

126. Electronic medical record phenotyping using the anchor and learn framework Yoni Halpern, Steven Horng, Youngduck Choi, David Sontag Journal of the American Medical Informatics Association (2016-04-23) https://doi.org/10.1093/jamia/ocw011

127. Data Programming: Creating Large Training Sets, Quickly Alexander Ratner, Christopher De Sa, Sen Wu, Daniel Selsam, Christopher Ré arXiv (2016-05-25) https://arxiv.org/abs/1605.07723v3

128. Data is the New Oil Michael Palmer ANA Marketing Maestros (2006-11) http://ana.blogs.com/maestros/2006/11/data_is_the_new.html

129. “Data is the New Oil” — A Ludicrous Proposition – Twenty One Hundred – Medium Michael Haupt Medium (2016-05-02) https://medium.com/twenty-one-hundred/data-is-the-new-oil-a-ludicrous-proposition-1d91bba4f294

130. Data Programming: Machine Learning with Weak Supervision Alex Ratner, Stephen Bach, Chris Ré (2016-09-19) http://hazyresearch.github.io/snorkel/blog/weak_supervision.html

131. Mining electronic health records: towards better research applications and clinical care Peter B. Jensen, Lars J. Jensen, Søren Brunak Nature Reviews Genetics (2012-05-02) https://doi.org/10.1038/nrg3208

132. Methods and dimensions of electronic health record data quality assessment: enabling reuse for clinical research N. G. Weiskopf, C. Weng Journal of the American Medical Informatics Association (2013-01-01) https://doi.org/10.1136/amiajnl-2011-000681

133. Impact of Electronic Health Record Systems on Information Integrity: Quality and Safety Implications Sue Bowman Perspectives in Health Information Management (2013) https://www.ncbi.nlm.nih.gov/pmc/articles/PMC3797550/

134. Secondary Use of EHR: Data Quality Issues and Informatics Opportunities Taxiarchis Botsis, Gunnar Hartvigsen, Fei Chen, Chunhua Weng Summit on Translational Bioinformatics (2010) https://www.ncbi.nlm.nih.gov/pmc/articles/PMC3041534/

135. Have DRG-based prospective payment systems influenced the number of secondary diagnoses in health care administrative data? Lisbeth Serdén, Rikard Lindqvist, Måns Rosén Health Policy (2003-08) https://doi.org/10.1016/s0168-8510(02)00208-7

136. Why Patient Matching Is a Challenge: Research on Master Patient Index (MPI) Data Discrepancies in Key Identifying Fields Beth Haenke Just, David Marc, Megan Munns, Ryan Sandefer Perspectives in Health Information Management (2016) https://www.ncbi.nlm.nih.gov/pmc/articles/PMC4832129/

137. Identifying and mitigating biases in EHR laboratory tests Rimma Pivovarov, David J. Albers, Jorge L. Sepulveda, Noémie Elhadad Journal of Biomedical Informatics (2014-10) https://doi.org/10.1016/j.jbi.2014.03.016

138. Using electronic health records for clinical research: The case of the EHR4CR project Georges De Moor, Mats Sundgren, Dipak Kalra, Andreas Schmidt, Martin Dugas, Brecht Claerhout, Töresin Karakoyun, Christian Ohmann, Pierre-Yves Lastic, Nadir Ammour, … Pascal Coorevits Journal of Biomedical Informatics (2015-02) https://doi.org/10.1016/j.jbi.2014.10.006

139. Healthcare Interoperability Standards Compliance Handbook Frank Oemig, Robert Snelick Springer International Publishing (2016) https://doi.org/10.1007/978-3-319-44839-8

140. How sample size influences research outcomes Jorge Faber, Lilian Martins Fonseca Dental Press Journal of Orthodontics (2014-08) https://doi.org/10.1590/2176-9451.19.4.027-029.ebo

141. A review of approaches to identifying patient phenotype cohorts using electronic health records Chaitanya Shivade, Preethi Raghavan, Eric Fosler-Lussier, Peter J Embi, Noemie Elhadad, Stephen B Johnson, Albert M Lai Journal of the American Medical Informatics Association (2014-03) https://doi.org/10.1136/amiajnl-2013-001935

142. STRATEGIES FOR EQUITABLE PHARMACOGENOMIC-GUIDED WARFARIN DOSING AMONG EUROPEAN AND AFRICAN AMERICAN INDIVIDUALS IN A CLINICAL POPULATION Laura K. Wiley, Jacob P. Vanhouten, David C. Samuels, Melinda C. Aldrich, Dan M. Roden, Josh F. Peterson, Joshua C. Denny Biocomputing 2017 (2016-11-22) https://doi.org/10.1142/9789813207813_0050

143. Epidemiological research labelled as a violation of privacy: the case of Estonia M. Rahu, M. McKee International Journal of Epidemiology (2008-02-26) https://doi.org/10.1093/ije/dyn022

144. Harnessing next-generation informatics for personalizing medicine: a report from AMIA’s 2014 Health Policy Invitational Meeting Laura K Wiley, Peter Tarczy-Hornoch, Joshua C Denny, Robert R Freimuth, Casey L Overby, Nigam Shah, Ross D Martin, Indra Neil Sarkar Journal of the American Medical Informatics Association (2016-02-05) https://doi.org/10.1093/jamia/ocv111

145. DataSHIELD: taking the analysis to the data, not the data to the analysis Amadou Gaye, Yannick Marcon, Julia Isaeva, Philippe LaFlamme, Andrew Turner, Elinor M Jones, Joel Minion, Andrew W Boyd, Christopher J Newby, Marja-Liisa Nuotio, … Paul R Burton International Journal of Epidemiology (2014-09-27) https://doi.org/10.1093/ije/dyu188

146. ViPAR: a software platform for the Virtual Pooling and Analysis of Research Data Kim W Carter, KW Carter, RW Francis, M Bresnahan, M Gissler, TK Grønborg, R Gross, N Gunnes, G Hammond, M Hornig, … Z Yusof International Journal of Epidemiology (2015-10-08) https://doi.org/10.1093/ije/dyv193

147. Reproducibility of computational workflows is automated using continuous analysis Brett K Beaulieu-Jones, Casey S Greene Nature Biotechnology (2017-03-13) https://doi.org/10.1038/nbt.3780

148. Stealing Machine Learning Models via Prediction APIs Florian Tramèr, Fan Zhang, Ari Juels, Michael K. Reiter, Thomas Ristenpart arXiv (2016-09-09) https://arxiv.org/abs/1609.02943v2

149. The Algorithmic Foundations of Differential Privacy Cynthia Dwork, Aaron Roth Foundations and Trends^®^ in Theoretical Computer Science (2013) https://doi.org/10.1561/0400000042

150. Membership Inference Attacks against Machine Learning Models Reza Shokri, Marco Stronati, Congzheng Song, Vitaly Shmatikov arXiv (2016-10-18) https://arxiv.org/abs/1610.05820v2

151. Enabling Privacy-Preserving GWASs in Heterogeneous Human Populations Sean Simmons, Cenk Sahinalp, Bonnie Berger Cell Systems (2016-07) https://doi.org/10.1016/j.cels.2016.04.013

152. Deep Learning with Differential Privacy Martin Abadi, Andy Chu, Ian Goodfellow, H. Brendan McMahan, Ilya Mironov, Kunal Talwar, Li Zhang Proceedings of the 2016 ACM SIGSAC Conference on Computer and Communications Security-CCS’16 (2016) https://doi.org/10.1145/2976749.2978318

153. Generating Multi-label Discrete Electronic Health Records using Generative Adversarial Networks Edward Choi, Siddharth Biswal, Bradley Malin, Jon Duke, Walter F. Stewart, Jimeng Sun arXiv (2017-03-19) https://arxiv.org/abs/1703.06490v1

154. Real-valued (Medical) Time Series Generation with Recurrent Conditional GANs Cristóbal Esteban, Stephanie L. Hyland, Gunnar Rätsch arXiv (2017-06-08) https://arxiv.org/abs/1706.02633v1

155. Privacy-preserving generative deep neural networks support clinical data sharing Brett K. Beaulieu-Jones, Zhiwei Steven Wu, Chris Williams, James Brian Byrd, Casey S. Greene Cold Spring Harbor Laboratory (2017-07-05) https://doi.org/10.1101/159756

156. Communication-Efficient Learning of Deep Networks from Decentralized Data Brendan McMahan, Eider Moore, Daniel Ramage, Seth Hampson, Blaise Aguera y Arcas (2017-04-10) http://proceedings.mlr.press/v54/mcmahan17a.html

157. Practical Secure Aggregation for Privacy Preserving Machine Learning Keith Bonawitz, Vladimir Ivanov, Ben Kreuter, Antonio Marcedone, H. Brendan McMahan, Sarvar Patel, Daniel Ramage, Aaron Segal, Karn Seth (2017) https://eprint.iacr.org/2017/281

158. Semi-supervised Knowledge Transfer for Deep Learning from Private Training Data Nicolas Papernot, Martín Abadi, Úlfar Erlingsson, Ian Goodfellow, Kunal Talwar (2016-11-02) https://openreview.net/forum?id=HkwoSDPgg

159. European Union regulations on algorithmic decision-making and a “right to explanation” Bryce Goodman, Seth Flaxman arXiv (2016-06-28) https://arxiv.org/abs/1606.08813v3

160. Overcoming the Winner’s Curse: Estimating Penetrance Parameters from Case-Control Data Sebastian Zöllner, Jonathan K. Pritchard The American Journal of Human Genetics (2007-04) https://doi.org/10.1086/512821

161. Sex bias in neuroscience and biomedical research Annaliese K. Beery, Irving Zucker Neuroscience & Biobehavioral Reviews (2011-01) https://doi.org/10.1016/j.neubiorev.2010.07.002

162. Generalization and Dilution of Association Results from European GWAS in Populations of Non-European Ancestry: The PAGE Study Christopher S. Carlson, Tara C. Matise, Kari E. North, Christopher A. Haiman, Megan D. Fesinmeyer, Steven Buyske, Fredrick R. Schumacher, Ulrike Peters, Nora Franceschini, Marylyn D. Ritchie, … PLoS Biology (2013-09-17) https://doi.org/10.1371/journal.pbio.1001661

163. New approaches to population stratification in genome-wide association studies Alkes L. Price, Noah A. Zaitlen, David Reich, Nick Patterson Nature Reviews Genetics (2010-06-15) https://doi.org/10.1038/nrg2813

164. Retraction P. Sebastiani, N. Solovieff, A. Puca, S. W. Hartley, E. Melista, S. Andersen, D. A. Dworkis, J. B. Wilk, R. H. Myers, M. H. Steinberg, … T. T. Perls Science (2011-07-21) https://doi.org/10.1126/science.333.6041.404-a

165. Leakage in data mining Shachar Kaufman, Saharon Rosset, Claudia Perlich, Ori Stitelman ACM Transactions on Knowledge Discovery from Data (2012-12-01) https://doi.org/10.1145/2382577.2382579

166. To predict and serve? Kristian Lum, William Isaac Significance (2016-10) https://doi.org/10.1111/j.1740-9713.2016.00960.x

167. Equality of Opportunity in Supervised Learning Moritz Hardt, Eric Price, Nathan Srebro arXiv (2016-10-07) https://arxiv.org/abs/1610.02413v1

168. Fair Algorithms for Infinite and Contextual Bandits Matthew Joseph, Michael Kearns, Jamie Morgenstern, Seth Neel, Aaron Roth arXiv (2016-10-29) https://arxiv.org/abs/1610.09559v4

169. The Framingham Heart Study and the epidemiology of cardiovascular disease: a historical perspective Syed S Mahmood, Daniel Levy, Ramachandran S Vasan, Thomas J Wang The Lancet (2014-03) https://doi.org/10.1016/s0140-6736(13)61752-3

170. Children of the 90s: Coming of age Helen Pearson Nature (2012-04-11) https://doi.org/10.1038/484155a

171. Nonparametric Estimation from Incomplete Observations E. L. Kaplan, Paul Meier Journal of the American Statistical Association (1958-06) https://doi.org/10.2307/2281868

172. Temporal disease trajectories condensed from population-wide registry data covering 6.2 million patients Anders Boeck Jensen, Pope L. Moseley, Tudor I. Oprea, Sabrina Gade Ellesøe, Robert Eriksson, Henriette Schmock, Peter Bjødstrup Jensen, Lars Juhl Jensen, Søren Brunak Nature Communications (2014-06-24) https://doi.org/10.1038/ncomms5022

173. Deepr: A Convolutional Net for Medical Records Phuoc Nguyen, Truyen Tran, Nilmini Wickramasinghe, Svetha Venkatesh arXiv (2016-07-26) https://arxiv.org/abs/1607.07519v1

174. Curiosity Creates Cures: The Value and Impact of Basic Research NIH (2012-05) https://www.nigms.nih.gov/Education/Documents/curiosity.pdf

175. Multi-omics integration accurately predicts cellular state in unexplored conditions for Escherichia coli Minseung Kim, Navneet Rai, Violeta Zorraquino, Ilias Tagkopoulos Nature Communications (2016-10-07) https://doi.org/10.1038/ncomms13090

176. Trans-species learning of cellular signaling systems with bimodal deep belief networks Lujia Chen, Chunhui Cai, Vicky Chen, Xinghua Lu Bioinformatics (2015-05-20) https://doi.org/10.1093/bioinformatics/btv315

177. Learning structure in gene expression data using deep architectures, with an application to gene clustering Aman Gupta, Haohan Wang, Madhavi Ganapathiraju Cold Spring Harbor Laboratory (2015-11-16) https://doi.org/10.1101/031906

178. Learning a hierarchical representation of the yeast transcriptomic machinery using an autoencoder model Lujia Chen, Chunhui Cai, Vicky Chen, Xinghua Lu BMC Bioinformatics (2016-01-11) https://doi.org/10.1186/s12859-015-0852-1

179. ADAGE-Based Integration of Publicly AvailablePseudomonas aeruginosaGene Expression Data with Denoising Autoencoders Illuminates Microbe-Host Interactions Jie Tan, John H. Hammond, Deborah A. Hogan, Casey S. Greene mSystems (2016-01-19) https://doi.org/10.1128/msystems.00025-15

180. Unsupervised extraction of stable expression signatures from public compendia with eADAGE Jie Tan, Georgia Doing, Kimberley A Lewis, Courtney E Price, Kathleen M Chen, Kyle C Cady, Barret Perchuk, Michael T Laub, Deborah A Hogan, Casey S Greene Cold Spring Harbor Laboratory (2016-10-03) https://doi.org/10.1101/078659

181. Gene expression inference with deep learning Yifei Chen, Yi Li, Rajiv Narayan, Aravind Subramanian, Xiaohui Xie Bioinformatics (2016-02-11) https://doi.org/10.1093/bioinformatics/btw074

182. DeepChrome: Deep-learning for predicting gene expression from histone modifications Ritambhara Singh, Jack Lanchantin, Gabriel Robins, Yanjun Qi arXiv (2016-07-07) https://arxiv.org/abs/1607.02078v1

183. Attend and Predict: Understanding Gene Regulation by Selective Attention on Chromatin Ritambhara Singh, Jack Lanchantin, Arshdeep Sekhon, Yanjun Qi arXiv (2017-08-01) https://arxiv.org/abs/1708.00339v3

184. Integrative Data Analysis of Multi-Platform Cancer Data with a Multimodal Deep Learning Approach Muxuan Liang, Zhizhong Li, Ting Chen, Jianyang Zeng IEEE/ACM Transactions on Computational Biology and Bioinformatics (2015-07-01) https://doi.org/10.1109/tcbb.2014.2377729

185. RNA mis-splicing in disease Marina M. Scotti, Maurice S. Swanson Nature Reviews Genetics (2015-11-23) https://doi.org/10.1038/nrg.2015.3

186. RNA splicing is a primary link between genetic variation and disease Y. I. Li, B. van de Geijn, A. Raj, D. A. Knowles, A. A. Petti, D. Golan, Y. Gilad, J. K. Pritchard Science (2016-04-28) https://doi.org/10.1126/science.aad9417

187. Deciphering the splicing code Yoseph Barash, John A. Calarco, Weijun Gao, Qun Pan, Xinchen Wang, Ofer Shai, Benjamin J. Blencowe, Brendan J. Frey Nature (2010-05-06) https://doi.org/10.1038/nature09000

188. Bayesian prediction of tissue-regulated splicing using RNA sequence and cellular context Hui Yuan Xiong, Yoseph Barash, Brendan J. Frey Bioinformatics (2011-07-29) https://doi.org/10.1093/bioinformatics/btr444

189. The human splicing code reveals new insights into the genetic determinants of disease H. Y. Xiong, B. Alipanahi, L. J. Lee, H. Bretschneider, D. Merico, R. K. C. Yuen, Y. Hua, S. Gueroussov, H. S. Najafabadi, T. R. Hughes, … B. J. Frey Science (2014-12-18) https://doi.org/10.1126/science.1254806

190. Integrative Deep Models for Alternative Splicing Anupama Jha, Matthew R Gazzara, Yoseph Barash Cold Spring Harbor Laboratory (2017-01-31) https://doi.org/10.1101/104869

191. Imputation for transcription factor binding predictions based on deep learning Qian Qin, Jianxing Feng PLOS Computational Biology (2017-02-24) https://doi.org/10.1371/journal.pcbi.1005403

192. Learning the Sequence Determinants of Alternative Splicing from Millions of Random Sequences Alexander B. Rosenberg, Rupali P. Patwardhan, Jay Shendure, Georg Seelig Cell (2015-10) https://doi.org/10.1016/j.cell.2015.09.054

193. MECHANISMS IN ENDOCRINOLOGY: Alternative splicing: the new frontier in diabetes research Jonàs Juan-Mateu, Olatz Villate, Décio L Eizirik European Journal of Endocrinology (2015-12-01) https://doi.org/10.1530/eje-15-0916

194. Absence of a simple code: how transcription factors read the genome Matthew Slattery, Tianyin Zhou, Lin Yang, Ana Carolina Dantas Machado, Raluca Gordân, Remo Rohs Trends in Biochemical Sciences (2014-09) https://doi.org/10.1016/j.tibs.2014.07.002

195. An integrated encyclopedia of DNA elements in the human genome Ian Dunham, Anshul Kundaje, Shelley F. Aldred, Patrick J. Collins, Carrie A. Davis, Francis Doyle, Charles B. Epstein, Seth Frietze, Jennifer Harrow, Rajinder Kaul, … Ewan Birney Nature (2012-09-05) https://doi.org/10.1038/nature11247

196. DNA binding sites: representation and discovery G. D. Stormo Bioinformatics (2000-01-01) https://doi.org/10.1093/bioinformatics/16.1.16

197. MEME SUITE: tools for motif discovery and searching T. L. Bailey, M. Boden, F. A. Buske, M. Frith, C. E. Grant, L. Clementi, J. Ren, W. W. Li, W. S. Noble Nucleic Acids Research (2009-05-20) https://doi.org/10.1093/nar/gkp335

198. Evaluation of methods for modeling transcription factor sequence specificity Matthew T Weirauch, Atina Cote, Raquel Norel, Matti Annala, Yue Zhao, Todd R Riley, Julio Saez-Rodriguez, Thomas Cokelaer, Anastasia Vedenko, Shaheynoor Talukder, … Timothy R Hughes Nature Biotechnology (2013-01-27) https://doi.org/10.1038/nbt.2486

199. High Resolution Models of Transcription Factor-DNA Affinities Improve In Vitro and In Vivo Binding Predictions Phaedra Agius, Aaron Arvey, William Chang, William Stafford Noble, Christina Leslie PLoS Computational Biology (2010-09-09) https://doi.org/10.1371/journal.pcbi.1000916

200. Enhanced Regulatory Sequence Prediction Using Gapped k-mer Features Mahmoud Ghandi, Dongwon Lee, Morteza Mohammad-Noori, Michael A. Beer PLoS Computational Biology (2014-07-17) https://doi.org/10.1371/journal.pcbi.1003711

201. Predicting the sequence specificities of DNA-and RNA-binding proteins by deep learning Babak Alipanahi, Andrew Delong, Matthew T Weirauch, Brendan J Frey Nature Biotechnology (2015-07-27) https://doi.org/10.1038/nbt.3300

202. RNA-protein binding motifs mining with a new hybrid deep learning based cross-domain knowledge integration approach Xiaoyong Pan, Hong-Bin Shen BMC Bioinformatics (2017-02-28) https://doi.org/10.1186/s12859-017-1561-8

203. Convolutional neural network architectures for predicting DNA–protein binding Haoyang Zeng, Matthew D. Edwards, Ge Liu, David K. Gifford Bioinformatics (2016-06-15) https://doi.org/10.1093/bioinformatics/btw255

204. Deep Motif Dashboard: Visualizing and Understanding Genomic Sequences Using Deep Neural Networks Jack Lanchantin, Ritambhara Singh, Beilun Wang, Yanjun Qi arXiv (2016-08-12) https://arxiv.org/abs/1608.03644v4

205. Convolutional Kitchen Sinks for Transcription Factor Binding Site Prediction Alyssa Morrow, Vaishaal Shankar, Devin Petersohn, Anthony Joseph, Benjamin Recht, Nir Yosef arXiv (2017-05-31) https://arxiv.org/abs/1706.00125v1

206. Predicting Transcription Factor Binding Sites with Convolutional Kernel Networks Dexiong Chen, Laurent Jacob, Julien Mairal Cold Spring Harbor Laboratory (2017-11-10) https://doi.org/10.1101/217257

207. Reverse-complement parameter sharing improves deep learning models for genomics Avanti Shrikumar, Peyton Greenside, Anshul Kundaje Cold Spring Harbor Laboratory (2017-01-27) https://doi.org/10.1101/103663

208. Separable Fully Connected Layers Improve Deep Learning Models For Genomics Amr Mohamed Alexandari, Avanti Shrikumar, Anshul Kundaje Cold Spring Harbor Laboratory (2017-06-05) https://doi.org/10.1101/146431

209. Predicting effects of noncoding variants with deep learning–based sequence model Jian Zhou, Olga G Troyanskaya Nature Methods (2015-08-24) https://doi.org/10.1038/nmeth.3547

210. DanQ: a hybrid convolutional and recurrent deep neural network for quantifying the function of DNA sequences Daniel Quang, Xiaohui Xie Nucleic Acids Research (2016-04-15) https://doi.org/10.1093/nar/gkw226

211. Sequence and chromatin determinants of cell-type-specific transcription factor binding A. Arvey, P. Agius, W. S. Noble, C. Leslie Genome Research (2012-09-01) https://doi.org/10.1101/gr.127712.111

212. Analysis of computational footprinting methods for DNase sequencing experiments Eduardo G Gusmao, Manuel Allhoff, Martin Zenke, Ivan G Costa Nature Methods (2016-02-22) https://doi.org/10.1038/nmeth.3772

213. ENCODE-DREAM in vivo Transcription Factor Binding Site Prediction Challenge(2017) https://www.synapse.org/#!Synapse:syn6131484/wiki/402026

214. FactorNet: a deep learning framework for predicting cell type specific transcription factor binding from nucleotide-resolution sequential data Daniel Quang, Xiaohui Xie Cold Spring Harbor Laboratory (2017-06-18) https://doi.org/10.1101/151274

215. Learning from mistakes: Accurate prediction of cell type-specific transcription factor binding Jens Keilwagen, Stefan Posch, Jan Grau Cold Spring Harbor Laboratory (2017-12-06) https://doi.org/10.1101/230011

216. Transfer String Kernel for Cross-Context DNA-Protein Binding Prediction Ritambhara Singh, Jack Lanchantin, Gabriel Robins, Yanjun Qi IEEE/ACM Transactions on Computational Biology and Bioinformatics (2016) https://doi.org/10.1109/tcbb.2016.2609918

217. Learning Transferable Features with Deep Adaptation Networks Mingsheng Long, Yue Cao, Jianmin Wang, Michael I. Jordan arXiv (2015-02-10) https://arxiv.org/abs/1502.02791v2

218. Domain-Adversarial Training of Neural Networks Yaroslav Ganin, Evgeniya Ustinova, Hana Ajakan, Pascal Germain, Hugo Larochelle, François Laviolette, Mario Marchand, Victor Lempitsky arXiv (2015-05-28) https://arxiv.org/abs/1505.07818v4

219. Learning Important Features Through Propagating Activation Differences Avanti Shrikumar, Peyton Greenside, Anshul Kundaje arXiv (2017-04-10) https://arxiv.org/abs/1704.02685v1

220. The state of the art of mammalian promoter recognition T. Werner Briefings in Bioinformatics (2003-01-01) https://doi.org/10.1093/bib/4.1.22

221. Detection of RNA polymerase II promoters and polyadenylation sites in human DNA sequence Sherri Matis, Ying Xu, Manesh Shah, Xiaojun Guan, J.Ralph Einstein, Richard Mural, Edward Uberbacher Computers & Chemistry (1996-03) https://doi.org/10.1016/s0097-8485(96)80015-5

222. Recognition of prokaryotic and eukaryotic promoters using convolutional deep learning neural networks Ramzan Kh. Umarov, Victor V. Solovyev PLOS ONE (2017-02-03) https://doi.org/10.1371/journal.pone.0171410

223. Cap analysis gene expression for high-throughput analysis of transcriptional starting point and identification of promoter usage T. Shiraki, S. Kondo, S. Katayama, K. Waki, T. Kasukawa, H. Kawaji, R. Kodzius, A. Watahiki, M. Nakamura, T. Arakawa, … Y. Hayashizaki Proceedings of the National Academy of Sciences (2003-12-08) https://doi.org/10.1073/pnas.2136655100

224. Genome-wide characterization of transcriptional start sites in humans by integrative transcriptome analysis R. Yamashita, N. P. Sathira, A. Kanai, K. Tanimoto, T. Arauchi, Y. Tanaka, S.-i. Hashimoto, S. Sugano, K. Nakai, Y. Suzuki Genome Research (2011-03-03) https://doi.org/10.1101/gr.110254.110

225. Enhancers: five essential questions Len A. Pennacchio, Wendy Bickmore, Ann Dean, Marcelo A. Nobrega, Gill Bejerano Nature Reviews Genetics (2013-03-18) https://doi.org/10.1038/nrg3458

226. A unified architecture of transcriptional regulatory elements Robin Andersson, Albin Sandelin, Charles G. Danko Trends in Genetics (2015-08) https://doi.org/10.1016/j.tig.2015.05.007

227. Basset: learning the regulatory code of the accessible genome with deep convolutional neural networks David R. Kelley, Jasper Snoek, John L. Rinn Genome Research (2016-05-03) https://doi.org/10.1101/gr.200535.115

228. DeepEnhancer: Predicting enhancers by convolutional neural networks Xu Min, Ning Chen, Ting Chen, Rui Jiang 2016 IEEE International Conference on Bioinformatics and Biomedicine (BIBM) (2016-12) https://doi.org/10.1109/bibm.2016.7822593

229. Genome-Wide Prediction of cis-Regulatory Regions Using Supervised Deep Learning Methods Yifeng Li, Wenqiang Shi, Wyeth W Wasserman Cold Spring Harbor Laboratory (2016-02-28) https://doi.org/10.1101/041616

230. Predicting Enhancer-Promoter Interaction from Genomic Sequence with Deep Neural Networks Shashank Singh, Yang Yang, Barnabas Poczos, Jian Ma Cold Spring Harbor Laboratory (2016-11-02) https://doi.org/10.1101/085241

231. A network-biology perspective of microRNA function and dysfunction in cancer Cameron P. Bracken, Hamish S. Scott, Gregory J. Goodall Nature Reviews Genetics (2016-10-31) https://doi.org/10.1038/nrg.2016.134

232. Evolution of microRNA diversity and regulation in animals Eugene Berezikov Nature Reviews Genetics (2011-11-18) https://doi.org/10.1038/nrg3079

233. Predicting effective microRNA target sites in mammalian mRNAs Vikram Agarwal, George W Bell, Jin-Wu Nam, David P Bartel eLife (2015-08-12) https://doi.org/10.7554/elife.05005

234. deepTarget: End-to-end Learning Framework for microRNA Target Prediction using Deep Recurrent Neural Networks Byunghan Lee, Junghwan Baek, Seunghyun Park, Sungroh Yoon arXiv (2016-03-30) https://arxiv.org/abs/1603.09123v2

235. deepMiRGene: Deep Neural Network based Precursor microRNA Prediction Seunghyun Park, Seonwoo Min, Hyunsoo Choi, Sungroh Yoon arXiv (2016-04-29) https://arxiv.org/abs/1605.00017v1

236. AUC-Maximized Deep Convolutional Neural Fields for Protein Sequence Labeling Sheng Wang, Siqi Sun, Jinbo Xu Machine Learning and Knowledge Discovery in Databases (2016) https://doi.org/10.1007/978-3-319-46227-1_1

237. MetaPSICOV: combining coevolution methods for accurate prediction of contacts and long range hydrogen bonding in proteins David T. Jones, Tanya Singh, Tomasz Kosciolek, Stuart Tetchner Bioinformatics (2014-11-26) https://doi.org/10.1093/bioinformatics/btu791

238. Identification of direct residue contacts in protein-protein interaction by message passing M. Weigt, R. A. White, H. Szurmant, J. A. Hoch, T. Hwa Proceedings of the National Academy of Sciences (2008-12-30) https://doi.org/10.1073/pnas.0805923106

239. Protein 3D Structure Computed from Evolutionary Sequence Variation Debora S. Marks, Lucy J. Colwell, Robert Sheridan, Thomas A. Hopf, Andrea Pagnani, Riccardo Zecchina, Chris Sander PLoS ONE (2011-12-07) https://doi.org/10.1371/journal.pone.0028766

240. A Unified Multitask Architecture for Predicting Local Protein Properties Yanjun Qi, Merja Oja, Jason Weston, William Stafford Noble PLoS ONE (2012-03-26) https://doi.org/10.1371/journal.pone.0032235

241. Improving prediction of secondary structure, local backbone angles and solvent accessible surface area of proteins by iterative deep learning Rhys Heffernan, Kuldip Paliwal, James Lyons, Abdollah Dehzangi, Alok Sharma, Jihua Wang, Abdul Sattar, Yuedong Yang, Yaoqi Zhou Scientific Reports (2015-06-22) https://doi.org/10.1038/srep11476

242. Protein secondary structure prediction based on position-specific scoring matrices 1 1Edited by G. Von Heijne David T Jones Journal of Molecular Biology (1999-09) https://doi.org/10.1006/jmbi.1999.3091

243. Deep Supervised and Convolutional Generative Stochastic Network for Protein Secondary Structure Prediction Jian Zhou, Olga G. Troyanskaya arXiv (2014-03-06) https://arxiv.org/abs/1403.1347v1

244. Protein contact prediction by integrating joint evolutionary coupling analysis and supervised learning Jianzhu Ma, Sheng Wang, Zhiyong Wang, Jinbo Xu Bioinformatics (2015-08-14) https://doi.org/10.1093/bioinformatics/btv472

245. Deep architectures for protein contact map prediction Pietro Di Lena, Ken Nagata, Pierre Baldi Bioinformatics (2012-07-30) https://doi.org/10.1093/bioinformatics/bts475

246. Predicting protein residue–residue contacts using deep networks and boosting Jesse Eickholt, Jianlin Cheng Bioinformatics (2012-10-09) https://doi.org/10.1093/bioinformatics/bts598

247. Improved Contact Predictions Using the Recognition of Protein Like Contact Patterns Marcin J. Skwark, Daniele Raimondi, Mirco Michel, Arne Elofsson PLoS Computational Biology (2014-11-06) https://doi.org/10.1371/journal.pcbi.1003889

248. RR Results - CASP12(2017) http://www.predictioncenter.org/casp12/rrc_avrg_results.cgi

249. CAMEO - Continuous Automated Model Evaluation(2017) http://www.cameo3d.org/

250. Predicting membrane protein contacts from non-membrane proteins by deep transfer learning Zhen Li, Sheng Wang, Yizhou Yu, Jinbo Xu arXiv (2017-04-24) https://arxiv.org/abs/1704.07207v1

251. Single-Particle Cryo-EM at Crystallographic Resolution Yifan Cheng Cell (2015-04) https://doi.org/10.1016/j.cell.2015.03.049

252. A Primer to Single-Particle Cryo-Electron Microscopy Yifan Cheng, Nikolaus Grigorieff, Pawel A. Penczek, Thomas Walz Cell (2015-04) https://doi.org/10.1016/j.cell.2015.03.050

253. SwarmPS: Rapid, semi-automated single particle selection software David Woolford, Geoffery Ericksson, Rosalba Rothnagel, David Muller, Michael J. Landsberg, Radosav S. Pantelic, Alasdair McDowall, Bernard Pailthorpe, Paul R. Young, Ben Hankamer, Jasmine Banks Journal of Structural Biology (2007-01) https://doi.org/10.1016/j.jsb.2006.04.006

254. Semi-automated selection of cryo-EM particles in RELION-1.3 Sjors H.W. Scheres Journal of Structural Biology (2015-02) https://doi.org/10.1016/j.jsb.2014.11.010

255. DeepPicker: A deep learning approach for fully automated particle picking in cryo-EM Feng Wang, Huichao Gong, Gaochao Liu, Meijing Li, Chuangye Yan, Tian Xia, Xueming Li, Jianyang Zeng Journal of Structural Biology (2016-09) https://doi.org/10.1016/j.jsb.2016.07.006

256. A deep convolutional neural network approach to single-particle recognition in cryo-electron microscopy Yanan Zhu, Qi Ouyang, Youdong Mao BMC Bioinformatics (2017-07-21) https://doi.org/10.1186/s12859-017-1757-y

257. Massively parallel unsupervised single-particle cryo-EM data clustering via statistical manifold learning Jiayi Wu, Yong-Bei Ma, Charles Congdon, Bevin Brett, Shuobing Chen, Yaofang Xu, Qi Ouyang, Youdong Mao PLOS ONE (2017-08-07) https://doi.org/10.1371/journal.pone.0182130

258. Protein–Protein Interactions Essentials: Key Concepts to Building and Analyzing Interactome Networks Javier De Las Rivas, Celia Fontanillo PLoS Computational Biology (2010-06-24) https://doi.org/10.1371/journal.pcbi.1000807

259. Extracting interactions between proteins from the literature Deyu Zhou, Yulan He Journal of Biomedical Informatics (2008-04) https://doi.org/10.1016/j.jbi.2007.11.008

260. Deep learning for extracting protein-protein interactions from biomedical literature Yifan Peng, Zhiyong Lu arXiv (2017-06-05) href="https://arxiv.org/abs/1706.01556v2

261. DeepPPI: Boosting Prediction of Protein–Protein Interactions with Deep Neural Networks Xiuquan Du, Shiwei Sun, Changlin Hu, Yu Yao, Yuanting Yan, Yanping Zhang Journal of Chemical Information and Modeling (2017-05-26) https://doi.org/10.1021/acs.jcim.7b00028

262. Sequence-based prediction of protein protein interaction using a deep-learning algorithm Tanlin Sun, Bo Zhou, Luhua Lai, Jianfeng Pei BMC Bioinformatics (2017-05-25) https://doi.org/10.1186/s12859-017-1700-2

263. Predicting protein–protein interactions from protein sequences by a stacked sparse autoencoder deep neural network Yan-Bin Wang, Zhu-Hong You, Xiao Li, Tong-Hai Jiang, Xing Chen, Xi Zhou, Lei Wang Molecular BioSystems (2017) https://doi.org/10.1039/c7mb00188f

264. Prediction of residue-residue contact matrix for protein-protein interaction with Fisher score features and deep learning Tianchuan Du, Li Liao, Cathy H. Wu, Bilin Sun Methods (2016-11) https://doi.org/10.1016/j.ymeth.2016.06.001

265. Reliable prediction of T-cell epitopes using neural networks with novel sequence representations Morten Nielsen, Claus Lundegaard, Peder Worning, Sanne Lise Lauemøller, Kasper Lamberth, Søren Buus, Søren Brunak, Ole Lund Protein Science (2003-05) https://doi.org/10.1110/ps.0239403

266. Gapped sequence alignment using artificial neural networks: application to the MHC class I system Massimo Andreatta, Morten Nielsen Bioinformatics (2015-10-29) https://doi.org/10.1093/bioinformatics/btv639

267. NetMHCpan, a method for MHC class I binding prediction beyond humans Ilka Hoof, Bjoern Peters, John Sidney, Lasse Eggers Pedersen, Alessandro Sette, Ole Lund, Søren Buus, Morten Nielsen Immunogenetics (2008-11-12) https://doi.org/10.1007/s00251-008-0341-z

268. NetMHCpan-3.0; improved prediction of binding to MHC class I molecules integrating information from multiple receptor and peptide length datasets Morten Nielsen, Massimo Andreatta Genome Medicine (2016-03-30) https://doi.org/10.1186/s13073-016-0288-x

269. MHCflurry: open-source class I MHC binding affinity prediction Timothy O’Donnell, Alex Rubinsteyn, Maria Bonsack, Angelika Riemer, Jeffrey Hammerbacher Cold Spring Harbor Laboratory (2017-08-09) https://doi.org/10.1101/174243

270. Predicting Peptide-MHC Binding Affinities With Imputed Training Data Alex Rubinsteyn, Timothy O’Donnell, Nandita Damaraju, Jeffrey Hammerbacher Cold Spring Harbor Laboratory (2016-05-22) https://doi.org/10.1101/054775

271. High-order neural networks and kernel methods for peptide-MHC binding prediction Pavel P. Kuksa, Martin Renqiang Min, Rishabh Dugar, Mark Gerstein Bioinformatics (2015-07-23) https://doi.org/10.1093/bioinformatics/btv371

272. Evaluation of machine learning methods to predict peptide binding to MHC Class I proteins Rohit Bhattacharya, Ashok Sivakumar, Collin Tokheim, Violeta Beleva Guthrie, Valsamo Anagnostou, Victor E. Velculescu, Rachel Karchin Cold Spring Harbor Laboratory (2017-06-23) https://doi.org/10.1101/154757

273. HLA class I binding prediction via convolutional neural networks Yeeleng S. Vang, Xiaohui Xie Bioinformatics (2017-04-21) https://doi.org/10.1093/bioinformatics/btx264

274. Network-based prediction of protein function Roded Sharan, Igor Ulitsky, Ron Shamir Molecular Systems Biology (2007-03-13) https://doi.org/10.1038/msb4100129

275. Learning the Structural Vocabulary of a Network Saket Navlakha Neural Computation (2017-02) https://doi.org/10.1162/neco_a_00924

276. deepNF: Deep network fusion for protein function prediction Vladimir Gligorijević, Meet Barot, Richard Bonneau Cold Spring Harbor Laboratory (2017-11-22) https://doi.org/10.1101/223339

277. Inductive Representation Learning on Large Graphs William L. Hamilton, Rex Ying, Jure Leskovec arXiv (2017-06-07) https://arxiv.org/abs/1706.02216v2

278. Stochastic Training of Graph Convolutional Networks Jianfei Chen, Jun Zhu arXiv (2017-10-29) https://arxiv.org/abs/1710.10568v1

279. Deep Learning Automates the Quantitative Analysis of Individual Cells in Live-Cell Imaging Experiments David A. Van Valen, Takamasa Kudo, Keara M. Lane, Derek N. Macklin, Nicolas T. Quach, Mialy M. DeFelice, Inbal Maayan, Yu Tanouchi, Euan A. Ashley, Markus W. Covert PLOS Computational Biology (2016-11-04) https://doi.org/10.1371/journal.pcbi.1005177

280. U-Net: Convolutional Networks for Biomedical Image Segmentation Olaf Ronneberger, Philipp Fischer, Thomas Brox Lecture Notes in Computer Science (2015) https://doi.org/10.1007/978-3-319-24574-4_28

281. Prospective identification of hematopoietic lineage choice by deep learning Felix Buggenthin, Florian Buettner, Philipp S Hoppe, Max Endele, Manuel Kroiss, Michael Strasser, Michael Schwarzfischer, Dirk Loeffler, Konstantinos D Kokkaliaris, Oliver Hilsenbeck, … Carsten Marr Nature Methods (2017-02-20) https://doi.org/10.1038/nmeth.4182

282. Reconstructing cell cycle and disease progression using deep learning Philipp Eulenberg, Niklas Koehler, Thomas Blasi, Andrew Filby, Anne E. Carpenter, Paul Rees, Fabian J. Theis, F. Alexander Wolf Cold Spring Harbor Laboratory (2016-10-17) https://doi.org/10.1101/081364

283. Automating Morphological Profiling with Generic Deep Convolutional Networks Nick Pawlowski, Juan C Caicedo, Shantanu Singh, Anne E Carpenter, Amos Storkey Cold Spring Harbor Laboratory (2016-11-02) https://doi.org/10.1101/085118

284. Generative Modeling with Conditional Autoencoders: Building an Integrated Cell Gregory R. Johnson, Rory M. Donovan-Maiye, Mary M. Maleckar arXiv (2017-04-28) https://arxiv.org/abs/1705.00092v1

285. Applications in image-based profiling of perturbations Juan C Caicedo, Shantanu Singh, Anne E Carpenter Current Opinion in Biotechnology (2016-06) https://doi.org/10.1016/j.copbio.2016.04.003

286. Large-scale image-based screening and profiling of cellular phenotypes Nicola Bougen-Zhukov, Sheng Yang Loh, Hwee Kuan Lee, Lit-Hsin Loo Cytometry Part A (2016-07-19) https://doi.org/10.1002/cyto.a.22909

287. Machine learning and computer vision approaches for phenotypic profiling Ben T. Grys, Dara S. Lo, Nil Sahin, Oren Z. Kraus, Quaid Morris, Charles Boone, Brenda J. Andrews The Journal of Cell Biology (2016-12-09) https://doi.org/10.1083/jcb.201610026

288. Single-cell genome sequencing: current state of the science Charles Gawad, Winston Koh, Stephen R. Quake Nature Reviews Genetics (2016-01-25) https://doi.org/10.1038/nrg.2015.16

289. Somatic mutation in single human neurons tracks developmental and transcriptional history M. A. Lodato, M. B. Woodworth, S. Lee, G. D. Evrony, B. K. Mehta, A. Karger, S. Lee, T. W. Chittenden, A. M. D’Gama, X. Cai, … C. A. Walsh Science (2015-10-01) https://doi.org/10.1126/science.aab1785

290. Single-cell transcriptome sequencing: recent advances and remaining challenges Serena Liu, Cole Trapnell F1000Research (2016-02-17) https://doi.org/10.12688/f1000research.7223.1

291. Single-Cell and Single-Molecule Analysis of Gene Expression Regulation Maria Vera, Jeetayu Biswas, Adrien Senecal, Robert H. Singer, Hye Yoon Park Annual Review of Genetics (2016-11-23) https://doi.org/10.1146/annurev-genet-120215-034854

292. Joint Profiling Of Chromatin Accessibility, DNA Methylation And Transcription In Single Cells Stephen J. Clark, Ricard Argelaguet, Chantriolnt-Andreas Kapourani, Thomas M. Stubbs, Heather J. Lee, Felix Krueger, Guido Sanguinetti, Gavin Kelsey, John C. Marioni, Oliver Stegle, Wolf Reik Cold Spring Harbor Laboratory (2017-05-17) https://doi.org/10.1101/138685

293. DeepCpG: accurate prediction of single-cell DNA methylation states using deep learning Christof Angermueller, Heather J. Lee, Wolf Reik, Oliver Stegle Genome Biology (2017-04-11) https://doi.org/10.1186/s13059-017-1189-z

294. Denoising genome-wide histone ChIP-seq with convolutional neural networks Pang Wei Koh, Emma Pierson, Anshul Kundaje Cold Spring Harbor Laboratory (2016-05-07) https://doi.org/10.1101/052118

295. Removal of batch effects using distribution-matching residual networks Uri Shaham, Kelly P. Stanton, Jun Zhao, Huamin Li, Khadir Raddassi, Ruth Montgomery, Yuval Kluger Bioinformatics (2017-04-13) https://doi.org/10.1093/bioinformatics/btx196

296. Single-Cell Genomics Unveils Critical Regulators of Th17 Cell Pathogenicity Jellert T. Gaublomme, Nir Yosef, Youjin Lee, Rona S. Gertner, Li V. Yang, Chuan Wu, Pier Paolo Pandolfi, Tak Mak, Rahul Satija, Alex K. Shalek, … Aviv Regev Cell (2015-12) https://doi.org/10.1016/j.cell.2015.11.009

297. Sensitive detection of rare disease-associated cell subsets via representation learning. Eirini Arvaniti, Manfred Claassen Cold Spring Harbor Laboratory (2016-03-31) https://doi.org/10.1101/046508

298. Interpretable dimensionality reduction of single cell transcriptome data with deep generative models Jiarui Ding, Anne E. Condon, Sohrab P. Shah Cold Spring Harbor Laboratory (2017-09-01) https://doi.org/10.1101/178624

299. A deep generative model for gene expression profiles from single-cell RNA sequencing Romain Lopez, Jeffrey Regier, Michael Cole, Michael Jordan, Nir Yosef arXiv (2017-09-07) https://arxiv.org/abs/1709.02082v3

300. Visualizing Data using t-SNE Laurens van der Maaten, Geoffrey Hinton Journal of Machine Learning Research (2008) http://www.jmlr.org/papers/v9/vandermaaten08a.html

301. Using neural networks for reducing the dimensions of single-cell RNA-Seq data Chieh Lin, Siddhartha Jain, Hannah Kim, Ziv Bar-Joseph Nucleic Acids Research (2017-07-31) https://doi.org/10.1093/nar/gkx681

302. Science Forum: The Human Cell Atlas Aviv Regev, Sarah A Teichmann, Eric S Lander, Ido Amit, Christophe Benoist, Ewan Birney, Bernd Bodenmiller, Peter J Campbell, Piero Carninci, Menna Clatworthy, … eLife (2017-12-05) https://doi.org/10.7554/elife.27041

303. Reversed graph embedding resolves complex single-cell developmental trajectories Xiaojie Qiu, Qi Mao, Ying Tang, Li Wang, Raghav Chawla, Hannah Pliner, Cole Trapnell Cold Spring Harbor Laboratory (2017-02-21) https://doi.org/10.1101/110668

304. Mastering the game of Go with deep neural networks and tree search David Silver, Aja Huang, Chris J. Maddison, Arthur Guez, Laurent Sifre, George van den Driessche, Julian Schrittwieser, Ioannis Antonoglou, Veda Panneershelvam, Marc Lanctot, … Demis Hassabis Nature (2016-01-27) https://doi.org/10.1038/nature16961

305. Compositional biases of bacterial genomes and evolutionary implications. S Karlin, J Mrázek, AM Campbell Journal of Bacteriology (1997-06) https://doi.org/10.1128/jb.179.12.3899-3913.1997

306. Accurate phylogenetic classification of variable-length DNA fragments Alice Carolyn McHardy, Héctor García Martín, Aristotelis Tsirigos, Philip Hugenholtz, Isidore Rigoutsos Nature Methods (2006-12-10) https://doi.org/10.1038/nmeth976

307. NBC: the Naive Bayes Classification tool webserver for taxonomic classification of metagenomic reads G. L. Rosen, E. R. Reichenberger, A. M. Rosenfeld Bioinformatics (2010-11-08) https://doi.org/10.1093/bioinformatics/btq619

308. Informatics for Unveiling Hidden Genome Signatures T. Abe Genome Research (2003-04-01) https://doi.org/10.1101/gr.634603

309. Metagenomic microbial community profiling using unique clade-specific marker genes Nicola Segata, Levi Waldron, Annalisa Ballarini, Vagheesh Narasimhan, Olivier Jousson, Curtis Huttenhower Nature Methods (2012-06-10) https://doi.org/10.1038/nmeth.2066

310. WGSQuikr: Fast Whole-Genome Shotgun Metagenomic Classification David Koslicki, Simon Foucart, Gail Rosen PLoS ONE (2014-03-13) https://doi.org/10.1371/journal.pone.0091784

311. Scalable metagenomic taxonomy classification using a reference genome database Sasha K. Ames, David A. Hysom, Shea N. Gardner, G. Scott Lloyd, Maya B. Gokhale, Jonathan E. Allen Bioinformatics (2013-07-04) https://doi.org/10.1093/bioinformatics/btt389

312. Large-scale machine learning for metagenomics sequence classification Kévin Vervier, Pierre Mahé, Maud Tournoud, Jean-Baptiste Veyrieras, Jean-Philippe Vert Bioinformatics (2015-11-20) https://doi.org/10.1093/bioinformatics/btv683

313. Combining gene prediction methods to improve metagenomic gene annotation Non G Yok, Gail L Rosen BMC Bioinformatics (2011) https://doi.org/10.1186/1471-2105-12-20

314. Machine learning for metagenomics: methods and tools Hayssam Soueidan, Macha Nikolski Metagenomics (2017-01-01) https://doi.org/10.1515/metgen-2016-0001

315. Utilizing Machine Learning Approaches to Understand the Interrelationship of Diet, the Human Gastrointestinal Microbiome, and Health Heather Guetterman, Loretta Auvil, Nate Russell, Michael Welge, Matt Berry, Lisa Gatzke, Colleen Bushell, Hannah Holscher The FASEB Journal (2016-04-01) http://www.fasebj.org/content/30/1_Supplement/406.3

316. Supervised classification of human microbiota Dan Knights, Elizabeth K. Costello, Rob Knight FEMS Microbiology Reviews (2011-03) https://doi.org/10.1111/j.1574-6976.2010.00251.x

317. A comprehensive evaluation of multicategory classification methods for microbiomic data Alexander Statnikov, Mikael Henaff, Varun Narendra, Kranti Konganti, Zhiguo Li, Liying Yang, Zhiheng Pei, Martin J Blaser, Constantin F Aliferis, Alexander V Alekseyenko Microbiome (2013) https://doi.org/10.1186/2049-2618-1-11

318. Machine Learning Meta-analysis of Large Metagenomic Datasets: Tools and Biological Insights Edoardo Pasolli, Duy Tin Truong, Faizan Malik, Levi Waldron, Nicola Segata PLOS Computational Biology (2016-07-11) https://doi.org/10.1371/journal.pcbi.1004977

319. DectICO: an alignment-free supervised metagenomic classification method based on feature extraction and dynamic selection Xiao Ding, Fudong Cheng, Changchang Cao, Xiao Sun BMC Bioinformatics (2015-10-07) https://doi.org/10.1186/s12859-015-0753-3

320. Correction: Class Prediction and Feature Selection with Linear Optimization for Metagenomic Count Data Zhenqiu Liu, Dechang Chen, Li Sheng, Amy Y. Liu PLoS ONE (2014-05-12) https://doi.org/10.1371/journal.pone.0097958

321. Fizzy: feature subset selection for metagenomics Gregory Ditzler, J. Calvin Morrison, Yemin Lan, Gail L. Rosen BMC Bioinformatics (2015-11-04) https://doi.org/10.1186/s12859-015-0793-8

322. A Bootstrap Based Neyman-Pearson Test for Identifying Variable Importance Gregory Ditzler, Robi Polikar, Gail Rosen IEEE Transactions on Neural Networks and Learning Systems (2015-04) https://doi.org/10.1109/tnnls.2014.2320415

323. Orphelia: predicting genes in metagenomic sequencing reads Katharina J. Hoff, Thomas Lingner, Peter Meinicke, Maike Tech Nucleic Acids Research (2009-05-08) https://doi.org/10.1093/nar/gkp327

324. FragGeneScan: predicting genes in short and error-prone reads Mina Rho, Haixu Tang, Yuzhen Ye Nucleic Acids Research (2010-08-28) https://doi.org/10.1093/nar/gkq747

325. Continuous Distributed Representation of Biological Sequences for Deep Proteomics and Genomics Ehsaneddin Asgari, Mohammad R. K. Mofrad PLOS ONE (2015-11-10) https://doi.org/10.1371/journal.pone.0141287

326. Fast model-based protein homology detection without alignment S. Hochreiter, M. Heusel, K. Obermayer Bioinformatics (2007-05-08) https://doi.org/10.1093/bioinformatics/btm247

327. Convolutional LSTM Networks for Subcellular Localization of Proteins Søren Kaae Sønderby, Casper Kaae Sønderby, Henrik Nielsen, Ole Winther Algorithms for Computational Biology (2015) https://doi.org/10.1007/978-3-319-21233-3_6

328. Neural network-based taxonomic clustering for metagenomics Steven D. Essinger, Robi Polikar, Gail L. Rosen The 2010 International Joint Conference on Neural Networks (IJCNN) (2010-07) href="https://doi.org/10.1109/ijcnn.2010.5596644

329. Clustering metagenomic sequences with interpolated Markov models David R Kelley, Steven L Salzberg BMC Bioinformatics (2010) https://doi.org/10.1186/1471-2105-11-544

330. METAGENOMIC TAXONOMIC CLASSIFICATION USING EXTREME LEARNING MACHINES Zeehasham Rasheed, Huzefa Rangwala Journal of Bioinformatics and Computational Biology (2012-10) https://doi.org/10.1142/s0219720012500151

331. Globoko ucenje na genomskih in filogenetskih podatkih Nina Mrzelj Univerza v Ljubljani, Fakulteta za racunalništvo in informatiko (2016) https://repozitorij.uni-lj.si/IzpisGradiva.php?id=85515

332. Influence of microbiome species in hard-to-heal wounds on disease severity and treatment duration Dagmar Chudobova, Kristyna Cihalova, Roman Guran, Simona Dostalova, Kristyna Smerkova, Radek Vesely, Jaromir Gumulec, Michal Masarik, Zbynek Heger, Vojtech Adam, Rene Kizek The Brazilian Journal of Infectious Diseases (2015-11) https://doi.org/10.1016/j.bjid.2015.08.013

333. Multi-Layer and Recursive Neural Networks for Metagenomic Classification Gregory Ditzler, Robi Polikar, Gail Rosen IEEE Transactions on NanoBioscience (2015-09) https://doi.org/10.1109/tnb.2015.2461219

334. TensorFlow vs. scikit-learn: The Microbiome Challenge Ali A. Faruqi Ali A. Faruqi (2016-07-09) http://alifar76.github.io/sklearn-metrics/

335. Advances in Optimizing Recurrent Networks Yoshua Bengio, Nicolas Boulanger-Lewandowski, Razvan Pascanu arXiv (2012-12-04) https://arxiv.org/abs/1212.0901v2

336. DeepNano: Deep recurrent neural networks for base calling in MinION nanopore reads Vladimír Boža, Broňa Brejová, Tomáš Vinař PLOS ONE (2017-06-05) https://doi.org/10.1371/journal.pone.0178751

337. Sequence to Sequence Learning with Neural Networks Ilya Sutskever, Oriol Vinyals, Quoc V. Le arXiv (2014-09-10) https://arxiv.org/abs/1409.3215v3

338. Creating a universal SNP and small indel variant caller with deep neural networks Ryan Poplin, Dan Newburger, Jojo Dijamco, Nam Nguyen, Dion Loy, Sam S. Gross, Cory Y. McLean, Mark A. DePristo Cold Spring Harbor Laboratory (2016-12-14) https://doi.org/10.1101/092890

339. A framework for variation discovery and genotyping using next-generation DNA sequencing data Mark A DePristo, Eric Banks, Ryan Poplin, Kiran V Garimella, Jared R Maguire, Christopher Hartl, Anthony A Philippakis, Guillermo del Angel, Manuel A Rivas, Matt Hanna, … Mark J Daly Nature Genetics (2011-04-10) https://doi.org/10.1038/ng.806

340. Training Genotype Callers with Neural Networks Rémi Torracinta, Fabien Campagne Cold Spring Harbor Laboratory (2016-12-30) https://doi.org/10.1101/097469

341. Xception: Deep Learning with Depthwise Separable Convolutions François Chollet arXiv (2016-10-07) https://arxiv.org/abs/1610.02357v3

342. Adaptive Somatic Mutations Calls with Deep Learning and Semi-Simulated Data Remi Torracinta, Laurent Mesnard, Susan Levine, Rita Shaknovich, Maureen Hanson, Fabien Campagne Cold Spring Harbor Laboratory (2016-10-04) https://doi.org/10.1101/079087

343. Toward an Integration of Deep Learning and Neuroscience Adam H. Marblestone, Greg Wayne, Konrad P. Kording Frontiers in Computational Neuroscience (2016-09-14) https://doi.org/10.3389/fncom.2016.00094

344. Deep Neural Networks In Computational Neuroscience Tim Christian Kietzmann, Patrick McClure, Nikolaus Kriegeskorte Cold Spring Harbor Laboratory (2017-05-04) https://doi.org/10.1101/133504

345. Neuroscience-Inspired Artificial Intelligence Demis Hassabis, Dharshan Kumaran, Christopher Summerfield, Matthew Botvinick Neuron (2017-07) https://doi.org/10.1016/j.neuron.2017.06.011

346. Using goal-driven deep learning models to understand sensory cortex Daniel LK Yamins, James J DiCarlo Nature Neuroscience (2016-02-23) https://doi.org/10.1038/nn.4244

347. Inferring single-trial neural population dynamics using sequential auto-encoders Chethan Pandarinath, Daniel J. O’Shea, Jasmine Collins, Rafal Jozefowicz, Sergey D. Stavisky, Jonathan C. Kao, Eric M. Trautmann, Matthew T. Kaufman, Stephen I. Ryu, Leigh R. Hochberg, … David Sussillo Cold Spring Harbor Laboratory (2017-06-20) https://doi.org/10.1101/152884

348. Machines that learn to segment images: a crucial technology for connectomics Viren Jain, H Sebastian Seung, Srinivas C Turaga Current Opinion in Neurobiology (2010-10) https://doi.org/10.1016/j.conb.2010.07.004

349. Model-based Bayesian inference of neural activity and connectivity from all-optical interrogation of a neural circuit Laurence Aitchison, Lloyd Russell, Adam M. Packer, Jinyao Yan, Philippe Castonguay, Michael Hausser, Srinivas C. Turaga (2017) http://papers.nips.cc/paper/6940-model-based-bayesian-inference-of-neural-activity-and-connectivity-from-all-optical-interrogation-of-a-neural-circuit

350. The Path to Personalized Medicine Margaret A. Hamburg, Francis S. Collins New England Journal of Medicine (2010-07-22) https://doi.org/10.1056/nejmp1006304

351. Biomedical Informatics for Computer-Aided Decision Support Systems: A Survey Ashwin Belle, Mark A. Kon, Kayvan Najarian The Scientific World Journal (2013) https://doi.org/10.1155/2013/769639

352. Advantages and disadvantages of using artificial neural networks versus logistic regression for predicting medical outcomes Jack V. Tu Journal of Clinical Epidemiology (1996-11) https://doi.org/10.1016/s0895-4356(96)00002-9

353. Use of an Artificial Neural Network for the Diagnosis of Myocardial Infarction William G. Baxt Annals of Internal Medicine (1991-12-01) https://doi.org/10.7326/0003-4819-115-11-843

354. Clinical Prediction Rules John H. Wasson, Harold C. Sox, Raymond K. Neff, Lee Goldman New England Journal of Medicine (1985-09-26) https://doi.org/10.1056/nejm198509263131306

355. The use of artificial neural networks in decision support in cancer: A systematic review Paulo J. Lisboa, Azzam F.G. Taktak Neural Networks (2006-05) https://doi.org/10.1016/j.neunet.2005.10.007

356. Estimating causal effects of treatments in randomized and nonrandomized studies. Donald B. Rubin Journal of Educational Psychology (1974) https://doi.org/10.1037/h0037350

357. Learning Representations for Counterfactual Inference Fredrik D. Johansson, Uri Shalit, David Sontag arXiv (2016-05-12) https://arxiv.org/abs/1605.03661v2

358. Causal Phenotype Discovery via Deep Networks David C. Kale, Zhengping Che, Mohammad Taha Bahadori, Wenzhe Li, Yan Liu, Randall Wetzel AMIA Annual Symposium Proceedings (2015) https://www.ncbi.nlm.nih.gov/pmc/articles/PMC4765623/

359. Modeling Missing Data in Clinical Time Series with RNNs Zachary C. Lipton, David C. Kale, Randall Wetzel arXiv (2016-06-13) https://arxiv.org/abs/1606.04130v5

360. Recurrent Neural Networks for Multivariate Time Series with Missing Values Zhengping Che, Sanjay Purushotham, Kyunghyun Cho, David Sontag, Yan Liu arXiv (2016-06-06) https://arxiv.org/abs/1606.01865v2

361. Predicting Complications in Critical Care Using Heterogeneous Clinical Data Vijay Huddar, Bapu Koundinya Desiraju, Vaibhav Rajan, Sakyajit Bhattacharya, Shourya Roy, Chandan K. Reddy IEEE Access (2016) https://doi.org/10.1109/access.2016.2618775

362. Phenotyping of Clinical Time Series with LSTM Recurrent Neural Networks Zachary C. Lipton, David C. Kale, Randall C. Wetzel arXiv (2015-10-26) https://arxiv.org/abs/1510.07641v2

363. Optimal medication dosing from suboptimal clinical examples: A deep reinforcement learning approach Shamim Nemati, Mohammad M. Ghassemi, Gari D. Clifford 2016 38th Annual International Conference of the IEEE Engineering in Medicine and Biology Society (EMBC) (2016-08) https://doi.org/10.1109/embc.2016.7591355

364. From vital signs to clinical outcomes for patients with sepsis: a machine learning basis for a clinical decision support system Eren Gultepe, Jeffrey P Green, Hien Nguyen, Jason Adams, Timothy Albertson, Ilias Tagkopoulos Journal of the American Medical Informatics Association (2014-03) https://doi.org/10.1136/amiajnl-2013-001815

365. Imaging-based enrichment criteria using deep learning algorithms for efficient clinical trials in mild cognitive impairment Vamsi K. Ithapu, Vikas Singh, Ozioma C. Okonkwo, Richard J. Chappell, N. Maritza Dowling, Sterling C. Johnson Alzheimer’s & Dementia (2015-12) https://doi.org/10.1016/j.jalz.2015.01.010

366. Integrated deep learned transcriptomic and structure-based predictor of clinical trials outcomes Artem V Artemov, Evgeny Putin, Quentin Vanhaelen, Alexander Aliper, Ivan V Ozerov, Alex Zhavoronkov Cold Spring Harbor Laboratory (2016-12-20) https://doi.org/10.1101/095653

367. Innovation in the pharmaceutical industry: New estimates of R&D costs Joseph A. DiMasi, Henry G. Grabowski, Ronald W. Hansen Journal of Health Economics (2016-05) https://doi.org/10.1016/j.jhealeco.2016.01.012

368. An analysis of the attrition of drug candidates from four major pharmaceutical companies Michael J. Waring, John Arrowsmith, Andrew R. Leach, Paul D. Leeson, Sam Mandrell, Robert M. Owen, Garry Pairaudeau, William D. Pennie, Stephen D. Pickett, Jibo Wang, … Alex Weir Nature Reviews Drug Discovery (2015-06-19) https://doi.org/10.1038/nrd4609

369. The Connectivity Map: Using Gene-Expression Signatures to Connect Small Molecules, Genes, and Disease J. Lamb Science (2006-09-29) https://doi.org/10.1126/science.1132939

370. A survey of current trends in computational drug repositioning Jiao Li, Si Zheng, Bin Chen, Atul J. Butte, S. Joshua Swamidass, Zhiyong Lu Briefings in Bioinformatics (2015-03-31) https://doi.org/10.1093/bib/bbv020

371. A review of connectivity map and computational approaches in pharmacogenomics Aliyu Musa, Laleh Soltan Ghoraie, Shu-Dong Zhang, Galina Galzko, Olli Yli-Harja, Matthias Dehmer, Benjamin Haibe-Kains, Frank Emmert-Streib Briefings in Bioinformatics (2017-01-09) https://doi.org/10.1093/bib/bbw112

372. A review of validation strategies for computational drug repositioning Adam S. Brown, Chirag J. Patel Briefings in Bioinformatics (2016-11-22) https://academic.oup.com/bib/article/doi/10.1093/bib/bbw110/2562646/A-review-of-validation-strategies-for

373. Drug Repositioning: A Machine-Learning Approach through Data Integration Francesco Napolitano, Yan Zhao, Vania M Moreira, Roberto Tagliaferri, Juha Kere, Mauro D’Amato, Dario Greco Journal of Cheminformatics (2013) https://doi.org/10.1186/1758-2946-5-30

374. Drug–Disease Association and Drug-Repositioning Predictions in Complex Diseases Using Causal Inference–Probabilistic Matrix Factorization Jihong Yang, Zheng Li, Xiaohui Fan, Yiyu Cheng Journal of Chemical Information and Modeling (2014-09-22) https://doi.org/10.1021/ci500340n

375. Drug repositioning for non-small cell lung cancer by using machine learning algorithms and topological graph theory Chien-Hung Huang, Peter Mu-Hsin Chang, Chia-Wei Hsu, Chi-Ying F. Huang, Ka-Lok Ng BMC Bioinformatics (2016-01-11) https://doi.org/10.1186/s12859-015-0845-0

376. Machine Learning Prediction of Cancer Cell Sensitivity to Drugs Based on Genomic and Chemical Properties Michael P. Menden, Francesco Iorio, Mathew Garnett, Ultan McDermott, Cyril H. Benes, Pedro J. Ballester, Julio Saez-Rodriguez PLoS ONE (2013-04-30) https://doi.org/10.1371/journal.pone.0061318

377. Large-scale integration of small-molecule induced genome-wide transcriptional responses, Kinome-wide binding affinities and cell-growth inhibition profiles reveal global trends characterizing systems-level drug action Dušica Vidovic, Amar Koleti, Stephan C Schürer Frontiers in Genetics https://doi.org/10.3389/fgene.2014.00342

378. Computational Discovery of Putative Leads for Drug Repositioning through Drug-Target Interaction Prediction Edgar D. Coelho, Joel P. Arrais, José Luís Oliveira PLOS Computational Biology (2016-11-28) https://doi.org/10.1371/journal.pcbi.1005219

379. Large-Scale Off-Target Identification Using Fast and Accurate Dual Regularized One-Class Collaborative Filtering and Its Application to Drug Repurposing Hansaim Lim, Aleksandar Poleksic, Yuan Yao, Hanghang Tong, Di He, Luke Zhuang, Patrick Meng, Lei Xie PLOS Computational Biology (2016-10-07) https://doi.org/10.1371/journal.pcbi.1005135

380. Pairwise input neural network for target-ligand interaction prediction Caihua Wang, Juan Liu, Fei Luo, Yafang Tan, Zixin Deng, Qian-Nan Hu 2014 IEEE International Conference on Bioinformatics and Biomedicine (BIBM) (2014-11) https://doi.org/10.1109/bibm.2014.6999129

381. L1000CDS2: LINCS L1000 characteristic direction signatures search engine Qiaonan Duan, St Patrick Reid, Neil R Clark, Zichen Wang, Nicolas F Fernandez, Andrew D Rouillard, Ben Readhead, Sarah R Tritsch, Rachel Hodos, Marc Hafner, … Avi Ma’ayan npj Systems Biology and Applications (2016-08-04) https://doi.org/10.1038/npjsba.2016.15

382. A guide to drug discovery: Hit and lead generation: beyond high-throughput screening Konrad H. Bleicher, Hans-Joachim Böhm, Klaus Müller, Alexander I. Alanine Nature Reviews Drug Discovery (2003-05) https://doi.org/10.1038/nrd1086

383. Hit discovery and hit-to-lead approaches György M. Keserű, Gergely M. Makara Drug Discovery Today (2006-08) https://doi.org/10.1016/j.drudis.2006.06.016

384. Influence Relevance Voting: An Accurate And Interpretable Virtual High Throughput Screening Method S. Joshua Swamidass, Chloé-Agathe Azencott, Ting-Wan Lin, Hugo Gramajo, Shiou-Chuan Tsai, Pierre Baldi Journal of Chemical Information and Modeling (2009-04-27) https://doi.org/10.1021/ci8004379

385. Modeling Industrial ADMET Data with Multitask Networks Steven Kearnes, Brian Goldman, Vijay Pande arXiv (2016-06-28) https://arxiv.org/abs/1606.08793v3

386. XenoSite: Accurately Predicting CYP-Mediated Sites of Metabolism with Neural Networks Jed Zaretzki, Matthew Matlock, S. Joshua Swamidass Journal of Chemical Information and Modeling (2013-12-23) https://doi.org/10.1021/ci400518g

387. Multi-task Neural Networks for QSAR Predictions George E. Dahl, Navdeep Jaitly, Ruslan Salakhutdinov arXiv (2014-06-04) https://arxiv.org/abs/1406.1231v1

388. Deep Neural Nets as a Method for Quantitative Structure–Activity Relationships Junshui Ma, Robert P. Sheridan, Andy Liaw, George E. Dahl, Vladimir Svetnik Journal of Chemical Information and Modeling (2015-02-23) https://doi.org/10.1021/ci500747n

389. Did Kaggle Predict Drug Candidate Activities? Or Not? Derek Lowe In the Pipeline (2012-12-11) http://blogs.sciencemag.org/pipeline/archives/2012/12/11/did_kaggle_predict_drug_candidate_activities_or_not

390. Deep learning as an opportunity in virtual screening Thomas Unterthiner, Andreas Mayr, Günter Klambauer, Marvin Steijaert, Jörg K. Wegner, Hugo Ceulemans, Sepp Hochreiter Neural Information Processing Systems 2014: Deep Learning and Representation Learning Workshop (2014) http://www.dlworkshop.org/23.pdf?attredirects=0

391. Massively Multitask Networks for Drug Discovery Bharath Ramsundar, Steven Kearnes, Patrick Riley, Dale Webster, David Konerding, Vijay Pande arXiv (2015-02-06) https://arxiv.org/abs/1502.02072v1

392. DeepTox: Toxicity Prediction using Deep Learning Andreas Mayr, Günter Klambauer, Thomas Unterthiner, Sepp Hochreiter Frontiers in Environmental Science (2016-02-02) https://doi.org/10.3389/fenvs.2015.00080

393. Computational Modeling of β-Secretase 1 (BACE-1) Inhibitors Using Ligand Based Approaches Govindan Subramanian, Bharath Ramsundar, Vijay Pande, Rajiah Aldrin Denny Journal of Chemical Information and Modeling (2016-10-24) https://doi.org/10.1021/acs.jcim.6b00290

394. The enumeration of chemical space Jean-Louis Reymond, Lars Ruddigkeit, Lorenz Blum, Ruud van Deursen Wiley Interdisciplinary Reviews: Computational Molecular Science (2012-04-18) https://doi.org/10.1002/wcms.1104

395. Accurate and efficient target prediction using a potency-sensitive influence-relevance voter Alessandro Lusci, David Fooshee, Michael Browning, Joshua Swamidass, Pierre Baldi Journal of Cheminformatics (2015-12) https://doi.org/10.1186/s13321-015-0110-6

396. Molecular Descriptors for Chemoinformatics Methods and Principles in Medicinal Chemistry (2009-07-15) https://doi.org/10.1002/9783527628766

397. Extended-Connectivity Fingerprints David Rogers, Mathew Hahn Journal of Chemical Information and Modeling (2010-05-24) https://doi.org/10.1021/ci100050t

398. Automatic chemical design using a data-driven continuous representation of molecules Rafael Gómez-Bombarelli, David Duvenaud, José Miguel Hernández-Lobato, Jorge Aguilera-Iparraguirre, Timothy D. Hirzel, Ryan P. Adams, Alán Aspuru-Guzik arXiv (2016-10-07) https://arxiv.org/abs/1610.02415v1

399. Chemception: A Deep Neural Network with Minimal Chemistry Knowledge Matches the Performance of Expert-developed QSAR/QSPR Models Garrett B. Goh, Charles Siegel, Abhinav Vishnu, Nathan O. Hodas, Nathan Baker arXiv (2017-06-20) https://arxiv.org/abs/1706.06689v1

400. Convolutional Networks on Graphs for Learning Molecular Fingerprints David K. Duvenaud, Dougal Maclaurin, Jorge Iparraguirre, Rafael Bombarell, Timothy Hirzel, Alan Aspuru-Guzik, Ryan P. Adams (2015) http://papers.nips.cc/paper/5954-convolutional-networks-on-graphs-for-learning-molecular-fingerprints

401. Deep Architectures and Deep Learning in Chemoinformatics: The Prediction of Aqueous Solubility for Drug-Like Molecules Alessandro Lusci, Gianluca Pollastri, Pierre Baldi Journal of Chemical Information and Modeling (2013-07-22) https://doi.org/10.1021/ci400187y

402. Molecular graph convolutions: moving beyond fingerprints Steven Kearnes, Kevin McCloskey, Marc Berndl, Vijay Pande, Patrick Riley Journal of Computer-Aided Molecular Design (2016-08) https://doi.org/10.1007/s10822-016-9938-8

403. Low Data Drug Discovery with One-Shot Learning Han Altae-Tran, Bharath Ramsundar, Aneesh S. Pappu, Vijay Pande ACS Central Science (2017-04-03) https://doi.org/10.1021/acscentsci.6b00367

404. Convolutional Embedding of Attributed Molecular Graphs for Physical Property Prediction Connor W. Coley, Regina Barzilay, William H. Green, Tommi S. Jaakkola, Klavs F. Jensen Journal of Chemical Information and Modeling (2017-07-25) https://doi.org/10.1021/acs.jcim.6b00601

405. Learning a Local-Variable Model of Aromatic and Conjugated Systems Matthew K. Matlock, Na Le Dang, S. Joshua Swamidass ACS Central Science (2018-01-03) https://doi.org/10.1021/acscentsci.7b00405

406. Covariant Compositional Networks For Learning Graphs Risi Kondor, Hy Truong Son, Horace Pan, Brandon Anderson, Shubhendu Trivedi arXiv (2018-01-07) https://arxiv.org/abs/1801.02144v1

407. MoleculeNet: a benchmark for molecular machine learning Zhenqin Wu, Bharath Ramsundar, Evan N. Feinberg, Joseph Gomes, Caleb Geniesse, Aneesh S. Pappu, Karl Leswing, Vijay Pande Chemical Science (2018) https://doi.org/10.1039/c7sc02664a

408. What do we know and when do we know it? Anthony Nicholls Journal of Computer-Aided Molecular Design (2008-02-06) https://doi.org/10.1007/s10822-008-9170-2

409. deepchem/deepchem GitHub (2017) https://github.com/deepchem/deepchem

410. Mol2vec: Unsupervised Machine Learning Approach with Chemical Intuition Sabrina Jaeger, Simone Fulle, Samo Turk Journal of Chemical Information and Modeling (2018-01-10) https://doi.org/10.1021/acs.jcim.7b00616

411. Structure-Based Virtual Screening for Drug Discovery: a Problem-Centric Review Tiejun Cheng, Qingliang Li, Zhigang Zhou, Yanli Wang, Stephen H. Bryant The AAPS Journal (2012-01-27) https://doi.org/10.1208/s12248-012-9322-0

412. Atomic Convolutional Networks for Predicting Protein-Ligand Binding Affinity Joseph Gomes, Bharath Ramsundar, Evan N. Feinberg, Vijay S. Pande arXiv (2017-03-30) https://arxiv.org/abs/1703.10603v1

413. TopologyNet: Topology based deep convolutional and multi-task neural networks for biomolecular property predictions Zixuan Cang, Guo-Wei Wei PLOS Computational Biology (2017-07-27) https://doi.org/10.1371/journal.pcbi.1005690

414. The PDBbind Database: Methodologies and Updates Renxiao Wang, Xueliang Fang, Yipin Lu, Chao-Yie Yang, Shaomeng Wang Journal of Medicinal Chemistry (2005-06) https://doi.org/10.1021/jm048957q

415. Boosting Docking-Based Virtual Screening with Deep Learning Janaina Cruz Pereira, Ernesto Raúl Caffarena, Cicero Nogueira dos Santos Journal of Chemical Information and Modeling (2016-12-27) https://doi.org/10.1021/acs.jcim.6b00355

416. Protein-Ligand Scoring with Convolutional Neural Networks Matthew Ragoza, Joshua Hochuli, Elisa Idrobo, Jocelyn Sunseri, David Ryan Koes arXiv (2016-12-08) https://arxiv.org/abs/1612.02751v1

417. Enabling future drug discovery byde novodesign Markus Hartenfeller, Gisbert Schneider Wiley Interdisciplinary Reviews: Computational Molecular Science (2011-04-25) https://doi.org/10.1002/wcms.49

418. De Novo Design at the Edge of Chaos Petra Schneider, Gisbert Schneider Journal of Medicinal Chemistry (2016-05-12) https://doi.org/10.1021/acs.jmedchem.5b01849

419. Generating Sequences With Recurrent Neural Networks Alex Graves arXiv (2013-08-04) https://arxiv.org/abs/1308.0850v5

420. Generating Focussed Molecule Libraries for Drug Discovery with Recurrent Neural Networks Marwin H. S. Segler, Thierry Kogej, Christian Tyrchan, Mark P. Waller arXiv (2017-01-05) https://arxiv.org/abs/1701.01329v1

421. Grammar Variational Autoencoder Matt J. Kusner, Brooks Paige, José Miguel Hernández-Lobato arXiv (2017-03-06) https://arxiv.org/abs/1703.01925v1

422. ChEMBL: a large-scale bioactivity database for drug discovery A. Gaulton, L. J. Bellis, A. P. Bento, J. Chambers, M. Davies, A. Hersey, Y. Light, S. McGlinchey, D. Michalovich, B. Al-Lazikani, J. P. Overington Nucleic Acids Research (2011-09-23) https://doi.org/10.1093/nar/gkr777

423. Molecular De Novo Design through Deep Reinforcement Learning Marcus Olivecrona, Thomas Blaschke, Ola Engkvist, Hongming Chen arXiv (2017-04-25) https://arxiv.org/abs/1704.07555v2

424. Sequence Tutor: Conservative Fine-Tuning of Sequence Generation Models with KL-control Natasha Jaques, Shixiang Gu, Dzmitry Bahdanau, José Miguel Hernández-Lobato, Richard E. Turner, Douglas Eck arXiv (2016-11-09) https://arxiv.org/abs/1611.02796v9

425. Understanding deep learning requires rethinking generalization Chiyuan Zhang, Samy Bengio, Moritz Hardt, Benjamin Recht, Oriol Vinyals arXiv (2016-11-10) https://arxiv.org/abs/1611.03530v2

426. Why does deep and cheap learning work so well? Henry W. Lin, Max Tegmark, David Rolnick arXiv (2016-08-29) https://arxiv.org/abs/1608.08225v3

427. The relationship between Precision-Recall and ROC curves Jesse Davis, Mark Goadrich Proceedings of the 23rd international conference on Machine learning-ICML’06 (2006) https://doi.org/10.1145/1143844.1143874

428. An open investigation of the reproducibility of cancer biology research Timothy M Errington, Elizabeth Iorns, William Gunn, Fraser Elisabeth Tan, Joelle Lomax, Brian A Nosek eLife (2014-12-10) https://doi.org/10.7554/elife.04333

429. Adversarial Examples, Uncertainty, and Transfer Testing Robustness in Gaussian Process Hybrid Deep Networks John Bradshaw, Alexander G. de G. Matthews, Zoubin Ghahramani arXiv (2017-07-08) https://arxiv.org/abs/1707.02476v1

430. What Uncertainties Do We Need in Bayesian Deep Learning for Computer Vision? Alex Kendall, Yarin Gal arXiv (2017-03-15) https://arxiv.org/abs/1703.04977v2

431. Multi-Task Learning Using Uncertainty to Weigh Losses for Scene Geometry and Semantics Alex Kendall, Yarin Gal, Roberto Cipolla arXiv (2017-05-19) https://arxiv.org/abs/1705.07115v1

432. On Calibration of Modern Neural Networks Chuan Guo, Geoff Pleiss, Yu Sun, Kilian Q. Weinberger arXiv (2017-06-14) https://arxiv.org/abs/1706.04599v2

433. Probabilistic Outputs for Support Vector Machines and Comparisons to Regularized Likelihood Methods John C. Platt ADVANCES IN LARGE MARGIN CLASSIFIERS http://citeseer.ist.psu.edu/viewdoc/summary?doi=10.1.1.41.1639

434. Confidence interval prediction for neural network models G. Chryssolouris, M. Lee, A. Ramsey IEEE Transactions on Neural Networks (1996) https://doi.org/10.1109/72.478409

435. A Baseline for Detecting Misclassified and Out-of-Distribution Examples in Neural Networks Dan Hendrycks, Kevin Gimpel arXiv (2016-10-07) https://arxiv.org/abs/1610.02136v2

436. Enhancing The Reliability of Out-of-distribution Image Detection in Neural Networks Shiyu Liang, Yixuan Li, R. Srikant arXiv (2017-06-08) https://arxiv.org/abs/1706.02690v3

437. Concrete Problems in AI Safety Dario Amodei, Chris Olah, Jacob Steinhardt, Paul Christiano, John Schulman, Dan Mané arXiv (2016-06-21) https://arxiv.org/abs/1606.06565v2

438. Adversarial Examples Are Not Easily Detected: Bypassing Ten Detection Methods Nicholas Carlini, David Wagner arXiv (2017-05-20) https://arxiv.org/abs/1705.07263v2

439. Dropout as a Bayesian Approximation: Representing Model Uncertainty in Deep Learning Yarin Gal, Zoubin Ghahramani arXiv (2015-06-06) https://arxiv.org/abs/1506.02142v6

440. Leveraging uncertainty information from deep neural networks for disease detection Christian Leibig, Vaneeda Allken, Murat Seçkin Ayhan, Philipp Berens, Siegfried Wahl Scientific Reports (2017-12) https://doi.org/10.1038/s41598-017-17876-z

441. Robustly representing inferential uncertainty in deep neural networks through sampling Patrick McClure, Nikolaus Kriegeskorte arXiv (2016-11-05) https://arxiv.org/abs/1611.01639v6

442. Bayesian Hypernetworks David Krueger, Chin-Wei Huang, Riashat Islam, Ryan Turner, Alexandre Lacoste, Aaron Courville arXiv (2017-10-13) https://arxiv.org/abs/1710.04759v1

443. Simple and Scalable Predictive Uncertainty Estimation using Deep Ensembles Balaji Lakshminarayanan, Alexander Pritzel, Charles Blundell arXiv (2016-12-05) https://arxiv.org/abs/1612.01474v3

444. Uncertainty in deep learning Yarin Gal PhD thesis, University of Cambridge (2016) http://www.cs.ox.ac.uk/people/yarin.gal/website/thesis/thesis.pdf

445. Do Deep Nets Really Need to be Deep? Lei Jimmy Ba, Rich Caruana arXiv (2013-12-21) https://arxiv.org/abs/1312.6184v7

446. Deep Neural Networks are Easily Fooled: High Confidence Predictions for Unrecognizable Images Anh Nguyen, Jason Yosinski, Jeff Clune arXiv (2014-12-05) https://arxiv.org/abs/1412.1897v4

447. “Why Should I Trust You?”: Explaining the Predictions of Any Classifier Marco Tulio Ribeiro, Sameer Singh, Carlos Guestrin arXiv (2016-02-16) https://arxiv.org/abs/1602.04938v3

448. Visualizing and Understanding Convolutional Networks Matthew D Zeiler, Rob Fergus arXiv (2013-11-12) https://arxiv.org/abs/1311.2901v3

449. Visualizing Deep Neural Network Decisions: Prediction Difference Analysis Luisa M Zintgraf, Taco S Cohen, Tameem Adel, Max Welling arXiv (2017-02-15) https://arxiv.org/abs/1702.04595v1

450. Interpretable Explanations of Black Boxes by Meaningful Perturbation Ruth C. Fong, Andrea Vedaldi 2017 IEEE International Conference on Computer Vision (ICCV) (2017-10) https://doi.org/10.1109/iccv.2017.371

451. Deep Inside Convolutional Networks: Visualising Image Classification Models and Saliency Maps Karen Simonyan, Andrea Vedaldi, Andrew Zisserman arXiv (2013-12-20) https://arxiv.org/abs/1312.6034v2

452. On Pixel-Wise Explanations for Non-Linear Classifier Decisions by Layer-Wise Relevance Propagation Sebastian Bach, Alexander Binder, Grégoire Montavon, Frederick Klauschen, Klaus-Robert Müller, Wojciech Samek PLOS ONE (2015-07-10) https://doi.org/10.1371/journal.pone.0130140

453. Investigating the influence of noise and distractors on the interpretation of neural networks Pieter-Jan Kindermans, Kristof Schütt, Klaus-Robert Müller, Sven Dähne arXiv (2016-11-22) https://arxiv.org/abs/1611.07270v1

454. Striving for Simplicity: The All Convolutional Net Jost Tobias Springenberg, Alexey Dosovitskiy, Thomas Brox, Martin Riedmiller arXiv (2014-12-21) https://arxiv.org/abs/1412.6806v3

455. Salient Deconvolutional Networks Aravindh Mahendran, Andrea Vedaldi Computer Vision – ECCV 2016 (2016) https://doi.org/10.1007/978-3-319-46466-4_8

456. Grad-CAM: Visual Explanations from Deep Networks via Gradient-based Localization Ramprasaath R. Selvaraju, Michael Cogswell, Abhishek Das, Ramakrishna Vedantam, Devi Parikh, Dhruv Batra arXiv (2016-10-07) https://arxiv.org/abs/1610.02391v3

457. Axiomatic Attribution for Deep Networks Mukund Sundararajan, Ankur Taly, Qiqi Yan arXiv (2017-03-04) https://arxiv.org/abs/1703.01365v2

458. An unexpected unity among methods for interpreting model predictions Scott Lundberg, Su-In Lee arXiv (2016-11-22) https://arxiv.org/abs/1611.07478v3

459. 17. A Value for n-Person Games L. S. Shapley Contributions to the Theory of Games (AM-28), Volume II (1953) https://doi.org/10.1515/9781400881970-018

460. Understanding Deep Image Representations by Inverting Them Aravindh Mahendran, Andrea Vedaldi arXiv (2014-11-26) https://arxiv.org/abs/1412.0035v1

461. Maximum Entropy Methods for Extracting the Learned Features of Deep Neural Networks Alex I Finnegan, Jun S Song Cold Spring Harbor Laboratory (2017-02-03) https://doi.org/10.1101/105957

462. Visualizing Deep Convolutional Neural Networks Using Natural Pre-images Aravindh Mahendran, Andrea Vedaldi International Journal of Computer Vision (2016-05-18) https://doi.org/10.1007/s11263-016-0911-8

463. Inceptionism: Going Deeper into Neural Networks Alexander Mordvintsev, Christopher Olah, Mike Tyka Google Research Blog (2015-06) http://googleresearch.blogspot.co.uk/2015/06/inceptionism-going-deeper-into-neural.html

464. Visualizing Higher-Layer Features of a Deep Network Dumitru Erhan, Yoshua Bengio, Aaron Courville, Pascal Vincent University of Montreal (2009-06) http://www.iro.umontreal.ca/~lisa/publications2/index.php/publications/show/247

465. Understanding Neural Networks Through Deep Visualization Jason Yosinski, Jeff Clune, Anh Nguyen, Thomas Fuchs, Hod Lipson arXiv (2015-06-22) https://arxiv.org/abs/1506.06579v1

466. Neural Machine Translation by Jointly Learning to Align and Translate Dzmitry Bahdanau, Kyunghyun Cho, Yoshua Bengio arXiv (2014-09-01) https://arxiv.org/abs/1409.0473v7

467. Show, Attend and Tell: Neural Image Caption Generation with Visual Attention Kelvin Xu, Jimmy Ba, Ryan Kiros, Kyunghyun Cho, Aaron Courville, Ruslan Salakhutdinov, Richard Zemel, Yoshua Bengio arXiv (2015-02-10) https://arxiv.org/abs/1502.03044v3

468. Genetic Architect: Discovering Genomic Structure with Learned Neural Architectures Laura Deming, Sasha Targ, Nate Sauder, Diogo Almeida, Chun Jimmie Ye arXiv (2016-05-23) https://arxiv.org/abs/1605.07156v1

469. RETAIN: An Interpretable Predictive Model for Healthcare using Reverse Time Attention Mechanism Edward Choi, Mohammad Taha Bahadori, Joshua A. Kulas, Andy Schuetz, Walter F. Stewart, Jimeng Sun arXiv (2016-08-19) https://arxiv.org/abs/1608.05745v4

470. GRAM: Graph-based Attention Model for Healthcare Representation Learning Edward Choi, Mohammad Taha Bahadori, Le Song, Walter F. Stewart, Jimeng Sun arXiv (2016-11-21) https://arxiv.org/abs/1611.07012v3

471. Sequence learning with recurrent networks: analysis of internal representations Joydeep Ghosh, Vijay Karamcheti Science of Artificial Neural Networks (1992-07-01) https://doi.org/10.1117/12.140112

472. Visualizing and Understanding Recurrent Networks Andrej Karpathy, Justin Johnson, Li Fei-Fei arXiv (2015-06-05) https://arxiv.org/abs/1506.02078v2

473. LSTMVis: A Tool for Visual Analysis of Hidden State Dynamics in Recurrent Neural Networks Hendrik Strobelt, Sebastian Gehrmann, Hanspeter Pfister, Alexander M. Rush arXiv (2016-06-23) https://arxiv.org/abs/1606.07461v2

474. Automatic Rule Extraction from Long Short Term Memory Networks W. James Murdoch, Arthur Szlam arXiv (2017-02-08) https://arxiv.org/abs/1702.02540v2

475. Unsupervised Representation Learning with Deep Convolutional Generative Adversarial Networks Alec Radford, Luke Metz, Soumith Chintala arXiv (2015-11-19) https://arxiv.org/abs/1511.06434v2

476. The Cancer Genome Atlas Pan-Cancer analysis project Kyle Chang, Chad J Creighton, Caleb Davis, Lawrence Donehower, Jennifer Drummond, David Wheeler, Adrian Ally, Miruna Balasundaram, Inanc Birol, Yaron SN Butterfield, … Joshua M Stuart Nature Genetics (2013-09-26) https://doi.org/10.1038/ng.2764

477. Extracting a Biologically Relevant Latent Space from Cancer Transcriptomes with Variational Autoencoders Gregory P. Way, Casey S. Greene Cold Spring Harbor Laboratory (2017-08-11) https://doi.org/10.1101/174474

478. Evaluating deep variational autoencoders trained on pan-cancer gene expression Gregory P. Way, Casey S. Greene arXiv (2017-11-13) https://arxiv.org/abs/1711.04828v1

479. GANs for Biological Image Synthesis Anton Osokin, Anatole Chessel, Rafael E. Carazo Salas, Federico Vaggi arXiv (2017-08-15) https://arxiv.org/abs/1708.04692v2

480. CytoGAN: Generative Modeling of Cell Images Peter Goldsborough, Nick Pawlowski, Juan C Caicedo, Shantanu Singh, Anne Carpenter Cold Spring Harbor Laboratory (2017-12-02) https://doi.org/10.1101/227645

481. Understanding Black-box Predictions via Influence Functions Pang Wei Koh, Percy Liang arXiv (2017-03-14) https://arxiv.org/abs/1703.04730v2

482. ActiVis: Visual Exploration of Industry-Scale Deep Neural Network Models Minsuk Kahng, Pierre Y. Andrews, Aditya Kalro, Duen Horng Chau arXiv (2017-04-06) https://arxiv.org/abs/1704.01942v2

483. Towards Better Analysis of Deep Convolutional Neural Networks Mengchen Liu, Jiaxin Shi, Zhen Li, Chongxuan Li, Jun Zhu, Shixia Liu arXiv (2016-04-24) https://arxiv.org/abs/1604.07043v3

484. Distilling Knowledge from Deep Networks with Applications to Healthcare Domain Zhengping Che, Sanjay Purushotham, Robinder Khemani, Yan Liu arXiv (2015-12-11) https://arxiv.org/abs/1512.03542v1

485. Rationalizing Neural Predictions Tao Lei, Regina Barzilay, Tommi Jaakkola arXiv (2016-06-13) https://arxiv.org/abs/1606.04155v2

486. Learning multiple layers of features from tiny images Alex Krizhevsky (2009) https://www.cs.toronto.edu/~kriz/learning-features-2009-TR.pdf

487. Functional Knowledge Transfer for High-accuracy Prediction of Under-studied Biological Processes Christopher Y. Park, Aaron K. Wong, Casey S. Greene, Jessica Rowland, Yuanfang Guan, Lars A. Bongo, Rebecca D. Burdine, Olga G. Troyanskaya PLoS Computational Biology (2013-03-14) https://doi.org/10.1371/journal.pcbi.1002957

488. DeepAD: Alzheimer’s Disease Classification via Deep Convolutional Neural Networks using MRI and fMRI Saman Sarraf, Danielle D. DeSouza, John Anderson, Ghassem Tofighi, Cold Spring Harbor Laboratory (2016-08-21) https://doi.org/10.1101/070441

489. DeepBound: Accurate Identification of Transcript Boundaries via Deep Convolutional Neural Fields Mingfu Shao, Jianzhu Ma, Sheng Wang Cold Spring Harbor Laboratory (2017-04-07) https://doi.org/10.1101/125229

490. A general framework for estimating the relative pathogenicity of human genetic variants Martin Kircher, Daniela M Witten, Preti Jain, Brian J O’Roak, Gregory M Cooper, Jay Shendure Nature Genetics (2014-02-02) https://doi.org/10.1038/ng.2892

491. Diet Networks: Thin Parameters for Fat Genomics Adriana Romero, Pierre Luc Carrier, Akram Erraqabi, Tristan Sylvain, Alex Auvolat, Etienne Dejoie, Marc-André Legault, Marie-Pierre Dubé, Julie G. Hussin, Yoshua Bengio International Conference on Learning Representations 2017 (2016-11-04) https://openreview.net/forum?id=Sk-oDY9ge&noteId=Sk-oDY9ge

492. Deep learning in neural networks: An overview Jürgen Schmidhuber Neural Networks (2015-01) https://doi.org/10.1016/j.neunet.2014.09.003

493. Deep Learning with Limited Numerical Precision Suyog Gupta, Ankur Agrawal, Kailash Gopalakrishnan, Pritish Narayanan arXiv (2015-02-09) https://arxiv.org/abs/1502.02551v1

494. Training deep neural networks with low precision multiplications Matthieu Courbariaux, Yoshua Bengio, Jean-Pierre David arXiv (2014-12-22) https://arxiv.org/abs/1412.7024v5

495. Taming the Wild: A Unified Analysis of Hogwild!-Style Algorithms Christopher De Sa, Ce Zhang, Kunle Olukotun, Christopher Ré arXiv (2015-06-22) https://arxiv.org/abs/1506.06438v2

496. Quantized Neural Networks: Training Neural Networks with Low Precision Weights and Activations Itay Hubara, Matthieu Courbariaux, Daniel Soudry, Ran El-Yaniv, Yoshua Bengio arXiv (2016-09-22) https://arxiv.org/abs/1609.07061v1

497. Distilling the Knowledge in a Neural Network Geoffrey Hinton, Oriol Vinyals, Jeff Dean arXiv (2015-03-09) https://arxiv.org/abs/1503.02531v1

498. Large-scale deep unsupervised learning using graphics processors Rajat Raina, Anand Madhavan, Andrew Y. Ng Proceedings of the 26th Annual International Conference on Machine Learning-ICML’09 (2009) https://doi.org/10.1145/1553374.1553486

499. Improving the speed of neural networks on CPUs Vincent Vanhoucke, Andrew Senior, Mark Z. Mao (2011) https://research.google.com/pubs/pub37631.html

500. On parallelizability of stochastic gradient descent for speech DNNS Frank Seide, Hao Fu, Jasha Droppo, Gang Li, Dong Yu 2014 IEEE International Conference on Acoustics, Speech and Signal Processing (ICASSP) (2014-05) https://doi.org/10.1109/icassp.2014.6853593

501. Caffe con Troll: Shallow Ideas to Speed Up Deep Learning Stefan Hadjis, Firas Abuzaid, Ce Zhang, Christopher Ré arXiv (2015-04-16) https://arxiv.org/abs/1504.04343v2

502. Growing pains for deep learning Chris Edwards Communications of the ACM (2015-06-25) href="https://doi.org/10.1145/2771283

503. Experiments on Parallel Training of Deep Neural Network using Model Averaging Hang Su, Haoyu Chen arXiv (2015-07-05) https://arxiv.org/abs/1507.01239v2

504. Efficient mini-batch training for stochastic optimization Mu Li, Tong Zhang, Yuqiang Chen, Alexander J. Smola Proceedings of the 20th ACM SIGKDD international conference on Knowledge discovery and data mining-KDD’14 (2014) https://doi.org/10.1145/2623330.2623612

505. CGBVS-DNN: Prediction of Compound-protein Interactions Based on Deep Learning Masatoshi Hamanaka, Kei Taneishi, Hiroaki Iwata, Jun Ye, Jianguo Pei, Jinlong Hou, Yasushi Okuno Molecular Informatics (2016-08-12) https://doi.org/10.1002/minf.201600045

506. cuDNN: Efficient Primitives for Deep Learning Sharan Chetlur, Cliff Woolley, Philippe Vandermersch, Jonathan Cohen, John Tran, Bryan Catanzaro, Evan Shelhamer arXiv (2014-10-03) https://arxiv.org/abs/1410.0759v3

507. Compressing Neural Networks with the Hashing Trick Wenlin Chen, James T. Wilson, Stephen Tyree, Kilian Q. Weinberger, Yixin Chen arXiv (2015-04-19) https://arxiv.org/abs/1504.04788v1

508. Deep Learning on FPGAs: Past, Present, and Future Griffin Lacey, Graham W. Taylor, Shawki Areibi arXiv (2016-02-13) https://arxiv.org/abs/1602.04283v1

509. In-Datacenter Performance Analysis of a Tensor Processing Unit Norman P. Jouppi, Cliff Young, Nishant Patil, David Patterson, Gaurav Agrawal, Raminder Bajwa, Sarah Bates, Suresh Bhatia, Nan Boden, Al Borchers, … Doe Hyun Yoon arXiv (2017-04-16) https://arxiv.org/abs/1704.04760v1

510. MapReduce Jeffrey Dean, Sanjay Ghemawat Communications of the ACM (2008-01-01) https://doi.org/10.1145/1327452.1327492

511. Distributed GraphLab Yucheng Low, Danny Bickson, Joseph Gonzalez, Carlos Guestrin, Aapo Kyrola, Joseph M. Hellerstein Proceedings of the VLDB Endowment (2012-04-01) https://doi.org/10.14778/2212351.2212354

512. Large Scale Distributed Deep Networks Jeffrey Dean, Greg S Corrado, Rajat Monga, Kai Chen, Matthieu Devin, Quoc V Le, Mark Z Mao, Marc’Aurelio Ranzato, Andrew Senior, Paul Tucker, … Andrew Y Ng Neural Information Processing Systems 2012 (2012-12) http://research.google.com/archive/large_deep_networks_nips2012.html

513. Taming the Wild: A Unified Analysis of Hogwild!-Style Algorithms Christopher De Sa, Ce Zhang, Kunle Olukotun, Christopher Ré Advances in neural information processing systems (2015-12) https://www.ncbi.nlm.nih.gov/pmc/articles/PMC4907892/

514. SparkNet: Training Deep Networks in Spark Philipp Moritz, Robert Nishihara, Ion Stoica, Michael I. Jordan arXiv (2015-11-19) https://arxiv.org/abs/1511.06051v4

515. MLlib: Machine Learning in Apache Spark Xiangrui Meng, Joseph Bradley, Burak Yavuz, Evan Sparks, Shivaram Venkataraman, Davies Liu, Jeremy Freeman, DB Tsai, Manish Amde, Sean Owen, … Ameet Talwalkar arXiv (2015-05-26) https://arxiv.org/abs/1505.06807v1

516. TensorFlow: Large-Scale Machine Learning on Heterogeneous Distributed Systems Martín Abadi, Ashish Agarwal, Paul Barham, Eugene Brevdo, Zhifeng Chen, Craig Citro, Greg S. Corrado, Andy Davis, Jeffrey Dean, Matthieu Devin, … Xiaoqiang Zheng arXiv (2016-03-14) https://arxiv.org/abs/1603.04467v2

517. fchollet/keras GitHub (2017) https://github.com/fchollet/keras

518. maxpumperla/elephas GitHub (2017) https://github.com/maxpumperla/elephas

519. Deep learning with COTS HPC systems Adam Coates, Brody Huval, Tao Wang, David Wu, Bryan Catanzaro, Ng Andrew (2013-02-13) http://www.jmlr.org/proceedings/papers/v28/coates13.html

520. Ensemble-Compression: A New Method for Parallel Training of Deep Neural Networks Shizhao Sun, Wei Chen, Jiang Bian, Xiaoguang Liu, Tie-Yan Liu arXiv (2016-06-02) https://arxiv.org/abs/1606.00575v2

521. Algorithms for Hyper-parameter Optimization James Bergstra, Rémi Bardenet, Yoshua Bengio, Balázs Kégl Proceedings of the 24th International Conference on Neural Information Processing Systems (2011) http://dl.acm.org/citation.cfm?id=2986459.2986743

522. Random Search for Hyper-Parameter Optimization James Bergstra, Yoshua Bengio Journal of Machine Learning Research (2012) http://www.jmlr.org/papers/v13/bergstra12a.html

523. Cloud computing and the DNA data race Michael C Schatz, Ben Langmead, Steven L Salzberg Nature Biotechnology (2010-07) https://doi.org/10.1038/nbt0710-691

524. The real cost of sequencing: scaling computation to keep pace with data generation Paul Muir, Shantao Li, Shaoke Lou, Daifeng Wang, Daniel J Spakowicz, Leonidas Salichos, Jing Zhang, George M. Weinstock, Farren Isaacs, Joel Rozowsky, Mark Gerstein Genome Biology (2016-03-23) https://doi.org/10.1186/s13059-016-0917-0

525. The case for cloud computing in genome informatics Lincoln D Stein Genome Biology (2010) https://doi.org/10.1186/gb-2010-11-5-207

526. One weird trick for parallelizing convolutional neural networks Alex Krizhevsky arXiv (2014-04-23) https://arxiv.org/abs/1404.5997v2

527. A view of cloud computing Michael Armbrust, Ion Stoica, Matei Zaharia, Armando Fox, Rean Griffith, Anthony D. Joseph, Randy Katz, Andy Konwinski, Gunho Lee, David Patterson, Ariel Rabkin Communications of the ACM (2010-04-01) https://doi.org/10.1145/1721654.1721672

528. Data Sharing Dan L. Longo, Jeffrey M. Drazen New England Journal of Medicine (2016-01-21) https://doi.org/10.1056/nejme1516564

529. Celebrating parasites Casey S Greene, Lana X Garmire, Jack A Gilbert, Marylyn D Ritchie, Lawrence E Hunter Nature Genetics (2017-03-30) https://doi.org/10.1038/ng.3830

530. Is Multitask Deep Learning Practical for Pharma? Bharath Ramsundar, Bowen Liu, Zhenqin Wu, Andreas Verras, Matthew Tudor, Robert P. Sheridan, Vijay Pande Journal of Chemical Information and Modeling (2017-08) https://doi.org/10.1021/acs.jcim.7b00146

531. Enhancing reproducibility for computational methods V. Stodden, M. McNutt, D. H. Bailey, E. Deelman, Y. Gil, B. Hanson, M. A. Heroux, J. P. A. Ioannidis, M. Taufer Science (2016-12-08) https://doi.org/10.1126/science.aah6168

532. DragoNN(2016-11-06) http://kundajelab.github.io/dragonn/

533. How transferable are features in deep neural networks? Jason Yosinski, Jeff Clune, Yoshua Bengio, Hod Lipson (2014) https://papers.nips.cc/paper/5347-how-transferable-are-features-in-deep-neural-networks

534. Deep Model Based Transfer and Multi-Task Learning for Biological Image Analysis Wenlu Zhang, Rongjian Li, Tao Zeng, Qian Sun, Sudhir Kumar, Jieping Ye, Shuiwang Ji Proceedings of the 21th ACM SIGKDD International Conference on Knowledge Discovery and Data Mining-KDD’15 (2015) https://doi.org/10.1145/2783258.2783304

535. Deep convolutional neural networks for annotating gene expression patterns in the mouse brain Tao Zeng, Rongjian Li, Ravi Mukkamala, Jieping Ye, Shuiwang Ji BMC Bioinformatics (2015-05-07) https://doi.org/10.1186/s12859-015-0553-9

536. Accurate Classification of Protein Subcellular Localization from High-Throughput Microscopy Images Using Deep Learning Tanel Pärnamaa, Leopold Parts G3: Genes|Genomes|Genetics (2017-04-08) https://doi.org/10.1534/g3.116.033654

537. Automated analysis of high?content microscopy data with deep learning Oren Z Kraus, Ben T Grys, Jimmy Ba, Yolanda Chong, Brendan J Frey, Charles Boone, Brenda J Andrews Molecular Systems Biology (2017-04) https://doi.org/10.15252/msb.20177551

538. Multimodal Deep Learning Jiquan Ngiam, Aditya Khosla, Mingyu Kim, Juhan Nam, Honglak Lee, Andrew Y. Ng Proceedings of the 28th International Conference on Machine Learning (2011) https://ccrma.stanford.edu/~juhan/pubs/NgiamKhoslaKimNamLeeNg2011.pdf

539. Deep Learning based multi-omics integration robustly predicts survival in liver cancer Kumardeep Chaudhary, Olivier B. Poirion, Liangqun Lu, Lana X. Garmire Cold Spring Harbor Laboratory (2017-03-08) https://doi.org/10.1101/114892

540. FIDDLE: An integrative deep learning framework for functional genomic data inference Umut Eser, L. Stirling Churchman Cold Spring Harbor Laboratory (2016-10-17) https://doi.org/10.1101/081380

541. Modeling Reactivity to Biological Macromolecules with a Deep Multitask Network Tyler B. Hughes, Na Le Dang, Grover P. Miller, S. Joshua Swamidass ACS Central Science (2016-08-24) https://doi.org/10.1021/acscentsci.6b00162

542. IBM edges closer to human speech recognition BI Intelligence Business Insider (2017-03-14) http://www.businessinsider.com/ibm-edges-closer-to-human-speech-recognition-2017-3

543. Achieving Human Parity in Conversational Speech Recognition W. Xiong, J. Droppo, X. Huang, F. Seide, M. Seltzer, A. Stolcke, D. Yu, G. Zweig arXiv (2016-10-17) https://arxiv.org/abs/1610.05256v2

544. English Conversational Telephone Speech Recognition by Humans and Machines George Saon, Gakuto Kurata, Tom Sercu, Kartik Audhkhasi, Samuel Thomas, Dimitrios Dimitriadis, Xiaodong Cui, Bhuvana Ramabhadran, Michael Picheny, Lynn-Li Lim, … Phil Hall arXiv (2017-03-06) https://arxiv.org/abs/1703.02136v1

545. Intriguing properties of neural networks Christian Szegedy, Wojciech Zaremba, Ilya Sutskever, Joan Bruna, Dumitru Erhan, Ian Goodfellow, Rob Fergus arXiv (2013-12-21) https://arxiv.org/abs/1312.6199v4

546. Explaining and Harnessing Adversarial Examples Ian J. Goodfellow, Jonathon Shlens, Christian Szegedy arXiv (2014-12-20) https://arxiv.org/abs/1412.6572v3

547. Towards the Science of Security and Privacy in Machine Learning Nicolas Papernot, Patrick McDaniel, Arunesh Sinha, Michael Wellman arXiv (2016-11-11) https://arxiv.org/abs/1611.03814v1

548. Feature Squeezing: Detecting Adversarial Examples in Deep Neural Networks Weilin Xu, David Evans, Yanjun Qi arXiv (2017-04-04) https://arxiv.org/abs/1704.01155v1

549. The Grey Literature — Proof of prespecified endpoints in medical research with the bitcoin blockchain Benjamin Gregory Carlisle (2014-08-25) https://www.bgcarlisle.com/blog/2014/08/25/proof-of-prespecified-endpoints-in-medical-research-with-the-bitcoin-blockchain/

550. The most interesting case of scientific irreproducibility? Daniel Himmelstein Satoshi Village (2017-03-08) http://blog.dhimmel.com/irreproducible-timestamps/

551. OpenTimestamps: a timestamping proof standard(2017-05-16) https://opentimestamps.org/

552. greenelab/deep-review GitHub (2017) https://github.com/greenelab/deep-review

